# Is it selfish to be filamentous in biofilms? Individual-based modeling links microbial growth strategies with morphology using the new and modular iDynoMiCS 2.0

**DOI:** 10.1101/2023.06.27.546816

**Authors:** Bastiaan J R Cockx, Tim Foster, Robert J Clegg, Kieran Alden, Sankalp Arya, Dov J Stekel, Barth F Smets, Jan-Ulrich Kreft

**Affiliations:** Department of Environmental and Resource Engineering, Technical University of Demark, Bygningstorvet, Bygning 115, 2800 Kgs. Lyngby, Denmark; Centre for Computational Biology & Institute of Microbiology and Infection & School of Biosciences, University of Birmingham, Edgbaston, Birmingham, B15 2TT, UK; School of Biosciences, University of Nottingham, Sutton Bonington Campus, Loughborough, Leicestershire, LE12 5RD, UK

## Abstract

Microbial communities are found in all habitable environments and often occur in assemblages with self-organized spatial structures developing over time. This complexity can only be understood, predicted, and managed by combining experiments with mathematical modeling. Individual-based models are particularly suited if individual heterogeneity, local interactions, and adaptive behavior are of interest. Here we present the completely overhauled software platform, the <u>i</u>ndividual-based <u>Dyn</u>amics <u>o</u>f <u>Mi</u>crobial <u>C</u>ommunities <u>S</u>imulator, iDynoMiCS 2.0, which enables researchers to specify a range of different models without having to program. Key new features and improvements are: (1) Substantially enhanced ease of use (graphical user interface, editor for model specification, unit conversions, data analysis and visualization and more). (2) Increased performance and scalability enabling simulations of up to 10 million agents in 3D biofilms. (3) Kinetics can be specified with any arithmetic function. (4) Agent properties can be assembled from orthogonal modules for pick and mix flexibility. (5) Force-based mechanical interaction framework enabling attractive forces and non-spherical agent morphologies as an alternative to the shoving algorithm. The new iDynoMiCS 2.0 has undergone intensive testing, from unit tests to a suite of increasingly complex numerical tests and the standard Benchmark 3 based on nitrifying biofilms. A second test case was based on the “biofilms promote altruism” study previously implemented in BacSim because competition outcomes are highly sensitive to the developing spatial structures due to positive feedback between cooperative individuals. We extended this case study by adding morphology to find that (i) filamentous bacteria outcompete spherical bacteria regardless of growth strategy and (ii) non-cooperating filaments outcompete cooperating filaments because filaments can escape the stronger competition between themselves. In conclusion, the new substantially improved iDynoMiCS 2.0 joins a growing number of platforms for individual-based modeling of microbial communities with specific advantages and disadvantages that we discuss, giving users a wider choice.

**Author summary:** Microbes are fascinating in their own right and play a tremendously important role in ecosystems. They often form complex, self-organized communities with spatial heterogeneity that is changing over time. Such complexity is challenging to understand and manage without the help of mathematical models. Individual-based models are one type of mathematical model that is particularly suited if differences between individual microbes, local interactions and adaptive behavior are important. We have developed a completely overhauled version of iDynoMiCS, a software that allows users to develop, run and analyze a wide range of individual-based models without having to program the software themselves. There are several capability enhancements and numerous small improvements, for example the ability to model different shapes of cells combined with physically realistic mechanical interactions between neighboring cells. We showcase this by simulating the competition between filaments, long chains of cells, with single cells and find that filaments outcompete single cells as they can spread quickly to new territory with higher levels of resources. Users now have a wider choice of platforms so we provide guidance on which platform might be most suitable for a given purpose.

## Introduction

Microbes are found everywhere on Earth where conditions are suitable for life, often as microbial communities in self-organized assemblages such as biofilms [1]. They have a long evolutionary history through which they diversified into a huge number of species with fascinating characteristics and behaviors. Microbes master metabolism and thus enable biogeochemical cycles. Yet the complexity arising from the high diversity of their communities undergoing spatio-temporal dynamics makes it challenging to understand, predict and manage these communities [2]. This challenge can be best met by an integration of *in situ* observations, experiments in mesocosms and laboratory models and mathematical modeling [2].

Microbes growing in biofilms are a good example. Due to metabolic transformations of resources diffusing into the self-assembling biofilms, the aggregates become spatially structured including metabolite and resulting physiological gradients while growth leads to clonal populations. These changing environmental conditions prompt differences in gene expression, phenotype and behavior compared to planktonic cells [3]. For example, biofilm-dwelling *Pseudomonas aeruginosa* up-regulate production of extracellular polysaccharides (EPS), while *Staphylococcus aureus* biofilms up-regulate enzymes involved in glycolysis and fermentation [4]. Even in single species populations, phenotypic heterogeneity can become substantial [5,6]. Coupled to the local environmental changes, biofilm microbes experience selective pressures different to planktonic microbes. These are just a few points, but they already demonstrate the challenge of complexity in biofilm communities.

Biofilms are also a good example of insights derived from mathematical modeling, going back to the 1970s [7]. Early models treated the biofilm as a continuum in one dimension (1D), which enabled insights into substrate consumption driving the formation of gradients and diffusional fluxes and vertical stratification [8–10]. A key insight from later 2D and 3D models enabling the emergence of complex spatial structures was that the physics of mass transfer is sufficient to understand the formation of finger-like biofilm structures, arising from a positive feedback in growth where the cells at the surface of the biofilm with best access to substrate grow best so that their offspring are even closer to the substrate source and grow even better [11–13]. The detailed reconstruction of early biofilm growth observed through advanced microscopy coupled with detailed individual-based modeling of mechanical cell-cell interactions demonstrated that mechanics alone is sufficient to understand and predict early biofilm formation in *Vibrio cholerae* (before substrate gradients cause growth limitations [14]).

In such individual-based models (IbMs), microbial cells are modeled as agents, partially autonomous physical entities with individual properties and behavior [15]. This enables understanding of how these individuals affect other individuals in the community and the environment and are affected by the other individuals and the environment in turn. Properties of the community such as spatial patterns, fitness, productivity and resilience emerge from the behavior of the individuals in that community. IbMs are thus particularly suited to capture the effects of local interactions, individuality and adaptative behavior on spatio-temporal dynamics. This includes stochastic events such as dispersal and community assembly, up-regulation of genes, mutations or horizontal gene transfer. For example, Hellweger [16] used an IbM to model the gene expression and differentiation of *Anabaena* spp. individuals within a filament, and were able to reproduce almost all of the patterns observed *in vitro*.

To facilitate the use of individual-based modeling of microbial communities for scientists with little experience of programming, the open source modeling platform iDynoMiCS (individual-based <u>Dyn</u>amics <u>o</u>f <u>Mi</u>crobial <u>C</u>ommunities <u>S</u>imulator) was introduced [17], which we now refer to as iDynoMiCS 1. It was the result of a collaborative effort merging features of previous models into a common basis for further development. iDynoMiCS has been facilitating a range of studies and influenced the design of other modeling platforms [18–23]. Here we present, after a long phase of development and testing, a completely overhauled new version, simply called iDynoMiCS 2.0. We came closer to the original aim of making iDynoMiCS easy to use for non-programmers while substantially enhancing its capabilities and computational efficiency. Key new features and improvements are: (1) Enhanced ease of use right from the start, from using a simple guided java installer, lack of dependence on other software installations, a graphical user interface (GUI) for running simulations, editing model specification (protocol) files, unit conversions, data analysis and visualization of live results or re-loaded past results, a collection of model examples and online wiki for guidance, autonomous adjustments for solver convergence. (2) Increased performance and scalability enabling simulations of up to 10 million agents in 3D biofilms. (3) Kinetics of chemical or agent-catalyzed reactions can now be specified with any user-chosen arithmetic function. Local or intracellular conditions can be incorporated in these expressions enabling adaptive behavior such as metabolic switching or change in kinetics due to mutations. (4) Agent properties can now be assembled from orthogonal modules giving the user pick and mix flexibility. The same is true for processes. Thus, the complexity of agents or processes can be adjusted to fit the modeling purpose. Due to the modular structure, it has become easier to implement novel functionality. (5) Force-based mechanical interaction framework enabling attractive forces and non-spherical agent morphologies, which was impossible with the shoving algorithm. We showcase this new feature in a case study demonstrating how the fitness of filaments benefits from escaping competition.

## Model development and description

The description of iDynoMiCS 2.0 and the case studies presented in this paper follow the ODD (Overview, Design concepts, Details) protocol for describing individual- and agent-based models [24,25]. However, iDynoMiCS 2.0 is not one specific model but a platform to facilitate the specification of a broad variety of models. Hence, this section aims to provide a general description of the platform but cannot cover all possible models that could be simulated. Thus, we also provide detailed model-specific ODD descriptions together with the presented case studies.

### Overview

The purpose of iDynoMiCS 2.0 is to facilitate the simulation of large groups of individual microbes and their interactions in a microbial population or community, either in a well-mixed chemostat-like or a spatially structured biofilm-like compartment. The aim is to study and predict how the interactions and properties of individual microbes lead to emergent properties and behaviors of microbial communities.

### Entities, state variables and scales

Entities, state variables and scales may vary from one model implementation to another. In a typical implementation, microbial cells are the principal agents. They can mediate both chemical and physical activities. Agents can have any number of properties and behaviors. Typical properties are position, mass, density, shape, composition and metabolic reactions. Typical behaviors are cell growth, division, death, extracellular polymeric substance (EPS) production and excretion. iDynoMiCS 2.0 refers to properties and behaviors of an agent as the “aspects” of an agent. Shared aspects can be set-up as a module and reused for all agents sharing these aspects. A typical agent is one or multiple orders of magnitudes smaller than the computational domain.

The simulated space (computational domain) in which agents reside is called a compartment. There are two types: spatially explicit compartments in 2D or 3D to simulate microbial assemblages such as biofilms and well-mixed compartments used to simulate the dynamics of a bulk liquid without agents or a planktonic community with or without inflows and outflows (batch, fed-batch or chemostat); these compartments are assumed to have no spatial structure and thus the concentrations of chemical species and agents are homogeneous.

Well-mixed and spatially explicit compartments may be connected to simulate how bulk and biofilm dynamics are coupled. Compartments can have multiple properties including: boundaries, physical dimensions, volume and a scaling factor. This scaling factor is used to translate between the size of a simulated representative volume element and the actual size of the system, it allows a smaller simulated compartment to represent a larger entity. There are no hard restrictions on compartment size, however, compartments typically have lengths up to a millimeter. Agents can only reside in a single compartment at any one time, but they can be transferred or move between spatially explicit and well-mixed compartments.

Dissolved chemical species in iDynoMiCS 2.0 are referred to as solutes. Compartments can contain any number of solutes. In well-mixed compartments, a solute is represented as a single concentration. In spatially explicit compartments, solutes are represented as 2D or 3D concentration fields in a Cartesian grid. The distance between grid nodes is referred to as the resolution, typically one to a few micrometers.

### Framework structure

In iDynoMiCS 2.0, models are structured, specified and instantiated hierarchically. The basic structure of a typical scenario is presented in Fig 1. The “Simulator” is the root of any model implementation. It loads the model specification from a protocol file provided by the user either through the GUI or the command terminal. General software control parameters are managed at the simulator level, as well as the scheduling of sub-models, the management of compartments and the storage of species modules (reusable sets of aspects). The simulator steps through its compartments and saves the model state at each global time-step. A compartment stores its shape and size, solutes and agents in the compartment and processes occurring within the compartment. Agents and processes store their properties as aspects. This layered structure of model input provides a level of modularity to the iDynoMiCS 2.0 software and model implementations.

**Fig 1.**
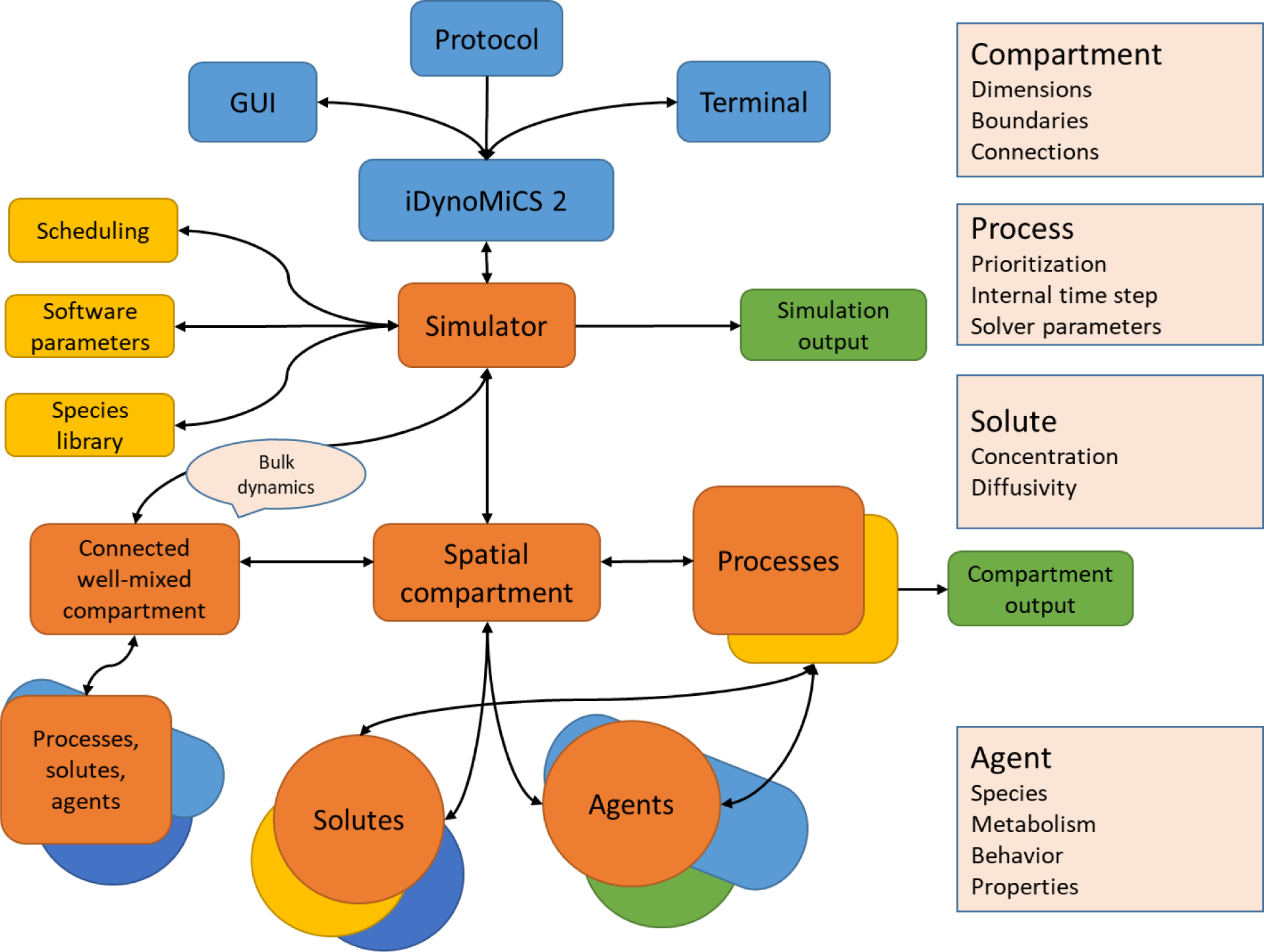
The basic structure of an iDynoMiCS 2.0 model. Interaction with the program takes place through the GUI or command line terminal. A protocol file specifying a model can be loaded to initialize the simulator. If parameters are missing from the protocol file, a default if loaded or the user is queried if no default exists. Scheduling ensures predictable handling of the compartments and the order of processes occurring within them. A species library is kept such that properties and/or behavior that are identical for agents of the same species can be looked up from the library. The simulator further ensures that the model state is saved at the end of each global time step. Spatially explicit and well-mixed compartments can be connected. Solutes concentration fields are stored as matrices, which include local solute concentrations, local diffusivity and reaction rates. The collective of agents represents the biofilm, agents may have many properties depending on user specifications, basic properties are species, mass and position of the agent. Processes act upon the information in the model system and describe the processes occurring in the model such as mechanical interactions or diffusion, or generate output from the active model state.

Modularity enables flexible combination and facilitates software maintenance and development. An iDynoMiCS 2.0 model is formulated as distinct modules describing specific parts of the model such as a compartment, process or an agent. An example model description is included in Box 2. These modules can be referenced to add the same object, property or process in another compartment or agent. For example, multiple agents of different species can implement the same module describing cell shape but implement a different module describing metabolism.

Within the software, modularity is implemented through the use of software interfaces and abstractions. These software interfaces ensure common functionality such that loading or storing of data, scheduling and initiation are handled in the same way for any software class implementing these interfaces. This concept makes it easier to add new features and extensions to iDynoMiCS 2.0, since an extension can be integrated into the framework without additional work to handle initiation, data handling, scheduling and other auxiliary functionality (Box S1).

Support for arithmetic and basic logic expressions provides flexibility to iDynoMiCS 2.0 models. Users can simulate any type of kinetic or physical interaction model that can be described by a standard arithmetic expression. Logic expressions are particularly useful in models with biological switches or thresholds. Typical examples are metabolic or morphological switches. Since any kind and number of aspects can be changed, the characteristics of an agent can completely change at runtime. Logic expressions can also be used to filter agents matching specific criteria, which can be useful for further analysis, or to color agents based on their properties. Logic expressions may incorporate arithmetic expressions.

### Process overview and scheduling

Many processes can take place in a microbial community, often simultaneously. To capture these processes in a computer model, they are represented by simplified mathematical models, discretized and handled in an asynchronous fashion. An iDynoMiCS 2.0 simulation steps through a number of global time-steps. Within each global time-step and for each compartment, sub-models describing activities occurring in the microbial community are executed. Processes included in a model depend on the purpose and design of the model. However, most simulations include: physical interactions, (bio-)chemical reactions, diffusion of chemical species, microbial growth and cell division. A detailed overview of implemented submodels follows later.

During one global time step, a list of processes, defined and scheduled by the user, are executed. Each process is assigned a time step size and a priority by the user, if not, the global time step and order of occurrence in the protocol file are adopted as time step and priority. A process can be repeated during a single global time step and may have smaller internal time steps. Processes are handled in a specific order; the process time step is used to determine what process is due first. If multiple processes are due simultaneously, the process priority is used to determine the order. Box 1 shows a scheduling example as implemented in the biofilms promote altruism case study presented later.

#### Box 1. Process schedule for the biofilms promote altruism case study within a single global time step.

1. (Bio-)chemical reactions and diffusion

a. Update solute grids and agent masses (e.g. (bio-)chemical conversion)
b. Update solute boundaries
2. Agent updates

a. Cell division
b. Differentiate (switch between sphere or rod shaped if applicable)
c. Update cell size
3. Mechanical relaxation

a. Update cell positions and mechanical stresses until relaxation criteria are met
4. Compartment reporting

a. Density grid (csv)
b. Compartment summary (csv)
c. Graphical output (pov/svg)
5. Save simulation state (xml/exi)

### Design concepts

#### Basic principles

Microbes are modeled as individual particles that interact with their environment. Mass is conserved and thus the mass balance of any ‘element’ in the model system should be closed.

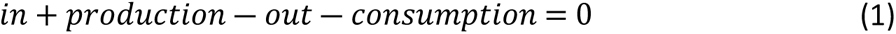

#### Emergence

System-level phenomena such as the distribution of microbial agents, the spatial structure of a biofilm, and chemical conditions all emerge from interactions between agents and their local environment. This local environment can consist of other agents, local chemical concentrations or a local surface.

#### Adaptation

iDynoMiCS 2.0 models can include adaptation. By sensing the local environment or internal states, an agent may change its characteristic using the differentiate aspect. Adaptation can also be stochastic, where heritable stochastic changes in an agent’s characteristics can lead to selection of the lineage best suited to survive or thrive in the simulated environment.

#### Objectives

Agents can have objectives. For example, agents may have the objective to move to a region with higher substrate availability (chemotaxis).

#### Learning and prediction

Although microbes have no brain, they can have memory and process signals. For example, chemotaxis requires memory. Also, event occurrences may be stored as an aspect of the agent and influence the behavior or characteristics of an agent.

#### Sensing

Locally sensed nutrient concentrations can affect growth rate or adaptation. Neighboring cells may affect the spatial direction of cell division in filamentous agents or affect physical displacement. A combination of the above supports microbial behaviors such as dormancy or chemotaxis.

#### Interaction

Agents interact with their local environment by consuming and producing solutes, including extracellular enzymes, or particles such as EPS. As a consequence, they indirectly interact with neighboring agents, leading to competition, cooperation or communication. The agents further interact with their physical environment by pushing or pulling other physical entities, including other agents as well as physical surfaces in the computational domain.

#### Stochasticity

Model implementations can include stochastic processes, examples are allocation of biomass to daughter cells upon cell division, placement of daughter cells relative to the position of the mother cell, stochastic movement (for example Levi flight), and mutations or other random perturbations. A random seed is saved so simulations can be continued with consecutive random numbers or repeated with an identical series of random numbers if desired.

#### Collectives

Microbes may aggregate actively through motility coupled with communication and the expression of surface proteins or passively through cell division and the production of an extracellular matrix [26]. Consequently, agents can form a local collective of unrelated or related cells. Moreover, communication can coordinate collective action that does not require aggregation. iDynoMiCS 2.0 has the basic building blocks to model coordinated collective behavior such as quorum sensing.

#### Observation

The model state is saved at the end of each global time-step. Additional compartment output can be saved, including physical and biological states of all agents and the chemical state, at any given time and location in the simulation.

### Initialization

The initial state of a simulation is highly flexible and should be provided by the user. This includes solute concentrations as well as positions and states of agents, which may be based on experimental data or generated to investigate hypothetical scenarios. iDynoMiCS 2.0 includes several helper classes to generate initial states. A ‘random spawner’ can be used to randomly distribute large numbers of agents of a certain type over a pre-defined region, applied in the initialization of the stress test and Benchmark 3 case study. A ‘distributed spawner’ has a similar function but distributes agents regularly at a pre-defined interval, applied in the initialization of the biofilms promote altruism case study. It is also possible to manually define an initial state for any individual or to utilize a previous simulation outcome as an initial state for a new simulation, for example to implement a perturbation.

### Input

All simulations are initiated using iDynoMiCS 2.0 protocol files. They follow the logical structure of the model setup and are structured using the Extensible Markup Language (XML). Both XML and EXI (Efficient XML Interchange) files can be used. It is recommended to include units when specifying parameters in the protocol file, units are converted to iDynoMiCS’s base unit system, which avoids unit conversion mistakes.

#### Box 2. An abbreviated protocol file showing the hierarchical structure.

Setting up a basic protocol is relatively simple and supported by the GUI. In this example, 30 copies of an agent of species “bacterium” are added to a rectangular domain. Bacterium is defined by the reusable “coccoid” aspect highlighted in green as well as a growth reaction which is only used for this species. The coccoid module describes basic physical properties and behavior of a generalized coccoid including agent density, division mass, etc. Bacterium contains the “reactions” aspect which is a list of all reactions the agent can catalyze (in this case only the crucial growth reaction). The reaction node includes an arithmetic expression defining the growth kinetics based on the local solute concentrations of two solutes (carbon and oxygen) and associated kinetic constants and stoichiometry. The protocol further describes the properties of the compartment, the solutes and processes to be loaded. The full protocol file is just 60 lines and included in S1.7.

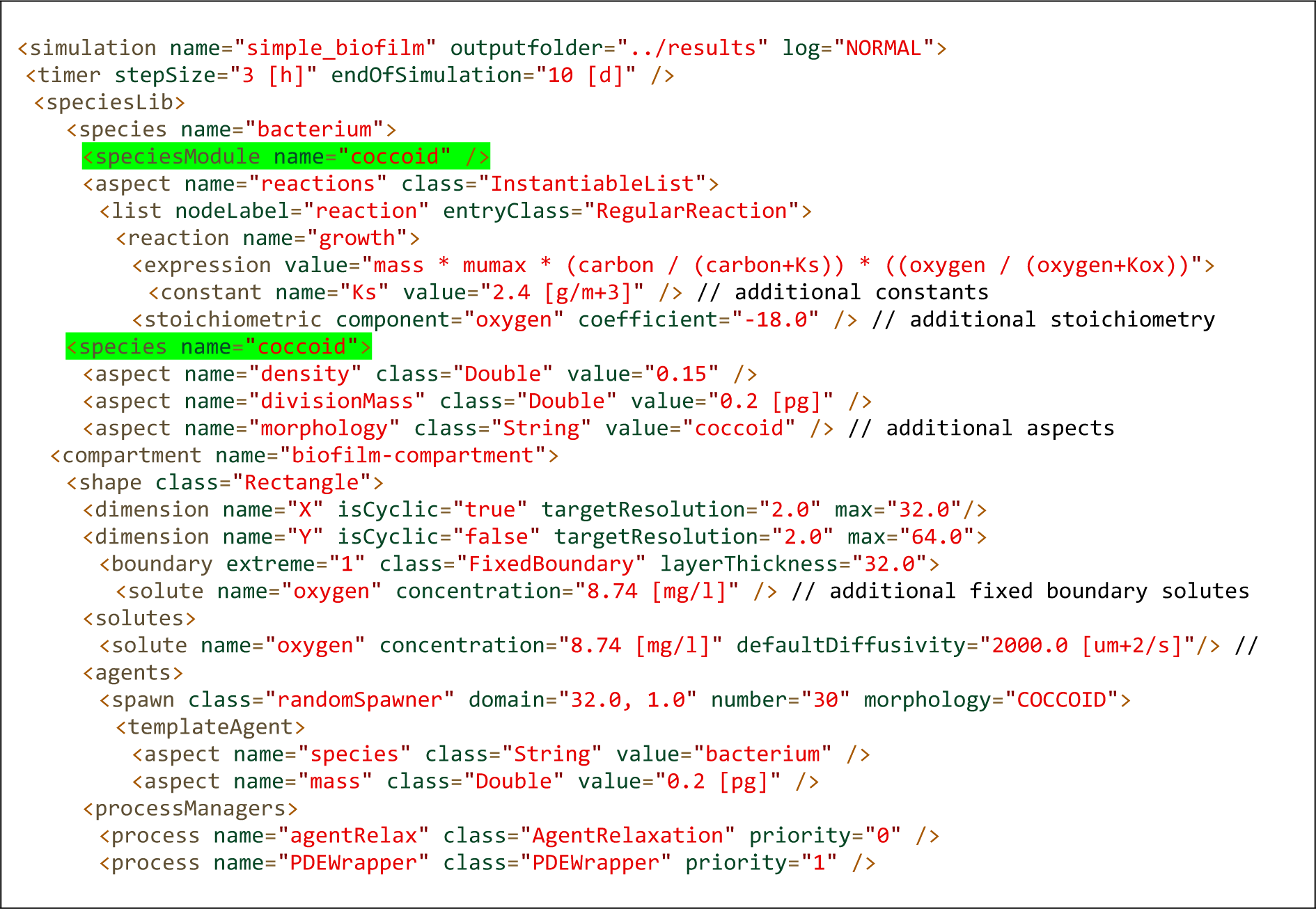

### Output

The model state is saved as XML or EXI file to reduce file size after each global time step and follows the same structure as a protocol file. The output also includes all information required to restart the simulation. It is also possible to save additional output per compartment to facilitate later analysis or visualization. Visual output includes SVG or POV formats to render the agents in a compartment. The hue, saturation or brightness of the agent can be adjusted based on its properties to convey additional information about the agent’s state. A CSV file listing agents with key properties of interest can also be produced, this list can be filtered to only include agents matching specific criteria. Further, the density of agents in the compartment can be reported for all agents or those matching specific filters (such as belonging to a certain species). Finally, a summary with key statistics on the compartment can be written such as mean solute concentrations, total agent counts and masses, agents matching specific criteria, etc. The summary is useful to quickly plot time series data.

### User interface

Simulations can be loaded and started through the command line or a GUI (Fig S9), which may be used to review, edit or create protocol files before running them. During the simulation, the GUI provides key information on the simulation state (such as substrate concentrations, species abundance, convergence of the reaction diffusion solver, etc.). The spatial domain can be rendered directly to display agent distributions and concentration gradients. Lastly, the GUI can be used to extract key data from iDynoMiCS 2.0 output files, convert between EXI and XML files and convert numbers between different unit systems including SI and iDynoMiCS 2.0 base units.

### Submodels

Here we describe key submodels currently implemented in the iDynoMiCS 2.0 platform. Specific model implementations may utilize a subset of these submodels or alternative submodels that extend capabilities further.

#### Physical representation of agents

Microbial cells in the model have position and extent in 3D continuous space, constructed from points and swept-sphere volumes (Fig 2). Specifically, spherical cells dubbed “cocci” are constructed from a single point with a spherical volume, rod-shaped cells dubbed “bacilli” are constructed from two points connected by a line-segment and a swept-sphere volume along the line-segment, filaments are represented as a chain of either one or both. Every point has a mass associated with it.

**Fig 2.**
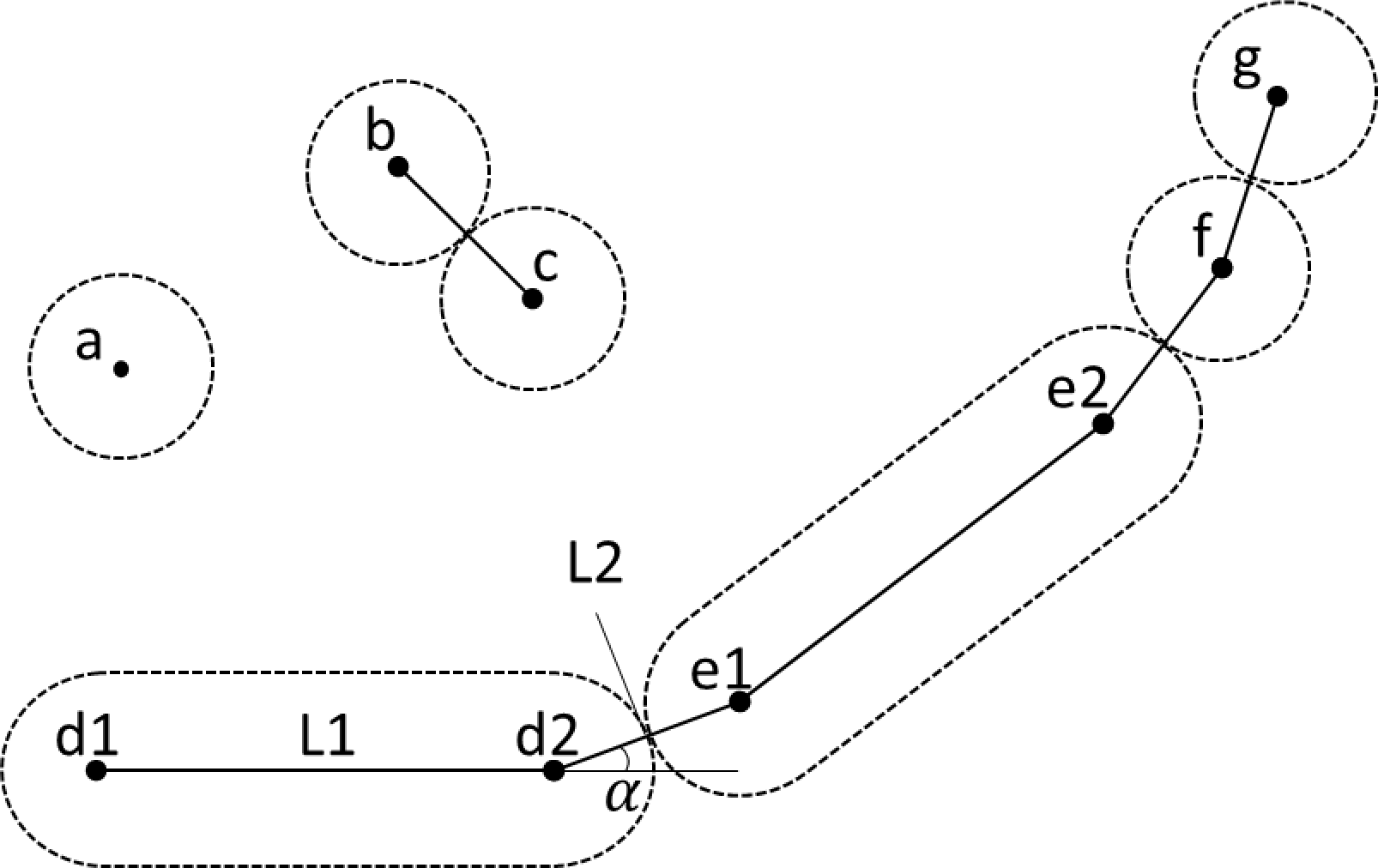
Different agent shapes in iDynoMiCS 2.0. Dashed lines indicate sphere-swept volumes of ‘dots’ or line-segments. Dots are mass-points indicating position and orientation of agents. Solid lines indicate mechanical interactions between points (forces between points modeled as springs): Collision interaction (b-c), spine interaction responsible for the rigidity of rod-shaped agents (d1-2 and e1-2), connecting interactions (d2-e1, e2-f, f-g). α is the angle between two elements of a filament. This angle can be counteracted by a torsion spring applying forces on d1, d2 and e1. L1 and L2 are the moment arms. The torsion spring applies force until the angle α reaches 180°, aligning the three points.

In order to simulate 2D models, a number of assumptions and adjustments have to be made. An implicit third dimension (z) is required to retain consistency for physical units such as volume or concentration, in iDynoMiCS 2.0 this third dimension is 1 µm thin. The 2D agent shapes are extruded into this virtual dimension, thus their pseudo 3D shapes have a uniform cross-sectional area and a thickness of 1 µm. This translation from 3D to 2D comes with several side effects such as a lower density of circle packing as compared to sphere packing [27]. To mitigate this effect, Clegg et al. [27] proposed a density scaling factor of 0.82 for 2D simulations with spherical agents. Appropriate scaling factors for other agent shapes or mixtures of agent shapes are unknown.

Another side effect of 2D simulations results from the constraint that the length of the virtual third dimension has to be identical for all agents. This can result in unwanted agent size effects for agents with very small or large radii. iDynoMiCS 2.0 can scale the density of agents in order to retain consistent agent diameters and lengths between 3D and 2D simulations. This method is described in detail in S1.12. This method is not a perfect solution as it will introduce new side effects, the local shifts in biomass concentrations will increase nutrient competition for large agents whilst decreasing for small agents. If there are large differences in agent size, it is recommended to test validity of 2D simulations with 3D simulations.

Additionally, biofilm structure and development can be affected in 2D simulations. Vertically stratified biofilms with chemical gradients from substratum surface to the biofilm-liquid interface dominating, or biofilms with gradients in the second dimension, parallel to the substratum surface, that are equivalent in the third dimension which is also parallel, can be accurately modeled in 2D. However, biofilms that form finger-like or other superstructures become artificially substrate limited due to the lack of mass transport in the third dimension. Consequently, these superstructures will form under different environmental conditions in 2D versus 3D.

#### Mechanical interaction framework

In the original iDynoMiCS 1, agent overlap was resolved with a shoving algorithm. In iDynoMiCS 2.0, mechanical interactions between agents (or with surfaces in their environment) as well as between different points within the same agent is based on forces. This Force-based Mechanics (FbM) framework builds on the mass-spring models of Janulevicius *et al.* [28], Celler *et al.* [29] and Storck *et al.* [30] but is no longer limited to spring forces. The shoving algorithm is still available as an alternative or for comparison with iDynoMiCS 1.

Before an interaction force can be calculated, the interaction needs to be detected. Direct links between two points of the same agent are stored as an aspect of the agent. Additionally, neighboring entities are found efficiently through a search of the quad- or octree that keeps agents spatially sorted. Through collision detection, as described and implemented by [31] and [30], physical interaction between neighboring agents is tested. The distance between two objects (which can be negative) is used in a force model to calculate the resulting force of the interaction. The quad-, octree and collision detection algorithms were modified to work with periodic boundaries.

#### Forces

The FbM solver exploits the fact that under conditions of very low Reynolds numbers, inertial forces on a particle become negligible [32,33]. Because of this, the sum of all forces applied to the mass-point (the net force ∑ *F*_*p*_) can be assumed to balance the mass-point’s drag force *F*_*D*_ (*F*_*D*_ = ∑ *F*_*p*_) near instantly. By applying Stokes’ law, the terminal velocity of the mass-point *v*_*t*_can be obtained, under low Reynolds number conditions the mass-point reaches this velocity near instantly and thus we can assume *v* ≈ *v*_*t*_ when Re << 1 (Eq 2) [34]. This simplification effectively halves the number of ordinary differential equations that need to be solved.

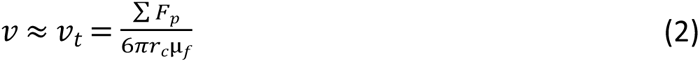

With *r*_*c*_ the diameter of the cell and *μ*_*f*_ are the dynamic viscosity of the fluid.

The mass-point’s velocity, and by extension the point displacement, is resolved using a forward Euler method or the second-order Heun’s method [35].

Multiple types of interactions may lead to a net force being exerted on any given point. These forces may be due to collision, close proximity repulsion or attraction, attachment and internal structure. A force model for these interactions can be provided through the protocol file. Default models are used if no model is provided. The default force model for agent collision is the Hertz soft sphere model [36] (Eq 3), where r_eff_ is the effective radius, E_eff_ the effective Young’s modulus and ξ the agent overlap.

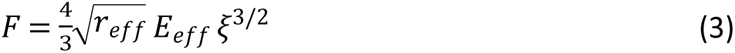

Rod-shaped cells and filaments have permanent connections between points (L1 & L2 in Fig 2). By default, these connections are modeled as linear springs following Hooke’s law (Eq 4), where k is the spring constant and *δl* the difference between the current length and the rest length.

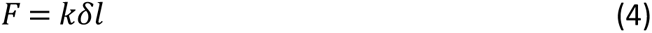

An internal force can also be specified to resist bending, for example for segments of a filament (Angle α in Fig 2). The default model for such torsion springs is the angular form of Hooke’s law (Eq 5), where κ is the spring constant, *δ*ϴ the difference between the actual angle and rest angle, L the length of the momentum arm.

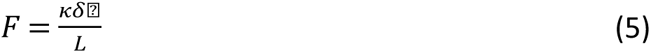

Forces for agents in proximity can also be specified, for example, close range repulsion and/or attraction, but are not applied by default.

#### Reactions

A reaction transforms chemicals; these chemical species may be modeled explicitly as solutes or types of the structured biomass within cells (e.g., regular biomass and storage compounds), or implicitly and so left out of the model description (e.g., water in aqueous environments). Reactions have a rate and stoichiometry. They may be catalyzed by agents or occur independently of agents in the environment.

Reaction rates may depend on the concentration of certain reactants, such as solutes or biomass, and other variables, such as temperature. Within iDynoMiCS 2.0, rate equations can be expressed through any type of arithmetic expression, which allows the use of almost any kinetic model, not only Monod type kinetics but also thermodynamics-based kinetics such as the models by Rittmann and McCarty [37] or Heijnen [38]. Such a thermodynamics-based IbM approach was previously implemented by Gogulancea et al. [39]

Positive stoichiometry signifies production, and negative stoichiometry consumption, when the reaction proceeds in the forward direction (has a positive rate). Thus, the production rate of a solute, *i*, by reaction, *j*, is given by:

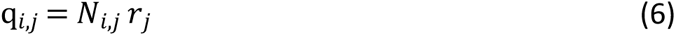

where *N* denotes stoichiometry and *r* denotes reaction rate.

#### Solutes in well-mixed environments

In compartments assumed to be well-mixed, there is no spatial structure for solutes nor for agents., thus the rate of change is an ordinary differential equation summing inputs, outputs and reactions

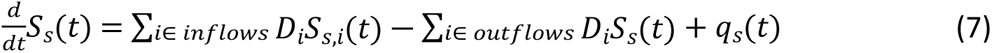

where *S_s_(t)* is the concentration of solute s at time *t* and *D_i_* the dilution rate for a given inflow/outflow. The production rate expression *q_s_(t)* combines environmental reactions, *q_s,env_*, with reactions catalyzed by each agent, *q_s,agent_*.

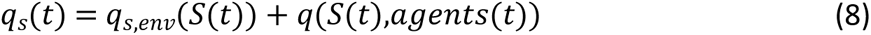

where *S(t)* is the local solute concentration.

#### Diffusion-Reaction of solutes

Within spatially explicit compartments, the dynamics of solutes are governed by two processes: Fickian diffusion and reactions.

For each solute, the rate of change for each solute is now given by the elliptic Partial Differential Equation (PDE) that combines Fickian diffusion, given by the divergence div of the diffusional flux driven by the concentration gradient ∇*S*_*s*_, and reaction [40].

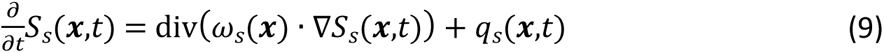

where ***x*** is the spatial position, ω*_s_* is the local diffusivity*, and S_s_(**x**,t)* the local concentration of solute *s*. Dynamics at compartment edges are governed by boundary conditions.

As in iDynoMiCS 1 [17] and other biofilm models, a pseudo steady-state assumption is made to model solute dynamics because reaction and diffusion processes are several orders of magnitude faster than biomass growth, decay and detachment processes [41]. Hence, reaction and diffusion rates rapidly reach a pseudo steady-state while the biomass distribution is changing so slowly that it can be considered ‘frozen’ [42]. This time scale separation drastically reduces the computational demand of the simulation but it should be checked whether the pseudo steady-state assumption is reasonable.

#### Boundaries

Spatially explicit compartment boundaries can be of different types. (a) Solid boundaries where neither agents nor solutes can pass through (Neumann or no flux boundary condition). (b) Fixed concentration or Dirichlet boundary conditions where the solute concentrations are fixed to preset values or determined dynamically by an ODE such as Eq 7 for a connected well-mixed compartment. (c) Periodic boundaries where agent and solutes can move through the boundary to the opposite side of the domain, which ensures identical concentrations on each side to avoid edge effects.

Well-mixed compartments may also exchange solutes and agents with other well-mixed or spatially explicit compartments through its boundaries. (d) A boundary connecting a spatially explicit with a well-mixed compartment, where solute concentration gradients in the spatially explicit compartment result in a diffusive flux through the boundary determined by the concentration gradient at the boundary. (e) An inflow boundary with preset solute concentrations and flow rate. (f) An outflow boundary with the concentrations matching the content of the well-mixed compartment and a preset flow rate, which may match the inflow. The well-mixed compartment may change volume over time if inflow and outflow rates do not match. An outflow boundary can be set to retain the agents present in the well-mixed compartment to model biomass retention in a retentostat.

#### Plasmid Dynamics

Plasmid dynamics incorporates conjugative transfer and segregative loss of plasmids. Plasmids are included as aspects of an agent, with loss defined as an event occurring upon agent division. The dynamics of conjugation modeled here follow the known behavior of F-pili driven plasmid transfer described in S1.11.

## Results

We start by summarizing the strategy and results of our model verification efforts that were focused on comparing our numerical solvers against analytical solutions in the simpler test cases and well tested solvers implemented in other software in the more complex test cases. This is followed by using benchmarks. First the standard Benchmark 3 for comparison of biofilm models as previously done for iDynoMiCS 1. Then a second, new benchmark that is highly sensitive to initial conditions and spatial structures due to positive feedbacks where we can compare iDynoMiCS 2.0 results with BacSim and also investigate the often-neglected effect of different biofilm spreading mechanisms. Finally, we demonstrate some of the new capabilities of iDynoMiCS 2.0 – simulating filaments which requires FbM – and show some surprising new results.

### Model verification and performance

#### Component testing and solver verification

iDynoMiCS 2.0 has undergone a rigorous verification process. We focused on numerical solvers, writing code to inspect the state at each iteration and diagnose convergence of solvers. This process helped eliminate bugs and software inefficiencies. It also demonstrated solutions were numerically correct with deviations of <0.1% in all test scenarios with known analytical solutions. The testing process included single-component testing (or unit testing), multi-component testing, stress testing and benchmarking, following a strategy of increasingly complex test scenarios where analytical solutions were known for simpler cases, numerical solutions from well-tested other solvers for intermediate cases and comparison to output of a set of other models for the most complex test cases. Single-component testing entailed the testing of individual parts of the framework against known solutions or predicted convergence. This included tests for collision detection and collision response (S1.2), (bio-)chemical conversion and material transport in a well-mixed compartment (S1.3), microbial growth in a well-mixed compartment (S1.3) and (bio-)chemical conversion and diffusion in a spatially explicit compartment (S1.3).

Multi-component testing where multiple parts of the framework are tested simultaneously was performed through using various test scenarios. The goal of these tests was to see whether multiple components worked correctly in unison and whether iDynoMiCS 2.0 can perform more complex scenarios stably and reproducibly. Test scenarios are listed in S1.9, protocol files for all scenarios are available on the iDynoMiCS 2.0 GitHub repository.

#### Stress testing

Stress testing built on multi-component testing, but tested scalability of performance and pushed the limits of domain size. In the process, software limitations and bottlenecks were removed. The stress test verified that iDynoMiCS 2.0 was capable of simulating the development of a 3D biofilm over 175 simulated days to reach >10 million agents (Fig 3, detailed description in S1.4) in 11.34 days on a single processor.

**Fig 3.**
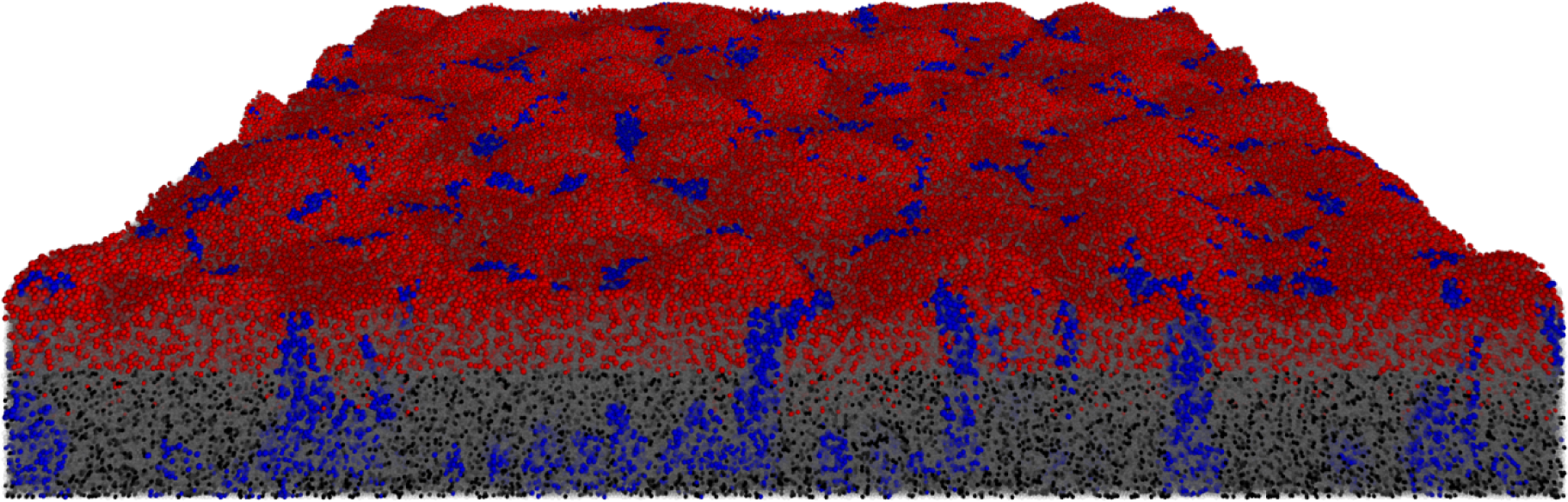
iDynoMiCS 2.0 was capable of simulating large 3D biofilms. A *nitrifying biofilm was initiated with 1,000 Ammonium Oxidizing Organisms (red) and 1,000 Nitrite Oxidizing Organisms (blue) in a 500×500×500 µm domain.* Both species produced EPS particles (gray semi-transparent). Agents that dropped below 20% of their division mass as a result of endogenous respiration (maintenance metabolism) became inactive (black). The 175-day biofilm contained 1.02×10^7^ agents (bacteria and EPS particles).

#### Benchmarking against various types of biofilm models

iDynoMiCS 2.0 was benchmarked against 1D or 2D continuum models, 2D Cellular Automata and 2D particle-based models including iDynoMiCS 1, using the multi-species nitrifying biofilm Benchmark Problem 3 or BM3 [43,44]. BM3 was the most complex benchmark developed by the International Water Association biofilm modeling task group to compare computational modeling approaches for biofilms and provide guidance for researchers. All models implemented the same processes. Unfortunately, some published BM3 results are limited to steady state concentrations of organic carbon (expressed as Chemical Oxygen Demand, COD) and ammonium in the bulk liquid that exchanges with the biofilm so we could only show that these results from iDynoMiCS 2.0 did not differ significantly from the distribution of results from the other models (Fig 4, Table SI9). iDynoMiCS 1 was previously shown to produce similar results to another particle-based model, NUFEB [21,45]. We also tested and confirmed that the physically realistic biomass spreading mechanism FbM in iDynoMiCS 2.0 gave similar results to the biomass spreading by shoving in iDynoMiCS 1, and for that purpose implemented shoving in iDynoMiCS 2.0 as well (Fig 4). Since FbM produces denser biofilms (in the absence of EPS particles that would distance cells explicitly), biofilm density had to be scaled accordingly for this comparison. An extensive BM3 description and results analysis is included in S1.5.

**Fig 4.**
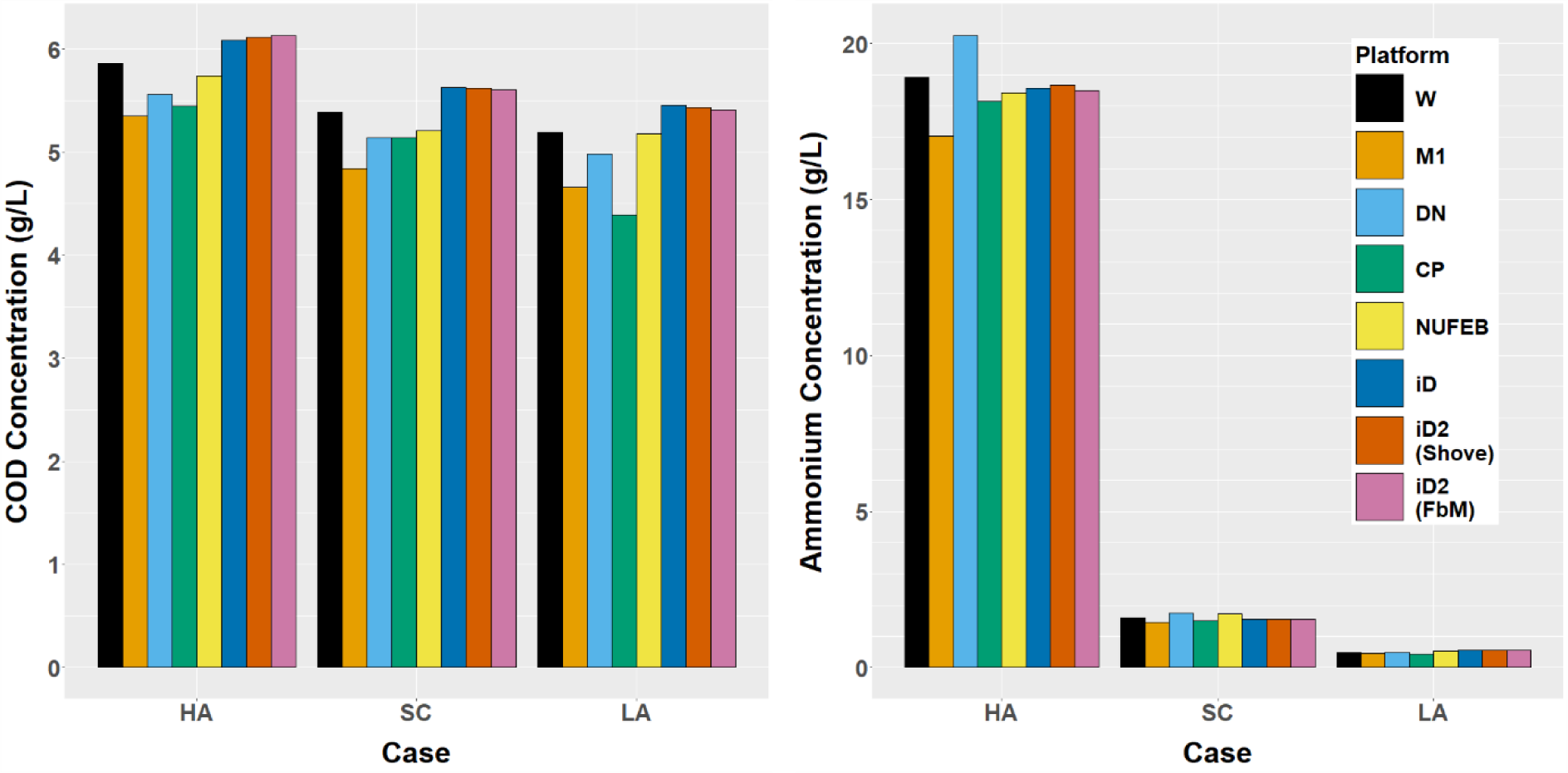
Comparing steady states in BM3. Steady state organic carbon (Chemical Oxygen Demand, COD) and ammonium concentrations in the bulk liquid for the three different BM3 cases (HA: High ammonium, SC: Standard case, LA: Low ammonium) across 7 model implementations (W: Wanner, M1: Morgenroth, DN: Dan Noguera, CP: Cristian Picioreanu, NUFEB: NUFEB, iD: iDynoMiCS 1, iD2: iDynoMiCS 2.0, either with shoving algorithm similar to iD or the new Force-based Mechanics).

### Comparing the effect of different biomass spreading mechanisms: Biofilms promote altruism case study

As a second and new benchmark for model testing and comparison, we have chosen a biofilm scenario where a positive feedback can amplify initially small differences leading to divergent results. This was thus a good opportunity to examine the effect of different biomass spreading mechanisms, comparing BacSim, the first implementation of shoving for biofilms [46] with iDynoMiCS 2.0. The scenario consisted of competing two growth strategies, a Rate Strategist (RS) that grew faster at every substrate concentration (higher *μ*_*max*_, same specific affinity/initial slope of the Monod kinetics) but had a lower growth yield, with a Yield Strategist (YS) that grew more slowly but had a higher growth yield and therefore converted the substrate diffusing into the biofilm with higher efficiency into biomass. This more economical use of resources is an example of altruistic behavior as it benefits selfish neighbors more than self [46]. Model parameters are in Table S11. We found that both IbMs generated the same qualitative outcomes, where YS won at lower initial density (Fig 5A and 5B), RS won at intermediate initial density (Fig 5E-H) and at even higher initial density, YS won again due to clusters of YS ending up on the biofilm surface by chance and then growing better as clusters of cells which are competing less with each other (Fig 5I-L). This combination of chance events – formation of a small cluster of YS cells at the biofilm surface – with positive feedback became obvious after longer simulations (Fig 5K and 5L).

**Fig 5.**
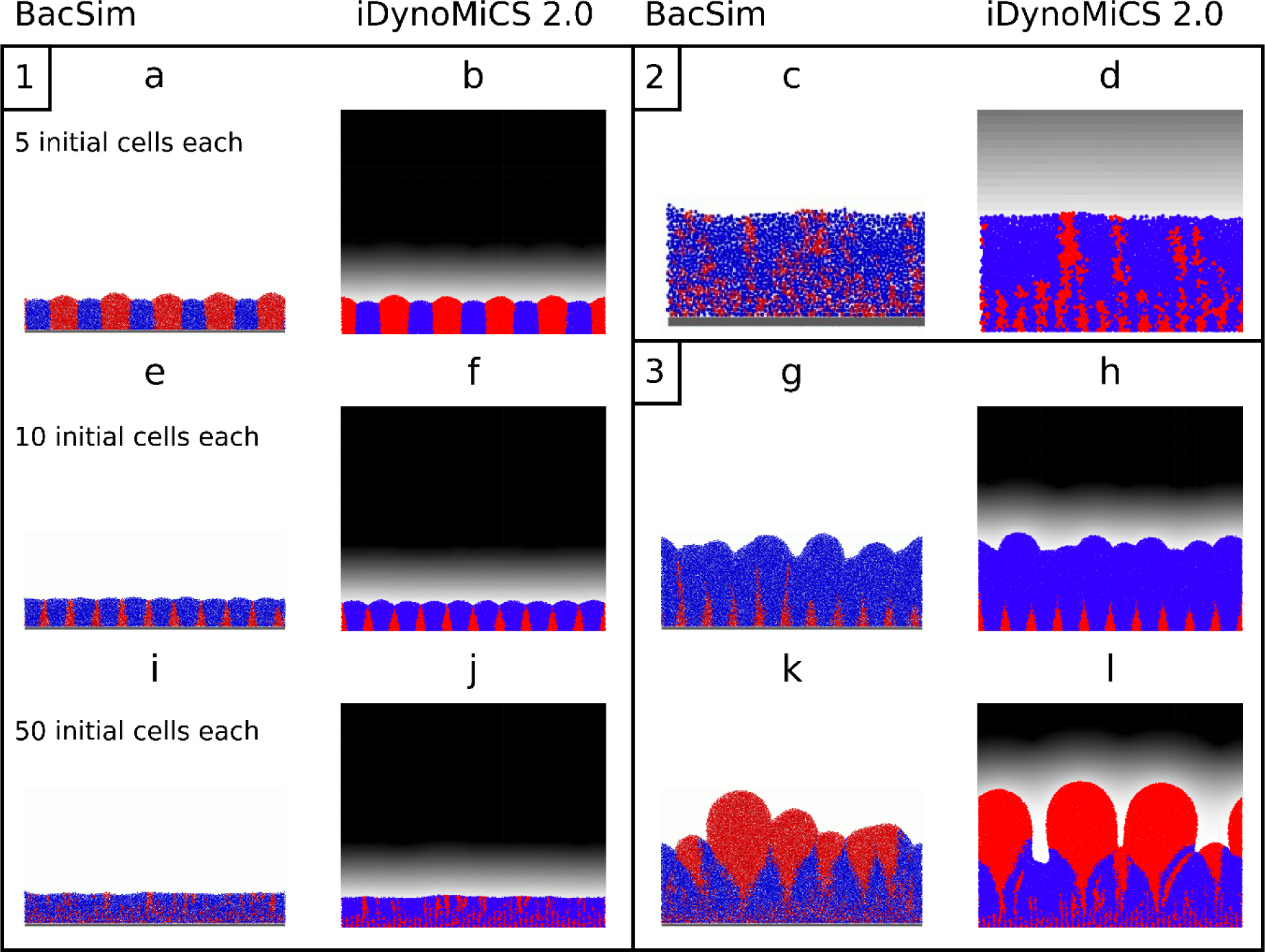
Biofilms promote altruism case study. Rate Strategist (RS, blue) and Yield Strategist (YS, red) competitions using the shoving algorithm in BacSim [46] (reproduced from “Kreft J-U (2004). Biofilms promote altruism. Microbiology 150: 2751–2760” with permission) were replicated in iDynoMiCS 2.0 with its force-based mechanics. Cells were initially placed in alternating, equidistant positions with increasing density from 5 cells per strategy (Scenario 1: a-b), 10 cells each (Scenario 2: e-h) to 50 cells each (Scenario 3: i-l and c-d). iDynoMiCS 2.0 panels show local oxygen concentration as a linear gray-level gradient from zero oxygen (0 mg/L, white) to a maximum concentration (S_ox_bulk_ = 1 mg/L, black). Box 1 shows 3-week-old biofilms. Box 2 zooms into panels i and j. Box 3 shows 10-week-old biofilms developed from the 3-week-old biofilms shown in the same position on the left.

While the qualitative outcomes were the same, the initial density thresholds separating regions where different strategies win were somewhat shifted. This was a result of the different biomass spreading algorithms: the shoving in BacSim led to more open spaces and increased mixing of cells locally, compared to force-based mechanical relaxation in iDynoMiCS 2.0, which in this case without EPS production only avoided overlap between agents but did not push cells further away (Fig S6). Note this resulted in increased overall biofilm density in iDynoMiCS 2.0. To compensate for this effect, we had to reduce agent density in iDynoMiCS 2.0 by 47% to maintain similar biofilm densities. Shoving can implicitly model the effect of EPS production generating space between agents, while EPS particles need to be included explicitly to generate space when using FbM simulations. iDynoMiCS 2.0 can do both.

### Filaments outcompeted cocci regardless of growth strategy and RS filaments expanded faster

Building on the biofilms promote altruism case study and the capability of iDynoMiCS 2.0 to simulate spherical, rod-shaped and filamentous microbes, we asked whether filamentous growth provides an additional advantage to Yield Strategists (YS). Since sufficiently large clusters of the cooperative YS cells outcompeted the Rate Strategists (RS) (Fig 5), we reasoned that growing as a filament, which can be considered to be a cluster of cells in one dimension, would give YS an additional advantage. Hence, we competed all combinations in a biofilm setting: coccoid RS vs. coccoid YS, filamentous RS vs coccoid YS, coccoid RS vs. filamentous YS and filamentous RS vs. filamentous YS. As filaments need a larger domain and freedom to bend and spread in all directions, these simulations required a 3D domain of sufficient size. The z-dimension was expanded to 12.5 µm, whilst keeping solute resolution at 1.5625 µm. Since the shoving algorithm cannot properly deal with filaments, the FbM of iDynoMiCS were required. It turned out that filaments were superior to cocci regardless of growth strategy – because filaments quickly gained access to the higher substrate concentrations at the top of the domain where the source of the substrate was located (Fig 6). This range expansion strategy of filaments is similar to cells producing EPS to rise quickly above biofilm neighbors towards the substrate source above, or trees growing faster to the top of the canopy to gain better access to sunlight [47], or the foraging strategy of cord-forming fungi that can form ‘bridges’ between discrete and sparsely scattered patches of nutrient resources [48]. Surprisingly, RS filaments won the competition against YS filaments in each case where the final outcome can be inferred (Fig 6 shows biofilm structures and Fig 7 shows population dynamics over time). A striking difference between Rate Strategist filaments and Yield Strategist filaments is the more open and less dense ‘forest’ structure produced by Rate Strategists. We suggest that the lower substrate consumption rate of the Yield Strategists allows their filaments to grow better in deeper regions of the biofilms than the Rate Strategist filaments, which gain a larger advantage when they happen to grow towards the top. Thus, the stronger competition (or self-inhibition) between Rate Strategist filaments favors expansion over density, leading to a ‘fluffier forest’ structure.

**Fig 6.**
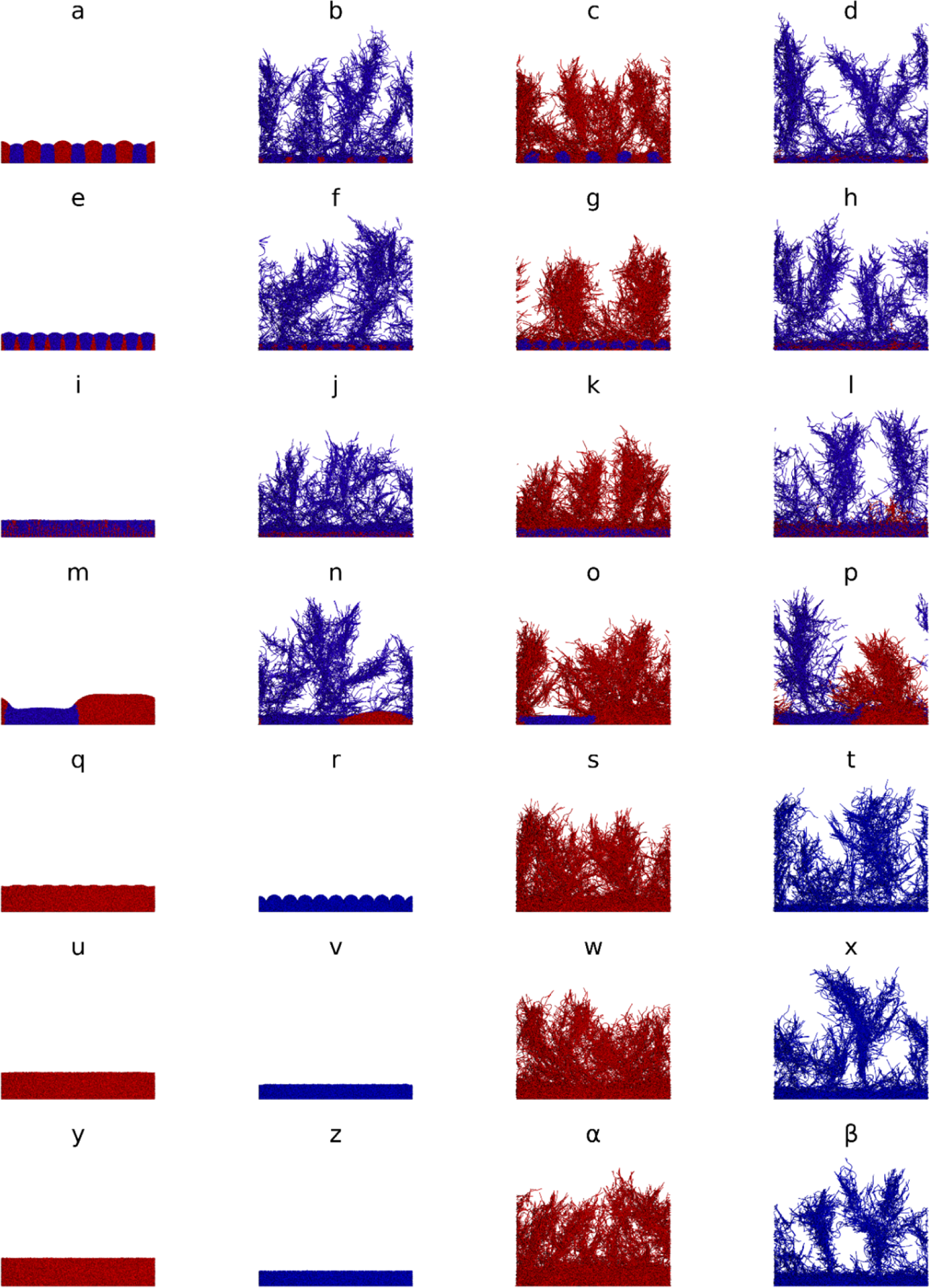
Filaments rule and gave Rate Strategists an advantage. Rate Strategists (RS, blue) and Yield Strategists (YS, red) competed in a 3D biofilm domain (200×200×12.5 µm) for 3 weeks. In the first 4 rows, strategies competed. Column 1 corresponds to spherical cell scenarios in Fig 2 of Ref [46] but were now simulated in 3D. In column 2, RS formed filaments and in column 3, YS formed filaments. Filaments won regardless of strategy. In column 4, both formed filaments and RS won or likely won. The last 3 rows show single species ‘controls’ with 10, 20 or 100 initial agents. The first two columns show simulations with spherical YS or RS agents while the last two columns show filament forming YS or RS agents. See Fig 7 for corresponding time courses. Duplicate simulations are shown in Fig SI7.

**Fig 7.**
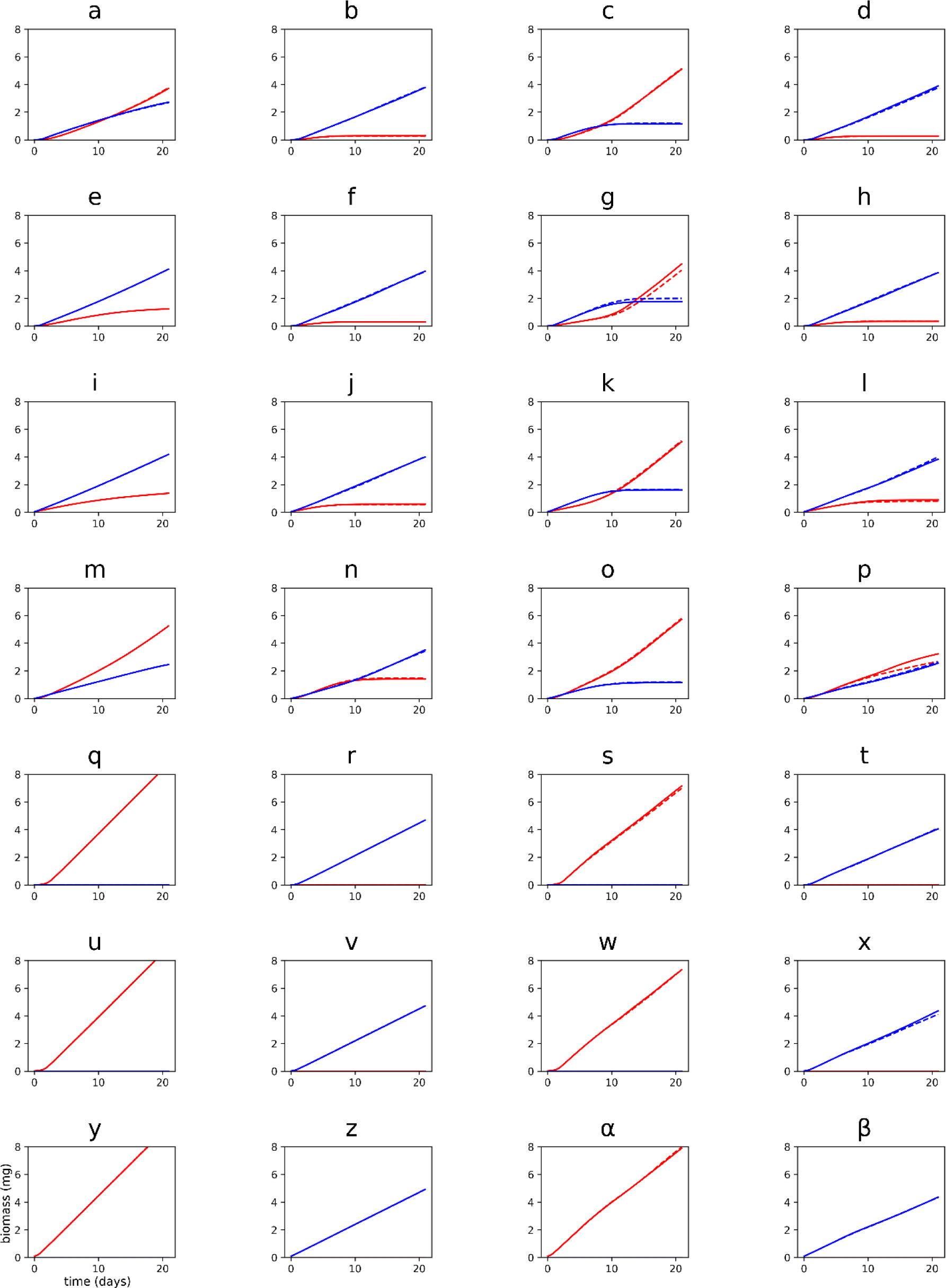
Growth curves corresponding to competitions of Rate Strategists (RS, blue) and Yield Strategists (YS, red) in. Fig 6. Duplicates are plotted with dashed lines. Divergence between replicates is most visible in panels g and p. In panel p, it is too early to definitely call the outcome of competition, but it is likely that RS would win given the biofilm structure after 3 weeks (Fig 6P).

## Discussion

Individual-based models in microbial ecology are uniquely capable of capturing local interactions, individual heterogeneity and adaptation, stochastic processes and emergent properties of biofilms or other spatially structured assemblages [49–58]. The number of publications utilizing IbMs in this field has rapidly increased since the 1990s (S1.13 Fig S10), due to an increased recognition of its potential and facilitated by an increased availability of ready-to-use IbM platforms (discussed below). From its inception, the goal of iDynoMiCS has always been to provide a well-tested, general-purpose platform for individual-based microbial community modeling, enabling users to specify their model in a structured text file rather than requiring them to program, thereby aiding microbial ecology and synthetic biology research, which seeks to reach a mechanistic and predictive understanding of the interactions of natural or synthetic microbes in the environment. The environment for microbes includes engineered reactors and buildings as well as plant and animal hosts. Here, we present iDynoMiCS 2.0, which has been rewritten from scratch to enable flexible agent and process specifications using orthogonal modules called “aspects”, thus removing key limitations of the original iDynoMiCS 1. In addition, we expanded the functionality, most importantly a force-based mechanical interaction framework enabling new agent morphologies such as rods and filaments, a new flexible approach to (bio-)chemical conversions enabling any type of kinetic expression, improvements that make the software easier to use including a GUI, a new protocol and program structure, unit conversions and a large number of efficiency improvements allowing for large scale (10+ million agents) simulations. We have further subjected iDynoMiCS 2.0 to rigorous testing to ensure that the platform and its individual components function correctly. In addition to standard unit tests, we have verified the accuracy of its numerical solvers by using a series of increasingly complex test cases from simpler ones with analytical solutions to more complex ones that have to be compared to independent solvers, culminating in the established Benchmark 3 comparison of nitrifying biofilms with different biofilm modeling approaches and a comparison with prior BacSim simulations of the biofilms promote altruism test case because its positive feedback in the growth of cooperative cell clusters results in higher sensitivity to local cluster formation in biofilms.

Biomass spreading mechanisms can affect model outcomes, especially when mixing of different species favors a species that can take advantage of becoming embedded in a cluster of cells of the other species, e.g., if the latter is more rapidly growing towards an energy source. Using nitrifying biofilms, it was previously shown that Cellular Automata (CA) cell division rules led to larger scale stochastic mixing of nitrite oxidizers into the (under specific conditions) more rapidly expanding incomplete ammonia oxidizers than a shoving algorithm that minimized cell-cell overlap, which percolated expansion through the biofilm leading to limited, localized mixing. This resulted in a substantially higher fitness of the nitrite oxidizers in the CA simulations as some of them were ‘going with the flow’ of the ammonia oxidizers towards the oxygen supply [15]. Here, we compared this shoving algorithm with FbM, although using the biofilms promote altruism case study rather than nitrifying biofilms. FbM led to even more limited and localized mixing, producing sharper boundaries between different clusters than shoving (Fig 5). As a result, the chance of clusters of yield strategists to emerge out of clusters of rate strategies (after the latter had overgrown them) and thus eventually winning the competition, was reduced. The biofilm structures resulting from FbM are reminiscent of the smooth, gradual boundaries of biomasses in continuum models of biofilms where biomass spreading is proportional to the pressure gradient [59] (Darcy’s law), but the mixing is more limited with the FbM. Note that continuum models where biomass spreading is driven by density-dependent diffusion can lead to complete mixing if counter-diffusion is not considered and gradual but large-scale mixing if it is [60]. Combining our new results with Kreft et al. (2001) [15], we can rank biomass spreading mechanisms in order of increasing and larger-scale mixing: FbM < shoving < CA. The importance of even subtle differences in biomass spreading mechanisms for biofilm pattern formation and population fitness should become more widely recognized as model predictions can be substantially affected.

Morphology matters. There is a great variety of shapes [61] and sizes [62] of microbes while microbes of the same species usually have similar shape and size. Young [63] argued that the variety and uniformity of microbial morphology, the ability of bacteria to actively modify their shapes based on internal or external cues and evolutionary selection towards specific shapes all suggest that bacterial morphology is as important as other traits. Different morphologies may be selected for by different selective pressures and ecological niches such as nutrient limitation, predation, attachment, passive dispersal, active motility and differentiation [63,64]. IbMs have previously been utilized to demonstrate how cell shape can play an important role in spatial organization and evolutionary fitness in biofilms [65,66]. Sphere-shaped, rod-shaped and filamentous microbes are commonly found and can already be modeled with iDynoMiCS 2.0 and the implemented ball-spring approach facilitates future extensions to branching filaments or other morphologies.

Filaments win. Our filament case study utilized FbM to simulate filaments consisting of sphere-shaped agents and demonstrated that filamentous growth can provide a strong competitive advantage under nutrient limiting conditions (Fig 6). The advantage derives from the ability of filaments to focus the growth of biomass into one direction rather than merely producing offspring adding to an existing heap of cells. This way, longer distances can be covered and new, nutrient rich territories colonized. This is similar to the strategy of cord-forming fungi who can quickly grow towards new resource hotspots [48,67] and microbes that push themselves towards the nutrient source by producing copious amounts of low-density EPS [47]. The advantage of clusters of yield strategists, who compete less and grow faster than a cluster of rate strategists (of the same size and at the same flux of resources into the cluster), turned into a disadvantage as rate strategists who have reached the top of the biofilm where substrate flux was highest experienced stronger positive feedback than yield strategists. This suggests that filamentous growth is a strategy to escape competition between siblings. Given the huge advantage of filamentous growth found here, the question why filamentous growth is not more common in bacteria arises. It is certainly common in fungi and in the ecologically similar streptomycete bacteria, probably because of improved foraging for scattered patches of resources separated by terrain low in suitable resources like oases in a desert. Also in stream biofilms, filament or chain formation as employed by *Diatoma* spp. enhances nutrient access [29]. Gradient microbes such as *Beggiatoa* spp. or the intriguing cable bacteria [68] form filaments because they need to access electrons from a reduced sediment and electron acceptors from the oxidized water layer above the sediment. Filamentous bacteria are also found in activated sludge flocs in wastewater treatment, where they have the advantage of growing out of the floc into the nutrient richer bulk liquid but are selected against at the settling stage where only fast sinking sludge flocs are recirculated into the activated sludge stage [69,70]. But what are the disadvantages of filaments? Depending on the environment, several disadvantages may arise. Filaments are not only more exposed to beneficial resources but also mechanical or physiological stresses and attack by phages or predators, although some predators may be deterred from grazing larger cells [63]. Further, packaging biomass into smaller propagules is advantageous for dispersal. Filaments also forsake the advantages of motility [63,71]. Cell size is also an important factor for pathogenesis, some bacteria may avoid forming filaments to prevent being killed by the host [64].

iDynoMiCS 2.0 is joining an increasing number of individual-based modeling platforms that focus on microbes and can support a range of specific models. These platforms can be roughly divided into two groups, based on their subcellular versus ecological dynamics origin and focus. The former group of platforms comes from systems and synthetic biology and seek to discover how specific microbial community behaviors or phenomena can be achieved through the creation of synthetic microbial communities [72]: CellModeller [73], BSim 2.0 [74] and gro [75]. They can simulate microbial communities made up of rod-shaped microbial agents with specific metabolic, sensing and signaling properties. All three can simulate gene regulatory networks and diffusion of signaling molecules in order to explore and/or design synthetic microbial communities. While gro can only simulate 2D systems, CellModeller can simulate both 2D and 3D systems, while BSim 2.0 can only simulate 3D systems. In models that use these platforms, growth kinetics are typically less important than gene regulation, hence growth is modeled as a simple rate, as in CellModeller, or a rate based on cell length, as in BSim 2.0. However, gro allows growth to be based on Monod kinetics. CellModeller and gro do not include environmental constraints such as physical boundaries, thus simulated microbes tend to grow outwards to form round colonies. BSim 2.0, however, can model physical spaces such as microfluidic chemostats where cells may, e.g., grow and release diffusing signaling molecules.

The latter group of platforms originate from larger scale microbial ecology models, primarily for biofilms, which seek to explore population dynamics and ecological and evolutionary processes in biofilms. These include iDynoMiCS [17], Biocellion [76], Simbiotics [77], BacArena [78], NUFEB [21], ACBM [79] and McComedy [23]. They focus on microbial growth and metabolism and mass transport such as diffusion of solutes in order to assess how different growth strategies or metabolic interactions affect the fitness of species growing in a biofilm or impact systems-level outcomes in wastewater treatment systems or bioreactors. iDynoMiCS, NUFEB and Simbiotics can all model growth using equations originating from enzyme kinetics that determine reaction rates from substrate concentrations, such as Monod kinetics. Reaction rates and diffusion are coupled and solved using partial differential equation (PDE) solvers. These solvers are made efficient by taking advantage of a separation of timescales, where, e.g., growth of microbes is on a much slower timescale than diffusion. iDynoMiCS and NUFEB both simulate a diffusion boundary layer - a region around the biofilm in which diffusion dominates the transport of solutes - as distinct from the rest of the spatial domain which is well-mixed.

BacArena and ACBM are unique in utilizing flux-balance analysis to estimate the metabolic flux through ‘individual’ grid elements, based on their local solute concentrations. Diffusion is then solved using a partial differential equation (PDE) solver. NUFEB can model agent growth based on thermodynamics, calculating the Gibb’s free energy of catabolism [39], which could also be done with iDynoMiCS 2.0 as users can specify any arithmetic function for reaction kinetics. Since BacArena and ACBM have to solve flux-balances, which is computationally demanding, the platforms are more restrictive in terms of model scale. BacArena only models agents in a fixed 2D grid, with one agent per grid cell, like a CA, while ACBM groups agents together when evaluating internal processes. The other biofilm modeling platforms simulate grid-free agents that evaluate internal processes on an individual basis. Agents can excrete small particles representing EPS. NUFEB, Simbiotics iDynoMiCS 2.0 also allow adhesive forces to be modeled. In NUFEB and iDynoMiCS 1, agents are spherical, while in Simbiotics, ACBM or iDynoMiCS 2.0, they can be spherical or rod-shaped.

Some of the models can simulate fluid motion or advection in addition to diffusion. CellModeller implements an implicit advection model which imposes a linear bulk flow in a given direction. NUFEB can simulate computationally demanding fluid dynamics explicitly through coupling with the fluid dynamics toolbox OpenFOAM, which can solve the fluid velocities based on the biofilm geometry (one way coupling). Forces are then applied to agents based on these flow velocities.

The suitability of different individual-based modeling platforms depends on the needs of the user. For exploring synthetic bacterial communities where gene regulation and signaling circuits are engineered into cells, CellModeller, gro or BSim 2.0 may be the most suitable platforms. When details of intracellular dynamics are less relevant or simply unknown and the focus is on interactions between agents and with the environment, such as in biofilms or other spatially structured habitats where mass transport is crucial, NUFEB and iDynoMiCS 2.0 may be the most suitable systems. iDynoMiCS 2.0 offers a highly modular and customizable modeling platform, with both 2D and 3D compartments, spherical, rod-shaped and filamentous microbial agents, a sophisticated reaction-diffusion system and growth models that can be based on any kind of arithmetic expression including enzyme kinetic and thermodynamic based models. It is more straightforward to specify and implement biology in iDynoMiCS 2.0. One key drawback in comparison to NUFEB is that iDynoMiCS 2.0 currently does not model fluid dynamics or advection, and thus if these are important characteristics of the system one wishes to model, NUFEB may be more suitable. BacArena and ACBM offer flux-balance models for metabolism, but therefore come with other limitations.

For specific applications, other Agent-based or related bottom-up modeling platforms are worth considering. IndiMeSH [80] is an IbM platform capable of simulating laboratory models of soil habitats. CHASTE [81], BioDynaMo [22], PhysiCell [82] and compuCell3D [83] have been primarily used for tissue modeling, which could make them applicable to the somewhat similar biofilms. Morpheus [84], like compuCell3D, utilizes a cellular Potts model to model multicellular systems. Further, there are several general purpose AbM libraries or toolkits, which facilitate the programming of a specific model by providing a wide range of common routines so models can be specified with a high-level language tailored for this purpose. These libraries could be suitable for certain microbial community models in addition to various other fields of research. They include NetLogo [85], FLAME [86], Mason [87], Repast simphony [88] and others.

iDynoMiCS 2.0 has been developed from scratch because the hierarchical inheritance of agent features in iDynoMiCS 1 prevented the fully flexible pick and mix approach to agent characteristics and processes that was required for further expansion of capabilities. Moreover, iDynoMiCS 2.0 sports several key enhancements, such as the ability to simulate spherical, rod-shaped and filamentous microbes and using Force-based Mechanics for biomass spreading, which we show can have important consequences. It can simulate larger 3D domains due to efficient neighbor searching, a faster converging reaction-diffusion solver and numerous other performance improvements. iDynoMiCS 2.0 was designed with both ease of use, from a GUI to unit conversions, and ease of extension in mind, providing a solid and well tested simulation platform for a wide variety of microbial community studies to come. We showcased the simulation of filamentous microbes using the biofilm promotes altruism case study and found that the rate strategists gained a stronger advantage from filamentous growth because their tips can escape from the stronger competition between themselves. This demonstrates just one of many new possibilities of iDynoMiCS 2.0.

## Acknowledgements

iDynoMiCS has undergone development and testing since 2005 thanks to many researchers in the groups of BFS and JUK and seen improvements by many others. We are grateful to all of them. TF, RJC, KA and JUK would like to thank the United Kingdom National Centre for the Replacement, Refinement & Reduction of Animals in Research (NC3Rs) for funding our development of individual-based modeling platforms for the gut environment (Grants NC/K000683/1 and NC/R001707/1). BJRC and BFS would like to thank the Integrated Water Technology (InWaTech) project, which promotes collaborative research between DTU and KAIST, for funding our work on work on microbial aggregation and granular biofilms. SA was funded by a University of Nottingham Vice Chancellor’s Scholarship. The funders had no role in study design, data collection and interpretation, decision to publish, or preparation of the manuscript.

## Code availability

All source code and data associated with this project is published under the GNU General Public License (GPL) compatible CeCILL license V2 and available as an online repository on https://github.com/kreft/iDynoMiCS-2

## S1 Supporting information

### S1.1 Introduction

The supplementary materials provide additional details on the iDynoMiCS 2.0 framework and the model implementations presented in the main manuscript. This includes a further description of the framework and detailed descriptions of the case studies with their parameters. Moreover, the model verification and benchmarking against prior work is presented.

### S1.2 Detailed description of Force-based Mechanics (FbM) and testing

The force-based mechanical interactions between agents and agents and surfaces in iDynoMiCS 2.0 rely on both correct detection of overlapping agents or collisions and correct responses. Detection is simple for stored interactions such as the interaction between the two points of a rod cell connected by a spring or the interaction between cells in a filament also connected by springs. In this case, detection is as simple as checking whether interaction data is stored as an aspect of the agent. In the case of collisions or attractive interactions, collision detection is utilized. Different shapes as well as periodic boundaries add complexity to this routine. For verification purposes, a total of 36 collision detection scenarios (Table S1) were tested and included in the software as unit-tests.

**Table S1.** In total, 36 collision detection scenarios were included as standard unit tests. All tests include two objects to create one of the following scenarios: object-object overlap (hit), no overlap (miss), overlap through a periodic boundary (periodic hit) and no overlap, but proximity through a periodic boundary (periodic miss). The sphere and rod objects correspond to agent shapes. Solid boundaries utilize an (infinite) plane object to allow for agent interactions. The voxel is a cube aligned with the coordinate grid. Numbers indicate the number of different configurations tested. In all tested scenarios the collision detection algorithm correctly detected the hits and misses.

|  | Sphere +<br>Sphere | Sphere +<br>Rod | Rod +<br>Rod | Plane +<br>Sphere | Plane +<br>rod | Voxel +<br>Sphere | Voxel +<br>rod |
| --- | --- | --- | --- | --- | --- | --- | --- |
| hit | 1 | 1 | 1 | 1 | 1 | 1 | 6 |
| miss | 1 | 1 | 1 | 1 | 1 | 1 | 4 |
| periodic hit | 1 | 1 | 1 | 1 | 1 | 1 | 3 |
| periodic miss | 1 | 1 | 1 |  |  | 1 | 1 |

Correct interaction response entails relaxing mechanical stresses between agents until a relaxed state is reached. Criteria for a relaxed state can either be a threshold value for tolerated residual interaction force in the model state or a threshold value for tolerated agent overlap (µm). In the test in Fig S1, an over-compressed initial state underwent 1,000 FbM iterations using its default parameters. Initial peak interaction forces dropped exponentially to asymptotically approach zero (the maximum residual interaction force (reached after 829 iterations) was less than 0.1 fN).

**Fig S1.**
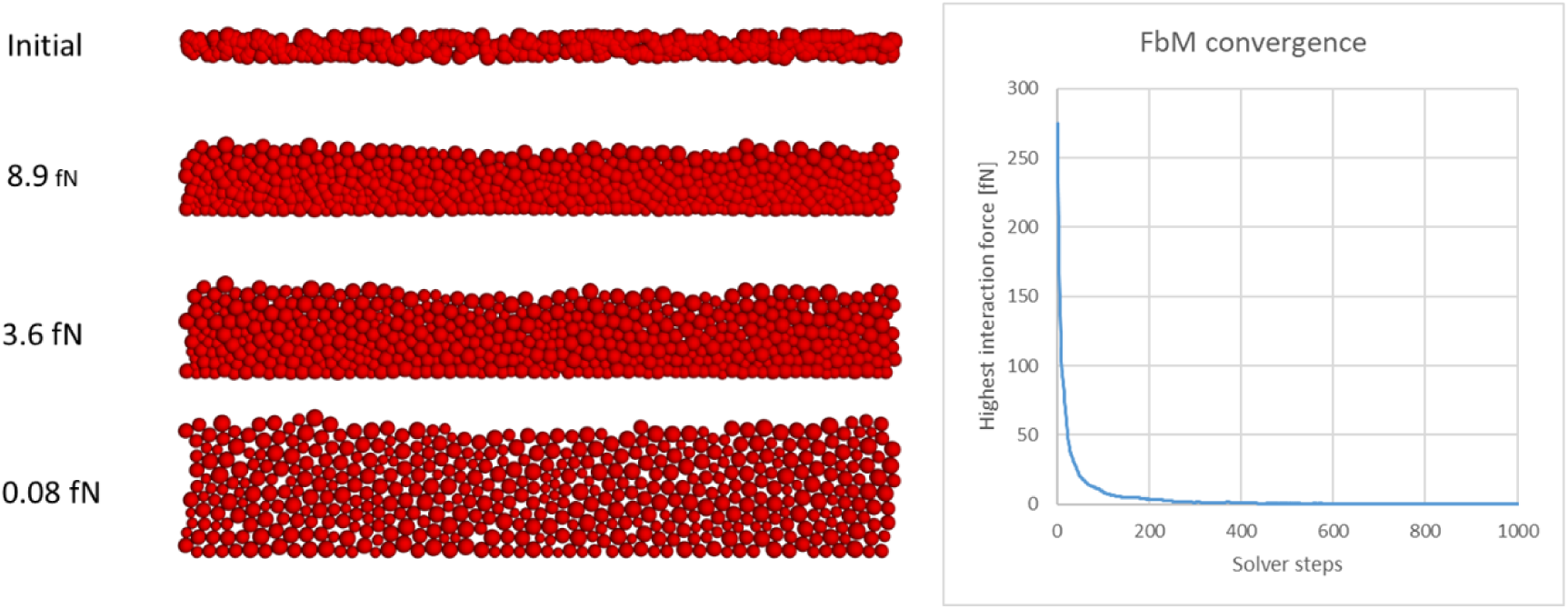
FbM led to rapid relaxation of mechanical stress from an initially over-compressed state. Left panels from the top showing highest interaction force next to the biofilm structure: 275.1 fN for the initial state, 8.9 fN after 100 steps, 3.6 fN after 200 steps, 0.08 fN after 1,000 steps. Panel on the right shows exponential drop of the highest interaction force towards zero, demonstrating convergence of the FbM solver.

### S1.3 Testing Reaction and Diffusion of Chemical Species

In iDynoMiCS 2.0, ODE and PDE solvers are responsible for modeling the diffusion and reaction (consumption or production) of solutes throughout the simulated system and are therefore responsible for the maintenance of mass balance within the model. Reactions can be chemical reactions or catalyzed by individual agents.

To test that these solvers work as intended, a range of test cases were run, which allowed the results from iDynoMiCS 2.0 to be compared with known analytical solutions. The tests were conducted starting with the simplest and proceeding to increasingly complex systems. The first two tests were non-spatial systems, used to test the ODE solver, while the latter two tests were for the more complex PDE solver in a 2D spatial system. All tests are described in full below.

#### 1. Non-growing Catalyst Agent in a Chemostat

For this simplest test, a single agent was simulated in a chemostat compartment, consuming the inflowing solute. Fresh medium with a fixed solute concentration flowed into the chemostat at a fixed flow rate. Spent medium flowed out of the chemostat at a rate equal to the inflow. The consumption of the solute by the agent was proportional to the solute concentration and to the agent’s mass. The agent was neither growing nor removed.

This system can be described by the following differential equation

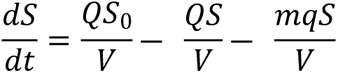

where S is the solute concentration of the substrate in the chemostat, S_0_ is the solute concentration in the inflowing medium, Q is the flow rate with dimension volume per time, V is the volume of the chemostat, t is time, q is the rate of solute consumption by the agent and m is the mass of the agent.

The steady-state solution for this differential equation is

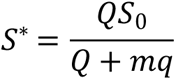

This system was simulated in iDynoMiCS 2.0, with timesteps of 100 minutes. The steady state predicted given parameters in Table S2 was 4.0 × 10^5^ pg µm^-3^ and the simulated concentration converged to this steady state exactly (Figure S2A).

**Table S2.** Parameters used in the numerical tests of the chemostat solver.

| Parameter | Non-growing agent in chemostat | Growing population in chemostat |
| --- | --- | --- |
| $S_0$ | $2.0 \text{ pg } \mu\text{m}^{-3}$ | $2.0 \text{ pg } \mu\text{m}^{-3}$ |
| $Q$ | $1.0 \mu\text{m}^3 \text{ min}^{-1}$ | $1.0 \mu\text{m}^3 \text{ min}^{-1}$ |
| $V$ | $1.0 \times 10^3 \mu\text{m}^3$ | $1.0 \times 10^3 \mu\text{m}^3$ |
| $q$ | $0.1 \mu\text{m}^3 \text{ min}^{-1} \text{ pg}^{-1}$ | |
| $K_s$ | | $1.0 \mu\text{m}^3 \text{ min}^{-1} \text{ pg}^{-1}$ |
| $m$ | $40 \text{ pg}$ | $10 \text{ pg}^*$ |
| $\mu_{\text{max}}$ | | $0.1 \text{ min}^{-1}$ |
| $Y$ | | $0.5 \text{ pg } \text{pg}^{-1}$ |

\*The mass for the simulation of the growing population refers only to the initial mass.

#### 2. Growing Population in a Chemostat

In this simulation, a growing population of agents in a chemostat consumed an inflowing substrate and converted it to biomass. Outflow removed both spent medium and agents, at a rate equal to the inflow. Agent removal was stochastic. The agents consumed substrate and grew according to Monod kinetics:

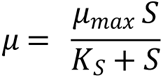

Where *µ* is the specific growth rate, *µ_max_* is the maximum specific growth rate and K_S_ is the half-saturation constant, the value of S at which µ = µ_max_/2.

Here, the rate of change of substrate concentration is given by

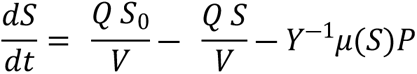

where *Y* is the biomass yield from the substrate and P is the concentration of the biomass of all (planktonic) agents in the chemostat, with the rate of change given by

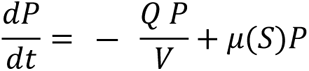

This system can be solved to find the steady states for both *P* and *S*, the washout steady state of *P*^∗^ = 0, *S*^∗^ = *S*_0_ [89], and the steady with agents present:

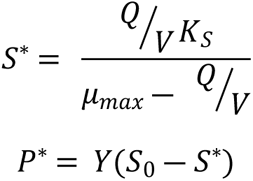

With the parameter values in Table S2, we obtain the following steady state predictions:

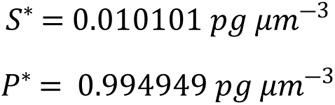

Running the simulations in iDynoMiCS 2.0 yielded the expected stable steady state (Figure S2B,C). The mean simulated values at steady state were S* = 0.9906 pg µm^-3^ and 0.0101 pg µm^-3^. These results differ from the expected steady states by 0.004% and 0.0003%, respectively.

#### 3. Thin Layer of Non-growing Cells in a Spatial Domain

In this test, a thin non-growing layer of cells, occupying one row of solver grid elements, was simulated at the bottom of a spatial compartment, with a concentration boundary layer above the cells, and a well-mixed region above that with the constant concentration of substrate *S_0_*. Since there was no gradient of biomass or reaction rates in the horizontal direction, this is effectively a 1D system for which an analytical solution for the flux, J, can be calculated according to Fick’s first law

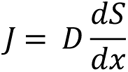

where *J* is the areal flux density through the diffusive region, *D* is the diffusivity of the solute *S* and *x* is the vertical distance (the direction for the flux and substrate concentration gradient).

Given that at steady state, flux must be constant along the x-axis in the region where the substrate is not consumed and then starting to decline where the substrate is consumed by the cells at the bottom, we can substitute J by the areal consumption rate at the cell-layer surface. Modeling a simple consumption rate proportional to biomass and substrate concentration, we obtain

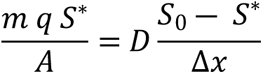

where S* is the steady-state concentration at the biofilm surface, A is the surface area of the biofilm and *Δx* is the depth of the diffusion-dominated boundary layer. This can be rearranged to

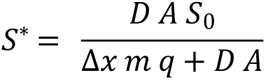

Setting the parameters as shown in Table S3, the predicted steady state concentration at the cell layer surface is *S*^∗^ = 1.8 × 10^―6^*pg μm*^―3^. This was matched in the simulation (Figure S2C). Deviations from the expected concentration are very small at each height, with the greatest deviation of 0.017% at a height of 8 µm.

**Table S3.**
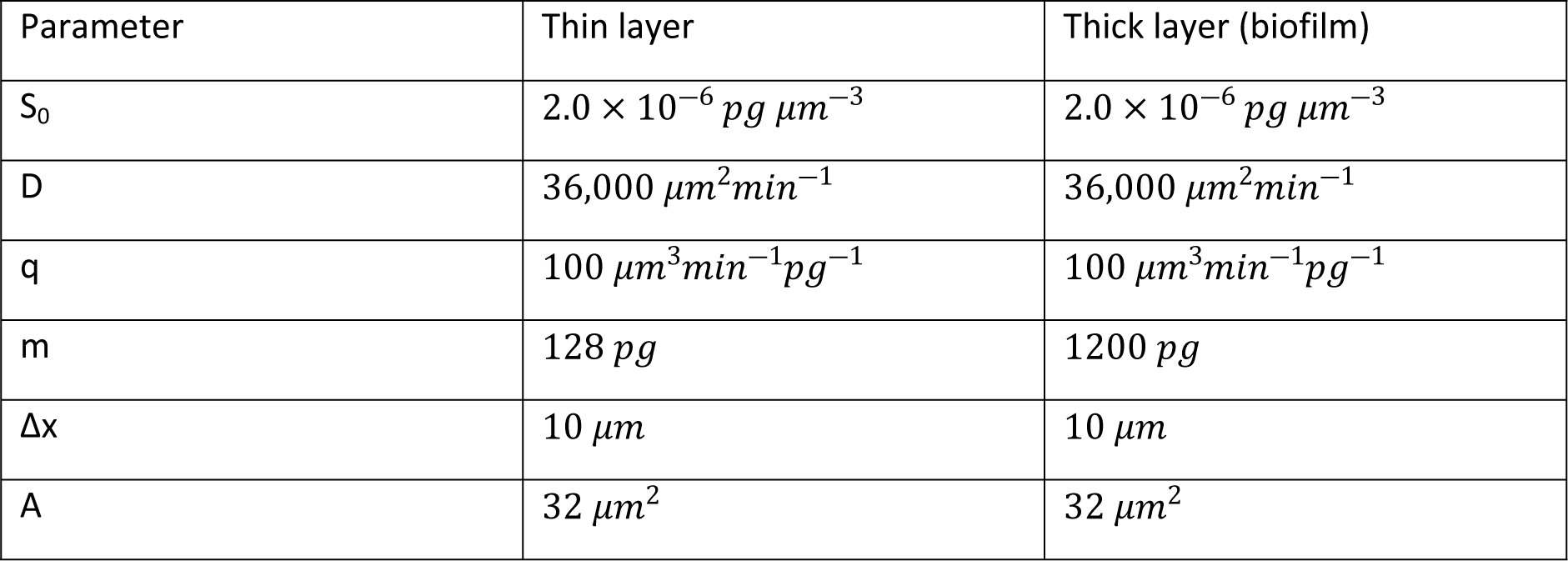
Parameters used in the numerical tests of the spatial domain in iDynoMiCS 2.0. The total biomass was higher for the thick layer, all other parameters were identical.

##### Biofilm - Thick Layer of Non-Growing Cells in a Spatial Domain

For a biofilm simulation with a thicker layer of cells, no analytical solution is available for the solute concentration at the surface of the biofilm. However, the nature of the boundary at the bottom of the domain, an inert, solid and flat surface with a no-flux (Neumann) boundary condition, provides another testable feature. As a result of the thick biofilm layer consuming substrate while it diffuses towards the bottom, the concentration gradient is expected to decrease from the maximum level in the diffusion boundary layer to become zero at the inert surface. The results of the test replicated the predicted features of the concentration gradient (Figure S2D), suggesting that the diffusion-reaction solver and the no-flux boundary conditions in iDynoMiCS 2.0 are functioning as expected. There was no horizontal gradient or any unexpected deviations at the horizontal, periodic boundaries.

**Fig S2.**
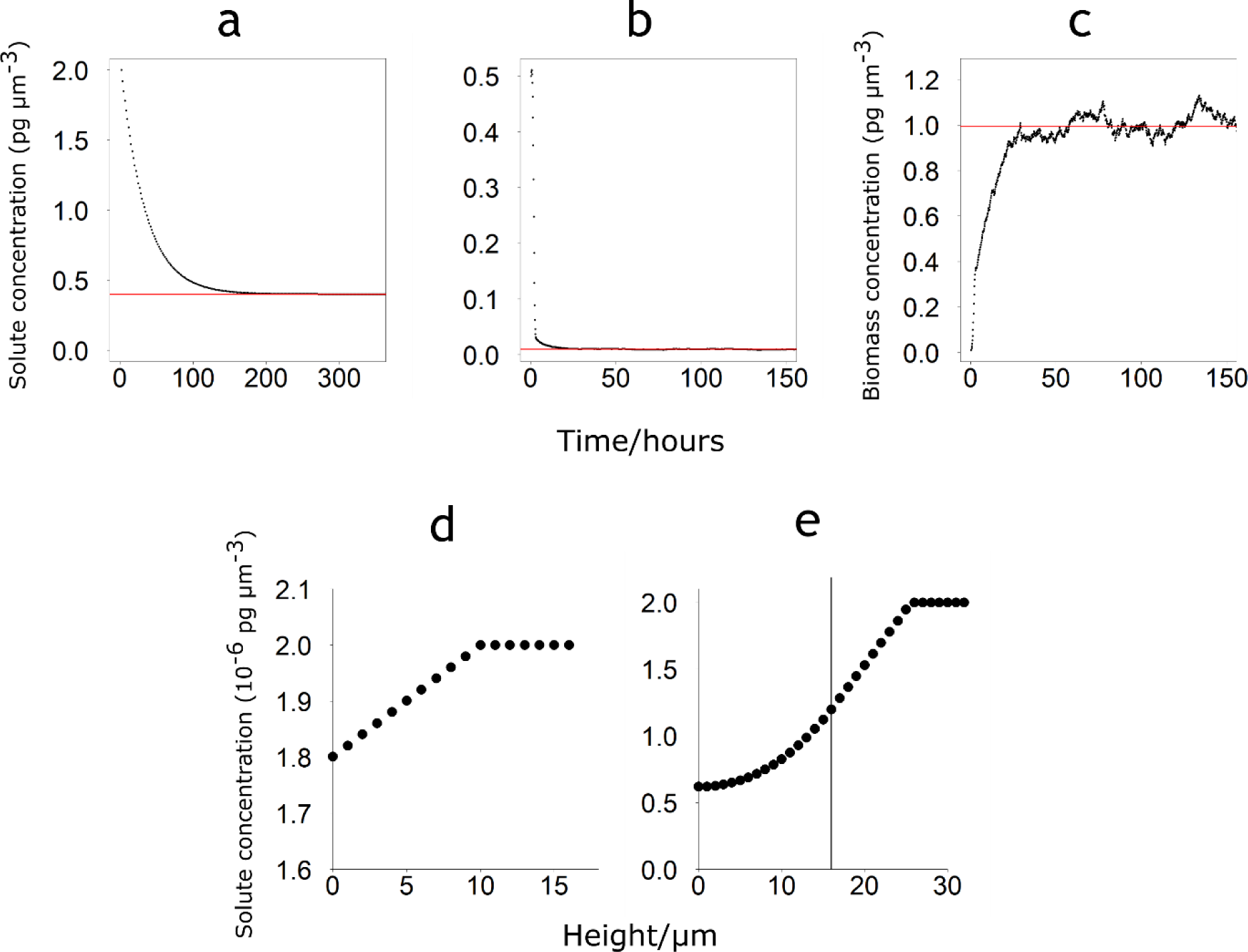
Results of numerical tests of the ODE and PDE solvers. Red lines show expected steady states. a – Results from the non-growing chemostat population. Concentration tended towards the expected steady state of 0.4 pg µm^-3^. b, c – Results from the growing chemostat population. Concentrations tended towards the expected steady states of 0.0101 pg µm^-3^ substrate (b) and 0.9949 pg µm^-3^ biomass (c). de) – Results from thin cell layer. Concentration at biofilm surface matched predicted concentration of 1.8 × 10^-6^ pg µm^-3^. e – Results from thick cell layer. Vertical line marks biofilm surface. The substrate concentration gradient was linear in the diffusion boundary layer above the biofilm surface and then decreased towards zero at the inert boundary at height 0, as expected.

### S1.4 Large scale stress-test

A two-species nitrifying biofilm model was set up to test the ability of iDynoMiCS 2.0 to simulate larger scale domains. The kinetics are based on Hubaux et al. [90]. A 500×500×500 µm spatial compartment with fixed concentrations at the top of the domain was initiated with 1,000 Ammonium Oxidizing Organisms (AOO) and 1,000 Nitrite Oxidizing Organisms (NOO), randomly distributed over the inert surface at the bottom of the spatial compartment. Model parameters are given in Table S4 and the stoichiometry and process kinetics are given in the Petersen matrix in Table S5.

**Table S4.**
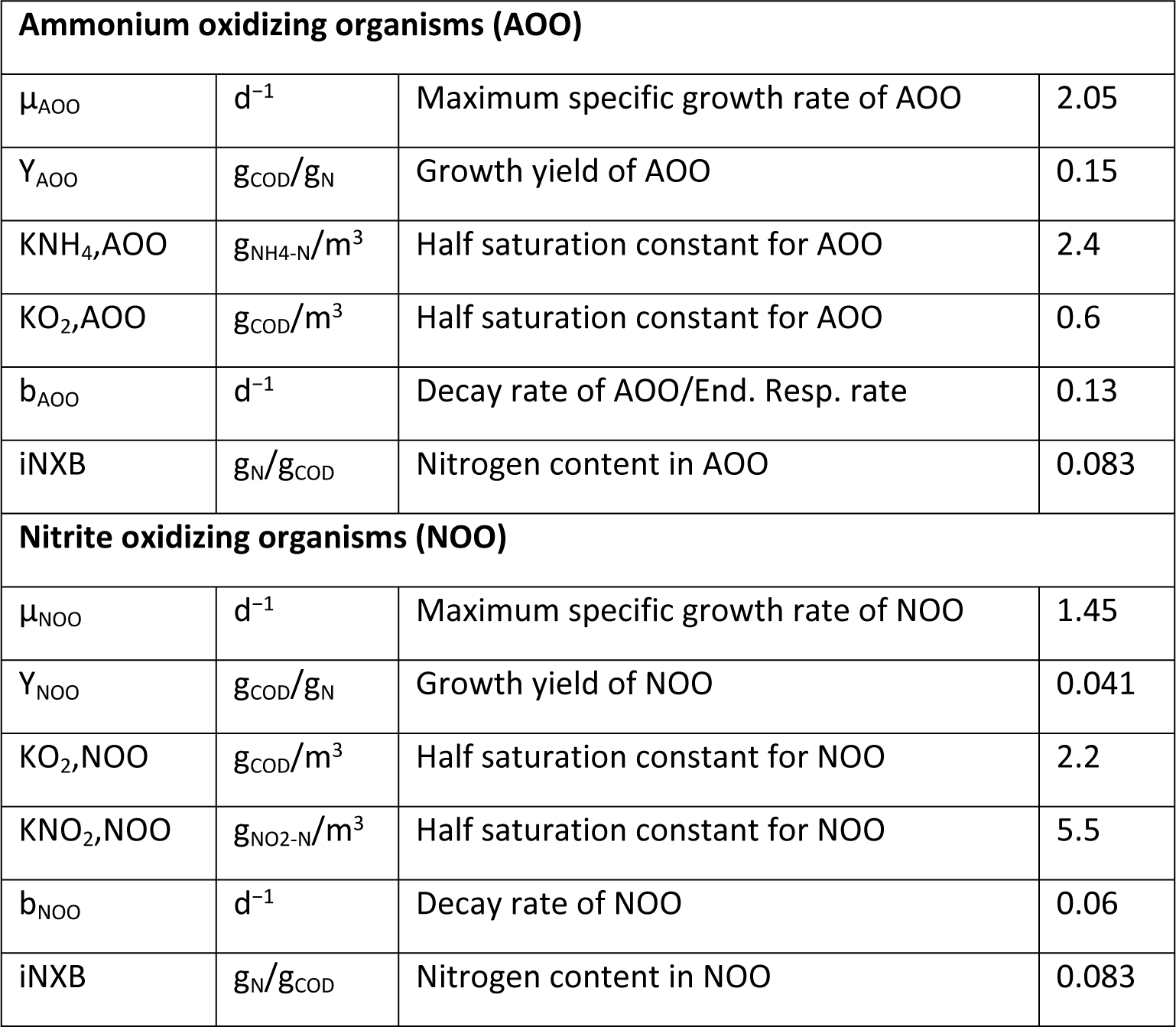
Parameters for the two-species nitrifying biofilm model. All kinetics for this model are based on Hubaux et al. [90].

| **Ammonium oxidizing organisms (AOO)** |  |  |  |
| --- | --- | --- | --- |
| $\mu_{AOO}$ | $d^{-1}$ | Maximum specific growth rate of AOO | 2.05 |
| $Y_{AOO}$ | $g_{COD}/g_N$ | Growth yield of AOO | 0.15 |
| $K_{NH_4,AOO}$ | $g_{NH_4-N}/m^3$ | Half saturation constant for AOO | 2.4 |
| $K_{O_2,AOO}$ | $g_{COD}/m^3$ | Half saturation constant for AOO | 0.6 |
| $b_{AOO}$ | $d^{-1}$ | Decay rate of AOO/End. Resp. rate | 0.13 |
| $i_{NXB}$ | $g_N/g_{COD}$ | Nitrogen content in AOO | 0.083 |
| **Nitrite oxidizing organisms (NOO)** |  |  |  |
| $\mu_{NOO}$ | $d^{-1}$ | Maximum specific growth rate of NOO | 1.45 |
| $Y_{NOO}$ | $g_{COD}/g_N$ | Growth yield of NOO | 0.041 |
| $K_{O_2,NOO}$ | $g_{COD}/m^3$ | Half saturation constant for NOO | 2.2 |
| $K_{NO_2,NOO}$ | $g_{NO_2-N}/m^3$ | Half saturation constant for NOO | 5.5 |
| $b_{NOO}$ | $d^{-1}$ | Decay rate of NOO | 0.06 |
| $i_{NXB}$ | $g_N/g_{COD}$ | Nitrogen content in NOO | 0.083 |

The simulation was run on a single core of an Intel Xeon E5 2660 processor with 256 GB memory, the biofilm surpassed 10 million agents after 11 days and 8 hours CPU time, less than the 171 days of simulated time of biofilm development. The simulation was stopped after 175 days of simulated time. The AOO and NOO populations initially grew exponentially as long as growth was not limited by substrate influx and then grew linearly (Figure SI3) while being limited by substrate influx, until reaching a steady state after around 100 days simulated time due to the balancing of overall growth and decay rates with only minor fluctuations in population size. There was no decline as bulk concentrations were kept constant. EPS and inert agents were assumed not to decay in this model, consequently these agent populations continued to increase.

**Fig S3.**
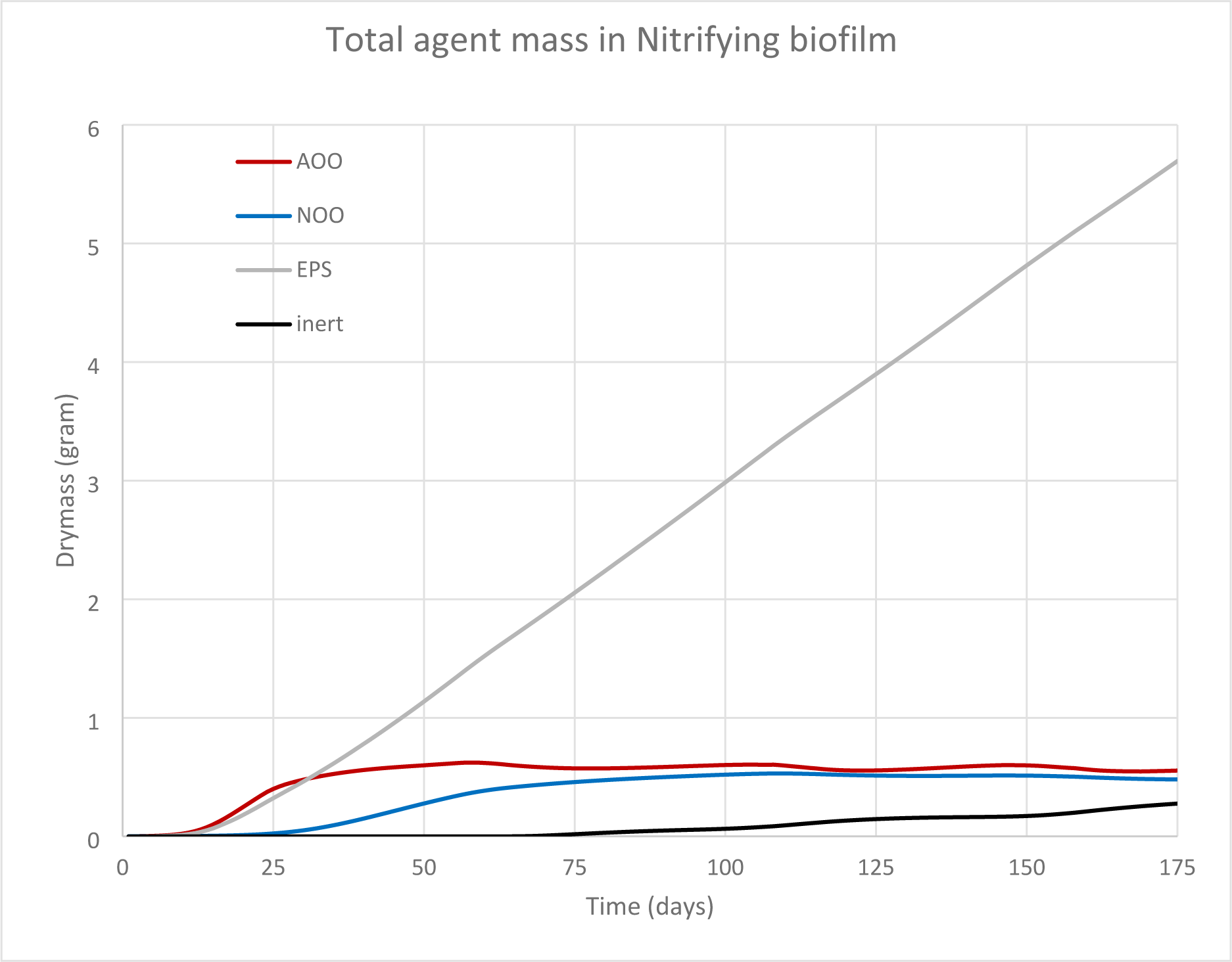
Agent mass in the large scale stress test simulation of a biofilm in 3D. The Autotrophic nitrifying biofilm was initiated with 1 mg Ammonium Oxidizing Organisms (red) and 1 mg Nitrite Oxidizing Organisms (blue). Both species produce EPS particles (gray). Agents that drop below 20% of their division mass as a result of endogenous respiration/decay became inactive (black). The 175-day biofilm contains 1.02×10^7^ agents.

**Table S5:**
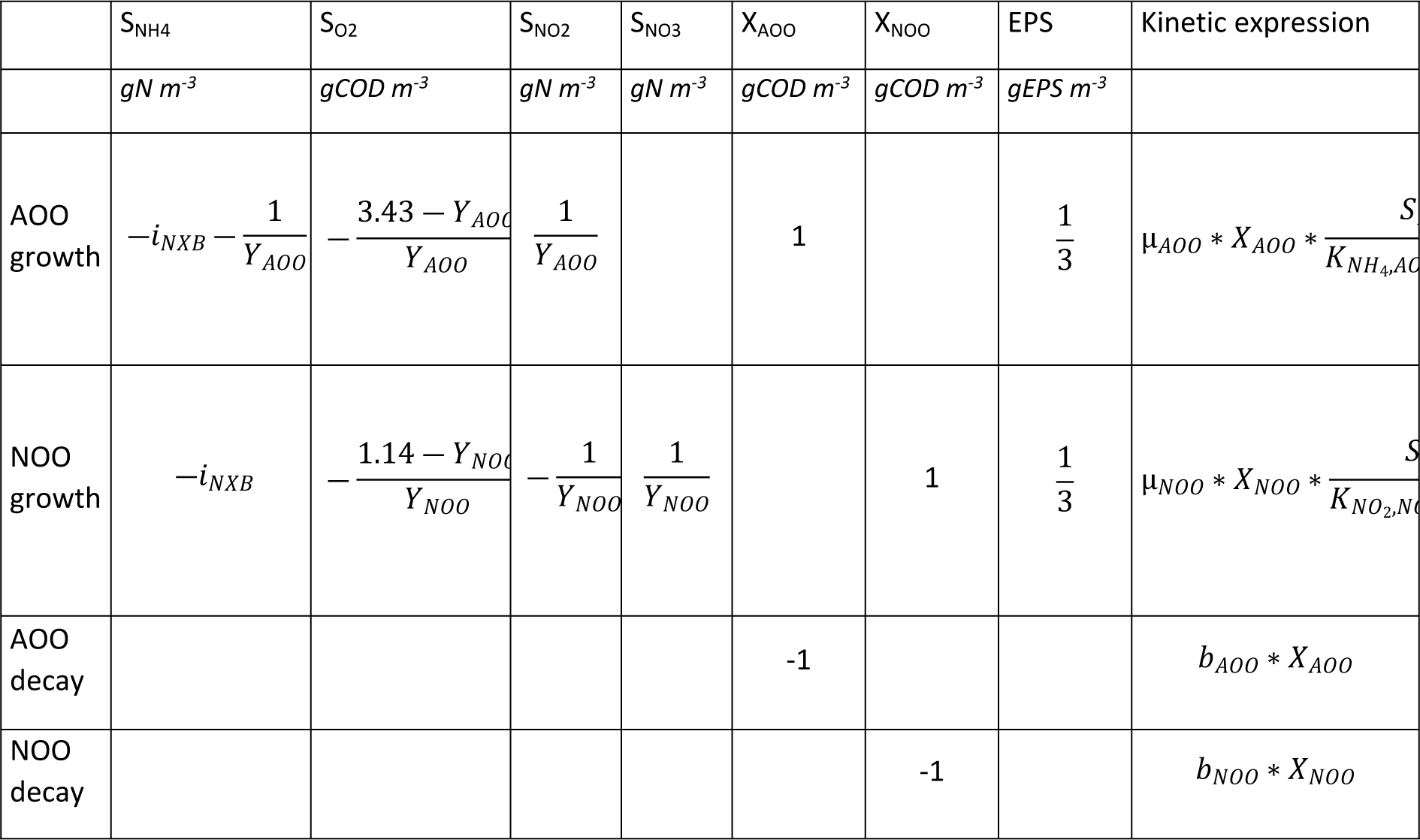
Petersen (stoichiometric) matrix for reactions in the stress test.

### S1.5 Benchmark 3, a comparison of biofilm modeling platforms

One of the longest established applications of biofilm modeling is modeling the treatment of wastewater. The International Water Association (IWA) set up a Biofilm Modeling Task group to compare computational modeling approaches to biofilms and provide guidance for researchers seeking to simulate biofilms. One of the key outputs of this work was the development of a series of Benchmark models – biofilm systems that could be modeled in a variety of modeling platforms to facilitate comparisons between different modeling approaches and establish the effects of different model designs and simplifying assumptions on simulation outputs [7]. The most complex of a set of benchmarks, Benchmark 3 (BM3), was designed to simulate microbial competition in a biofilm, with a source of chemical oxygen demand (COD) being oxidized by a population of heterotrophs and a population of autotrophs oxidizing ammonia to nitrate. This can be thought of as lumping two-step nitrification by ammonia- and nitrite-oxidizing organisms into a single process or modeling one-step nitrification by comammox (complete ammonia oxidizers). BM3 is necessarily limited to what all models are capable of simulating.

The IWA Biofilm Modeling Task Group ran BM3 simulations on a wide range of modeling platforms, with a variety of different approaches to modeling biofilms. Later, BM3 was also used for model validation in the development of iDynoMiCS [17] and NUFEB [21]. Here, iDynoMiCS 2.0 is compared against a selection of four models from the original IWA task group, as well as NUFEB and iDynoMiCS 1. A summary of the different models and their approaches to BM3 follows:

- W – a one-dimensional continuum biomass model run on the AQUASIM software [91] and developed by Peter Reichert and Oskar Wanner [92,93]
- M1 – a variant of the W model with a fixed boundary-layer thickness by Eberhard Morgenroth et al. [94]
- DN – a two-dimensional cellular automaton model developed by Daniel Noguera and colleagues [95]
- CP – a two-dimensional individual-based model, with biomass spreading via shoving, developed by Cristian Picioreanu and colleagues [96]
- NUFEB – A three-dimensional individual-based model that uses a platform derived from a molecular dynamics simulator by Li et al. [21]
- iDynoMiCS 1 – An individual-based model by Lardon et al. [17] used here for 2D simulations. This platform is the precursor to the one described in this paper, and the implementation of BM3 is very similar

As this set of modeling platforms represents a variety of different modeling approaches, they provide a valuable set of results against which to compare iDynoMiCS 2.0. The BM3 scenario has previously been used by Lardon et al. [17] and Li et al. [21] to benchmark iDynoMiCS 1 and NUFEB, respectively. Note that NUFEB has also been directly compared with iDynoMiCS 1 based on the BM3 scenario but varying seven model parameters sampled with a Latin hypercube.

As this set of modeling platforms represents a variety of different modeling approaches, they provide a valuable set of results against which to compare the results of the BM3 simulation in iDynoMiCS 2.0. A description of the implementation of the BM3 model in iDynoMiCS 2.0 follows, henceforth referred to by the abbreviation BM3-iD2.

#### BM3-iD2 Model Description

Previous descriptions of BM3 did not explicitly state two critical details that we had to infer by trial and error. One was that the oxygen concentration in the bulk liquid was kept constant and the other was that the biomass density of the biofilm had to be tuned by scaling the biomass density of the agents. Hence, to facilitate reproduction, we give a full description of BM3 here, using the ODD protocol as a framework, with parts of the description that are already covered by the ODD description of iDynoMiCS 2.0 omitted. The description of BM3-iD2 follows the description of BM3 in Wanner et al. [7].

#### Overview

##### Purpose and patterns

This model simulates multi-species biofilms growing in an aqueous environment as commonly found both in nature and in treatment systems for wastewater and drinking water. The biofilm is composed of two species representing microbial functional groups – an aerobic heterotroph and an aerobic autotrophic nitrifier. Both of these species undergo inactivation processes which transform an agent’s active biomass to inert biomass, meaning that there are three types of biomass present in the biofilm: heterotrophic, autotrophic and inert. The two microbial species compete for oxygen and for space in the biofilm and are transformed into the same inert biomass, leading to vertical stratification of the three different types of biomass through the biofilm.

The purpose of the BM3-iD2 model is to allow comparison between iDynoMiCS 2.0 and other biofilm models. Previous publications did not report time series and only some reported biomass distributions, which limits comparisons to various characteristics of the steady state, including solute concentrations in the bulk liquid, biomass concentrations and to some extent biomass distribution. A close match to other implementations of BM3 would demonstrate that differences in biomass spreading mechanisms between the models have little impact on overall transformation and growth rates in the biofilms and suggest that iDynoMiCS 2.0 is a reliable modeling platform. Deviations would suggest that differences between models, primarily different biomass spreading mechanisms, could affect predictions of overall biofilm performance.

**Table S6.** Parameters used in the Benchmark 3 simulations.

| Parameter | Value |  |  |
| --- | --- | --- | --- |
|  | Standard case | High ammonium | Low ammonium |
| Ammonium influent concentration | $6 \text{ g m}^{-3}$ | $30 \text{ g m}^{-3}$ | $1.5 \text{ g m}^{-3}$ |
| Dilution rate | $0.0111 \text{ min}^{-1}$ | | |
| Volume of bulk liquid | $4.0 \times 10^6 \mu\text{m}^3$ | | |
| COD influent concentration | $30 \text{ g m}^{-3}$ | | |
| Carrier surface area | $320 \mu\text{m}^2$ | | |
| Biofilm thickness | $500 \mu\text{m}$ | | |
| Constant oxygen concentration in bulk liquid | $10 \text{ g m}^{-3}$ | | |
| Biofilm density | $10 \text{ g L}^{-1}$ | | |
| Agent density Shoving | $12.5 \text{ g L}^{-1}$ | | |
| Agent density FbM | $10.08 \text{ g L}^{-1}$ | | |
| Agent division dry mass | $4 \text{ pg}$ | | |
| Boundary layer thickness | $0 \mu\text{m}$ | | |
| Shove Factor | 1.05 |  |  |

##### Entities, State Variables and Scales

The computational domain for BM3-iD2 is a 2-dimensional, spatially explicit compartment with a width of 320 µm. As detailed in Submodels, 2D simulations have a virtual third dimension with a thickness of 1 µm, meaning the effective surface area at the base of the biofilm is 320 µm^2^. This domain represents a vertical slice of a biofilm that contains all simulated microbial agents. In order to maintain the defined biofilm thickness of 500 µm, all agents with a central point greater than 500 µm above the base are removed from the simulation at the beginning of each simulated time step. The biofilm compartment is coupled to a well-mixed bulk liquid compartment with a volume of 0.4 µL, which receives a constant inflow of 0.26 µL h^-1^, with outflow of the bulk liquid at the same rate. Inflowing bulk liquid contains three solutes at fixed concentrations: organic carbon measured as chemical oxygen demand (COD) at 30 g m^-3^, oxygen at 10 g m^-3^ and ammonium at three different concentrations (Table S6). Solutes are well-mixed in the bulk compartment and in the upper portion of the spatial domain above the boundary layer. In the portion of the spatial domain that contains the boundary layer and biofilm, solutes diffuse through a grid with a resolution of 20 µm. The principal agents in BM3-iD2 are the microbial agents, of which there are two types – autotrophs and heterotrophs. Both species are modeled as spherical cells (coccoids), with a division mass of 4 pg. Agent biomass is composed of active and inert portions for both species.

Most models in the original IWA task group could directly set a biofilm biomass density as a parameter, and this is defined in BM3 as 10 g L^-1^. However, as iDynoMiCS-2 is an individual-based model, users can only set the density of agents, with biofilm density an emergent property. In order to match the biofilm density in other models, simulations were run with a variety of agent densities until a biofilm density matching the other models was obtained. Since the emergent biofilm density depended on the agent relaxation method used, in simulations using shoving, an agent (cellular) biomass density of 12.5 g L^-1^ was used, while in simulations using Force-based Mechanics, an agent biomass density of 10.08 g L^-1^ was used. It was also discovered that the biofilm density used when running BM3 in the original iDynoMiCS 1 was incorrectly stated in the publication [17] as an agent biomass density of 15 g L^-1^, but this led to a final biofilm density of ∼ 12 g L^-1^. Hence these simulations were rerun with the modified agent density used in BM3-iD2 to match a biofilm density of 10 g L^-1^. These new results in iDynoMiCS 1 are also presented here.

##### Process Overview and Scheduling

The BM3-iD2 simulation proceeds in global timesteps representing 12 minutes of simulated time. Within this timestep, various core processes are simulated in a set order, while other processes (specifically, data reporting processes) occur less regularly than the global timestep. The order of processes in the spatial domain is as follows:

1. Agent removal – Agents with centers higher than 500 µm above the base of the biofilm are removed
2. Mechanical relaxation – Either shoving or Force-based Mechanical relaxation to minimize agent overlaps
3. Reaction-diffusion – Agents determine their reaction rates, based on solute concentrations and biomass amounts. Active agents also grow and divide. Solute concentration grids are updated according to reaction rates, and the boundary with the bulk compartment is updated (see *Submodels*)
4. Reporting (only every 2 simulated hours) – Biomass density grids and totals of different biomasses are written to files

In the bulk compartment, there is a simpler series of processes as follows:

1. Solute concentrations are updated according to inflows, outflows and diffusion into the biofilm (as determined by the boundary between the two compartments)
2. A file recording solute concentrations is updated.

These two sets of processes are carried out separately within each timestep, with the bulk compartment carrying out its processes before the biofilm compartment.

#### Design Concepts

The majority of the design concepts in BM3-iD2 are identical to the design concepts of the modeling platform itself, iDynoMiCS 2.0. Therefore, for a fuller description of the design concepts, see the Methods section of this paper. Design concepts that are specific to the BM3-iD2 model are described below.

##### Emergence

The interactions between the two species, especially the competition for oxygen and for space, because the top of the biofilm is maintained at a constant height, lead to particular distributions of biomass within the biofilm, which in turn determine the steady state concentrations of COD and ammonium in the bulk liquid.

##### Interaction

Agents interact with one another and with solutes in their local environment. Physical interactions cause agents to push against one another as they grow, causing a flow of actively growing agents and their neighbors upwards towards the top of the biofilm. Consumption of solutes by agents determines the rates of solute diffusion into the biofilm and also facilitates competition between agents, with agents near the top of the biofilm having access to solutes at greater concentrations.

iDynoMiCS 2.0 has two main agent overlap relaxation methods: A Shoving algorithm and Force-based Mechanics, described in detail in *Submodels*. In order to establish whether these different relaxation methods affected the results of BM3, simulations were run with both methods, with agent density adjusted for each method, to achieve an overall biofilm density of 10 g L^-1^.

**Table S7.**
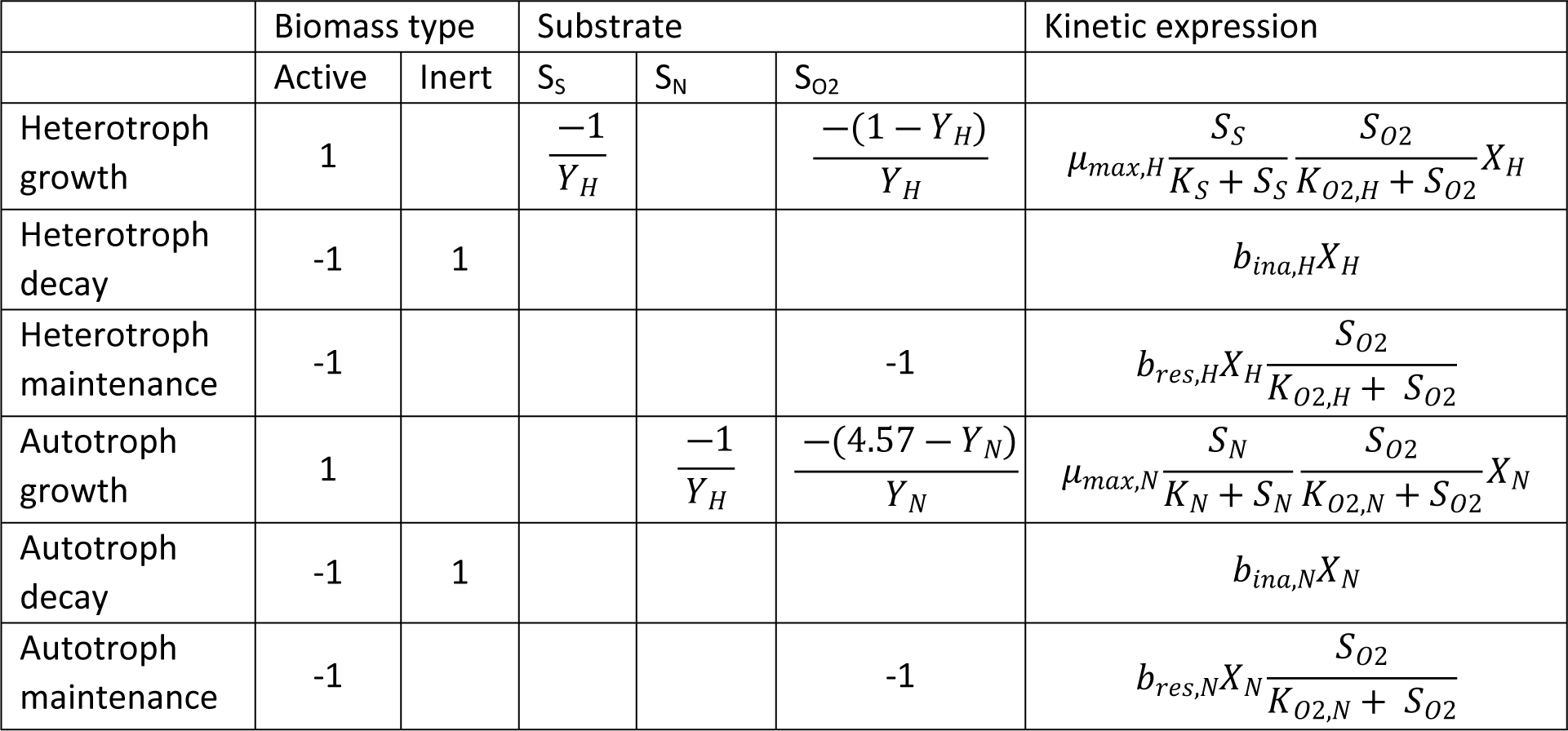
**Petersen (stoichiometric) matrix for reactions in the Benchmark 3 simulations**, adapted from Rittmann [43] and Lardon et al. [17]. Biomasses are denoted with X. Specifically, X_H_ = heterotroph active biomass, X_N_ = nitrifier (autotroph) active biomass. Substrate concentrations are denoted with S, S_S_ for the organic substrate COD, S_N_ for ammonium and S_O2_ for oxygen. For descriptions of the other parameters, see Table S8.

|  | Biomass type |  | Substrate |  |  | Kinetic expression |
| --- | --- | --- | --- | --- | --- | --- |
|  | Active | Inert | S<sub>S</sub> | S<sub>N</sub> | S<sub>O2</sub> |  |
| Heterotroph growth | 1 | | $\frac{-1}{Y_H}$ | | $\frac{-(1 - Y_H)}{Y_H}$ | $\mu_{max,H} \frac{S_S}{K_S + S_S} \frac{S_{O2}}{K_{O2,H} + S_{O2}} X_H$ |
| Heterotroph decay | -1 | 1 | | | | $b_{ina,H} X_H$ |
| Heterotroph maintenance | -1 | | | | -1 | $b_{res,H} X_H \frac{S_{O2}}{K_{O2,H} + S_{O2}}$ |
| Autotroph growth | 1 | | | $\frac{-1}{Y_H}$ | $\frac{-(4.57 - Y_N)}{Y_N}$ | $\mu_{max,N} \frac{S_N}{K_N + S_N} \frac{S_{O2}}{K_{O2,N} + S_{O2}} X_N$ |
| Autotroph decay | -1 | 1 | | | | $b_{ina,N} X_N$ |
| Autotroph maintenance | -1 | | | | -1 | $b_{res,N} X_N \frac{S_{O2}}{K_{O2,N} + S_{O2}}$ |

#### Initialization

50 agents of each species are placed randomly within the bottom 160 µm of the spatial compartment before the first timestep of the simulation. Each of these agents starts with 10 pg active biomass, meaning they are expected to divide in the first timestep as the division mass is 4 pg, introducing some stochastic variation in total agent masses. Initial solute concentrations are set to the values in the bulk inflow.

#### Submodels

##### Bulk solute dynamics

The concentrations of COD and ammonium are solved in the bulk compartment according to Equation 3. However, the concentration of oxygen in the bulk had to be fixed at 10 g m^-3^ to match the results reported for BM3. In the well-mixed region of the spatial compartment that does not contain any agents, concentrations are set to those in the bulk compartment. In the rest of the spatial domain, solutes diffuse through the solute grid and are consumed by agents at rates according to the agent reactions.

**Table S8.**
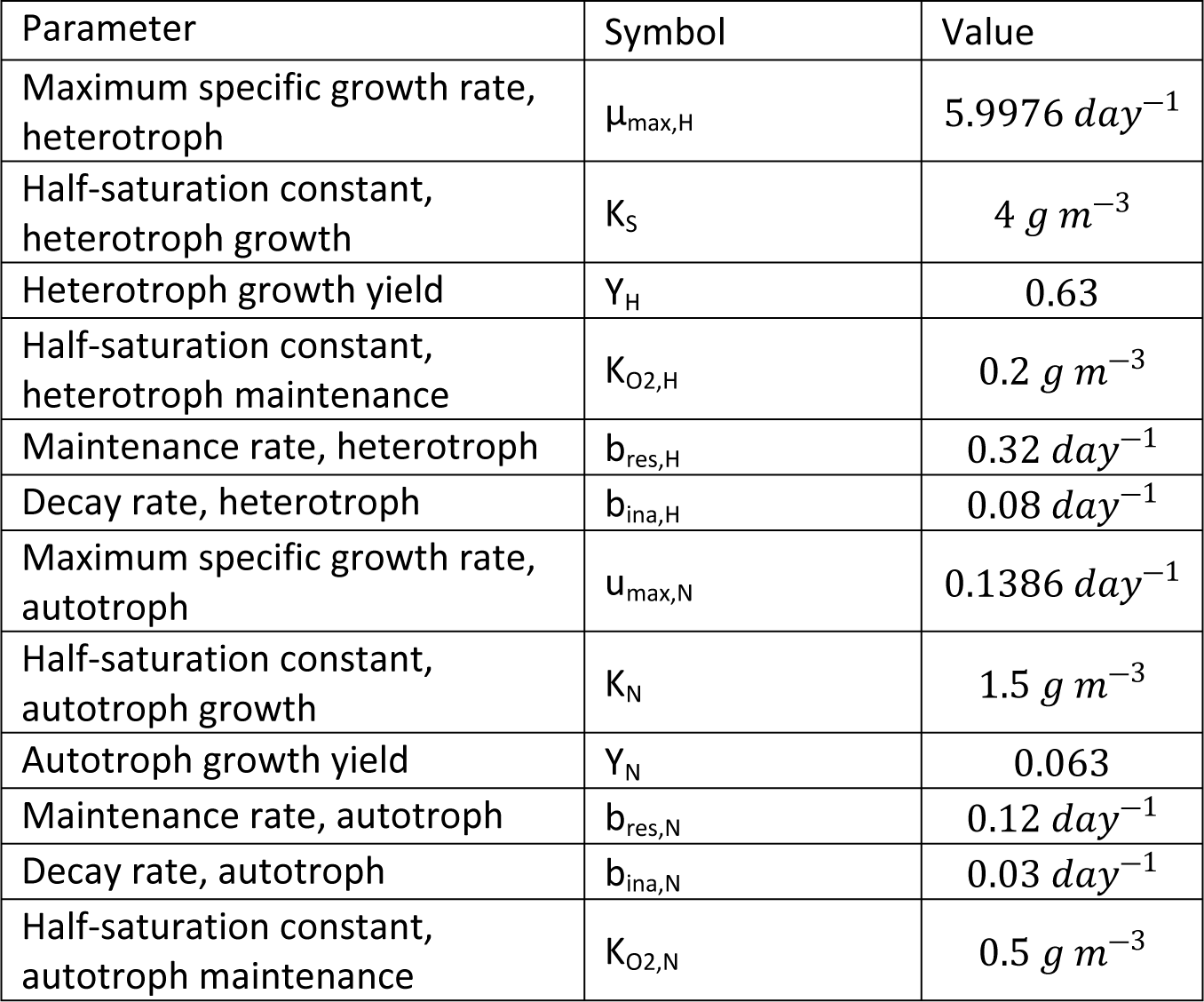
Kinetic parameters in the Benchmark 3 models.

#### Agent Reactions

Both species carry out three different reactions, a metabolism reaction, a maintenance reaction and an inactivation reaction. The metabolism reaction is equivalent to growth, increasing biomass, and using COD as electron donor and oxygen as electron acceptor in the case of heterotrophs and ammonium as electron donor and oxygen as electron acceptor in the case of autotrophs. The maintenance reaction represents the endogenous respiratory consumption of biomass for cell maintenance and oxidizes biomass with oxygen. The inactivation reaction lumps any additional loss of active biomass into a single decay process which converts metabolically active biomass to inert biomass. The growth and respiration reactions proceed according to Monod kinetics, while the inactivation reaction is governed by first order kinetics. See Tables S7 for the Petersen matrix and S8 for the kinetic parameters.

##### BM3-iD2 Results

Once parameters were finalized, three replicates of each combination of case and relaxation method were simulated in iDynoMiCS 2.0 for 120 simulated days. Additionally, three replicates of each case were simulated in iDynoMiCS 1 with the newly adjusted agent density parameter for the purposes of comparison. Solute concentrations generally reached steady state within 20 simulated days, while biomass took longer to reach steady state (Figure S4). To compare iDynoMiCS 2.0 to the other models that have run BM3, various output variables were compared (Figures S4, S5, Table S9). These included steady state concentrations of COD and ammonium, steady state densities of the various biomass forms and the distribution of biomass within the biofilm.

#### Solute Concentrations

Unsurprisingly, the BM3 results from iDynoMiCS 1 and iDynoMiCS 2.0 were very similar (Figure S4). These models have very similar basic designs and in a simple biofilm model, behave very similarly. Furthermore, there is no clear impact of the biomass spreading mechanism used in iDynoMiCS 2.0 as the Shoving or Force-based Mechanics simulation results were very similar. However, different agent densities were required for these two spreading methods to produce an overall biofilm density of 10 g L^-1^, because the FbM produced denser biofilms than the Shoving algorithm in the absence of this adjustment. This is because Shoving is generally used to model the effect of EPS production – increased distance between cells – implicitly by using a Shoving factor, a multiplier on the radius of the cells. There is no EPS in BM3, including EPS would reduce biofilm density in a mechanistic way.

Steady state bulk liquid concentrations from the results with Shoving were compared with the results from the other models using the multivariate version of the t-test, Hotelling’s 1-sample T^2^ test. In the case of results from simulations with the Shoving relaxation algorithm, results from iDynoMiCS 2.0 did not differ significantly from the distribution of the other models (Table S9). Despite this, the steady state COD concentrations in iDynoMiCS 1 and 2.0 were generally higher than those in the other models (Figure S4, Table S9). In simulations using Force-based mechanics, there was a significant difference between iDynoMiCS 2.0 and the other models in the high ammonium case (Figure S9). This suggests that using force-based mechanics leads to a more pronounced difference in the BM3 results, due to differences in the spreading and distribution of biomass.

#### Biomass distribution

Another key output of the BM3-iD2 model is the biomass density and vertical distribution. As both species in the model can have both active and inert biomass, there are three different biomass types with different concentrations and distributions - heterotrophic, autotrophic and inert. The total areal densities of these biomass types are compared between models in Table S10. Vertical distributions of the various biomass types in the different cases are shown in Figure S6. These show a qualitatively similar pattern to the CP model [44], with fast-growing heterotrophs dominating the top of the biofilm, while autotrophs grow more slowly and are at their most abundant in the middle or bottom of the biofilm. Autotrophs vary widely in abundance between the different cases, being at very low numbers in the low ammonium case. This is to be expected, given that their energy source is at a low concentration. In all three cases, the bottom of the biofilm is dominated by inert biomass due to the lower substrate concentrations at the bottom of the biofilm reducing growth relative to maintenance and inactivation. This is most pronounced in the low ammonium case, which has the highest proportion of inert biomass of the three cases.

Given the differences in modeling approaches of the various IWA task group models, one might expect iDynoMiCS 1 and iDynoMiCS 2.0 to produce results closer to the NUFEB and CP models than to any of the other IWA task group models. Although, the NUFEB results were close, differences with CP are larger. In fact, the results from the W platform were the closest match in steady state solute concentrations, while those from the M1 platform were the closest match for overall biomass densities. This is particularly interesting given that these are both 1-dimensional platforms utilizing the AQUASIM software rather than agent-based. It is possible that the similarity derives from a closer match in biomass distribution than with the other models due to less stochastic mixing of the biomass. However, as the biomass distributions were not published for these models, this is difficult to determine. Hotellings 1-sample T^2^ tests were carried out to compare the areal biomass densities in iDynoMiCS 2.0 to that in the IWA models. The results from iDynoMiCS 2.0 do not differ significantly from the set of results from the IWA models.

**Fig S4.**
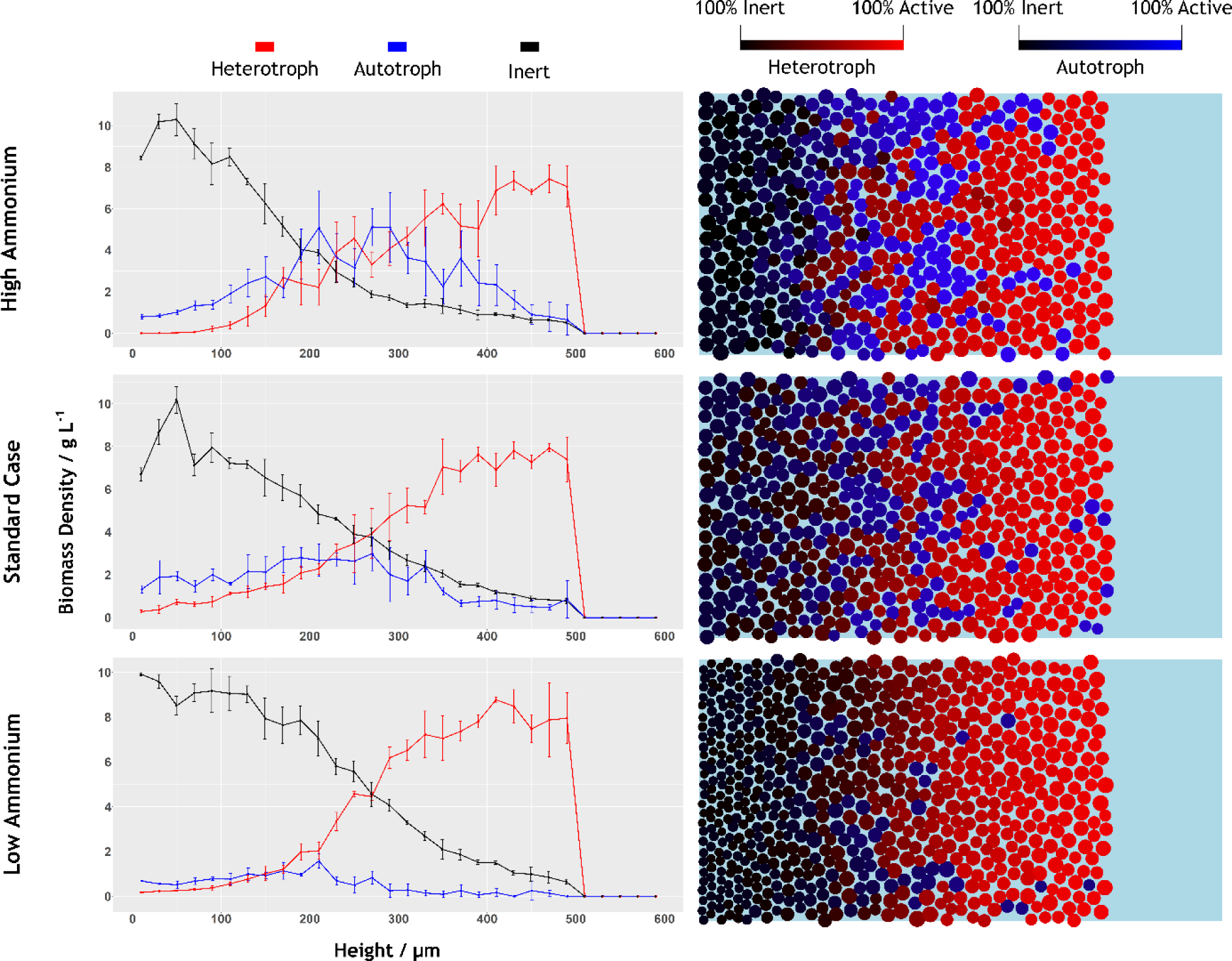
BM3-iD2 biomass distribution in the three cases. Left column shows the average areal density of each biomass type. Error bars show the standard deviation based on the three replicates run for each case. Right column shows an example from the final timestep of one replicate for each case. Coloring of agents shows the proportion of biomass that is active or inert.

**Fig S5.**
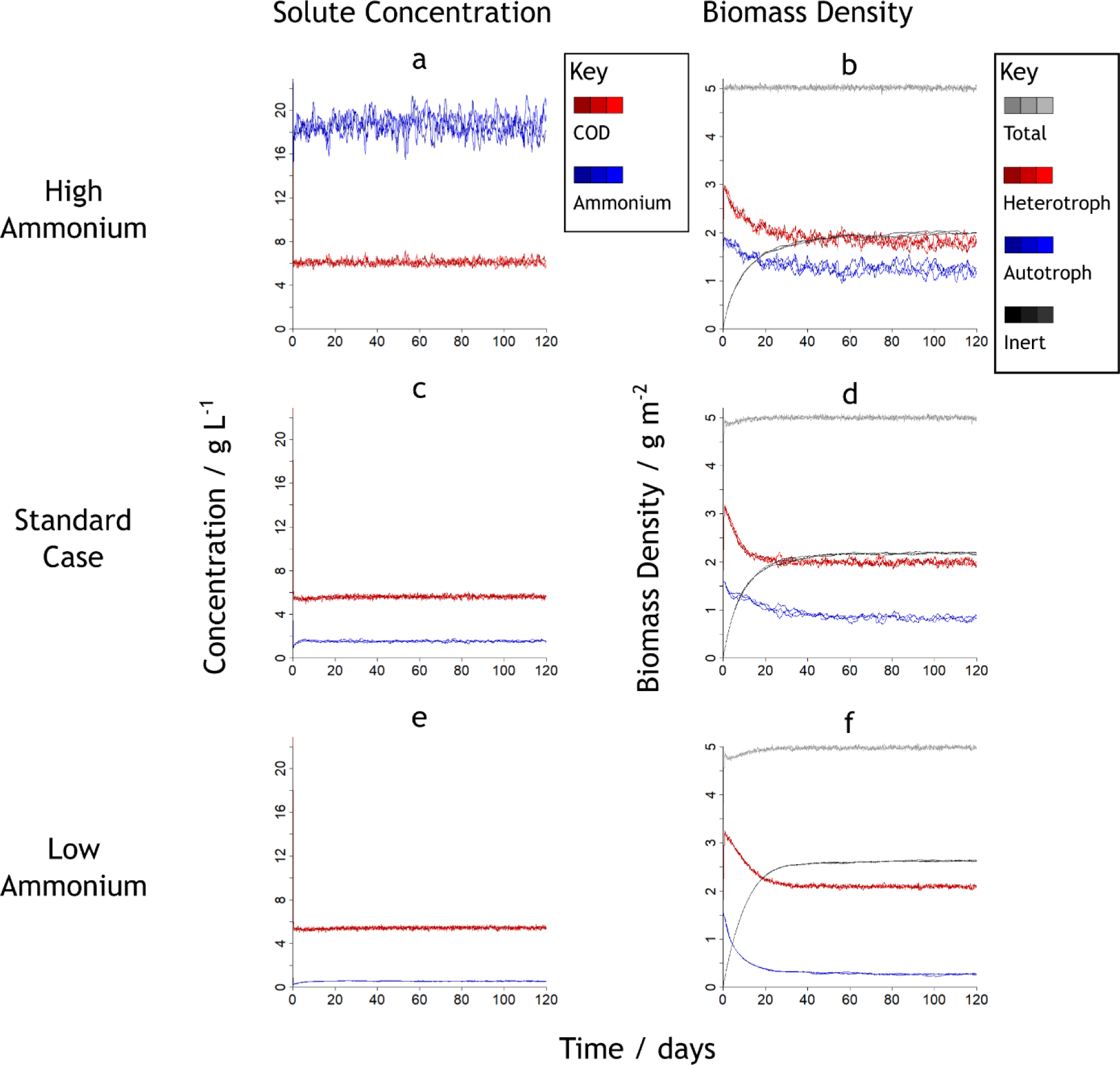
BM3-iD2 solute concentrations and areal biomass densities over time in the three simulated cases. Lines of different shades of the same color represent different replicates of the simulations run with different random number seeds. Three replicates were run for each case. Results are from the simulations with the Shoving biomass spreading algorithm.

**Table S9.** Steady state substrate concentrations in the various IWA task group models and in iDynoMiCS 1 and iDynoMiCS 2.0. Results for the latter models were averaged over the stochastic steady states. Hotelling’s T^2^ tests were performed to compare the results from iDynoMiCS 2.0 to those from all other models, including the IWA models, NUFEB and iDynoMiCS 1.

|  |  | High ammonium |  | Standard case |  | Low ammonium |  |
| --- | --- | --- | --- | --- | --- | --- | --- |
|  |  | COD | Ammonium | COD | Ammonium | COD | Ammonium |
| IWA models | CP | 5.45 | 18.15 | 5.14 | 1.50 | 4.39 | 0.44 |
|  | DN | 5.56 | 20.26 | 5.14 | 1.74 | 4.98 | 0.48 |
|  | W | 5.86 | 18.93 | 5.39 | 1.59 | 5.19 | 0.48 |
|  | M1 | 5.35 | 17.03 | 4.84 | 1.45 | 4.66 | 0.45 |
| NUFEB |  | 5.74 | 18.42 | 5.21 | 1.72 | 5.18 | 0.53 |
| iDynoMiCS models | iDynoMiCS 1 | 6.08 | 18.58 | 5.63 | 1.55 | 5.45 | 0.55 |
|  | **iDynoMiCS 2.0 (Shoving)** | 6.11 | 18.68 | 5.62 | 1.54 | 5.43 | 0.54 |
| | Hotelling's $T^2$<br>Test p-value | 0.0548 | | 0.0523 | | 0.1306 | |
|  | **iDynoMiCS 2.0 (FbM)** | 6.13 | 18.50 | 5.60 | 1.55 | 5.41 | 0.55 |
| | Hotelling's $T^2$<br>Test p-value | **0.0431** | | 0.0646 | | 0.0765 | |

**Table S10.**
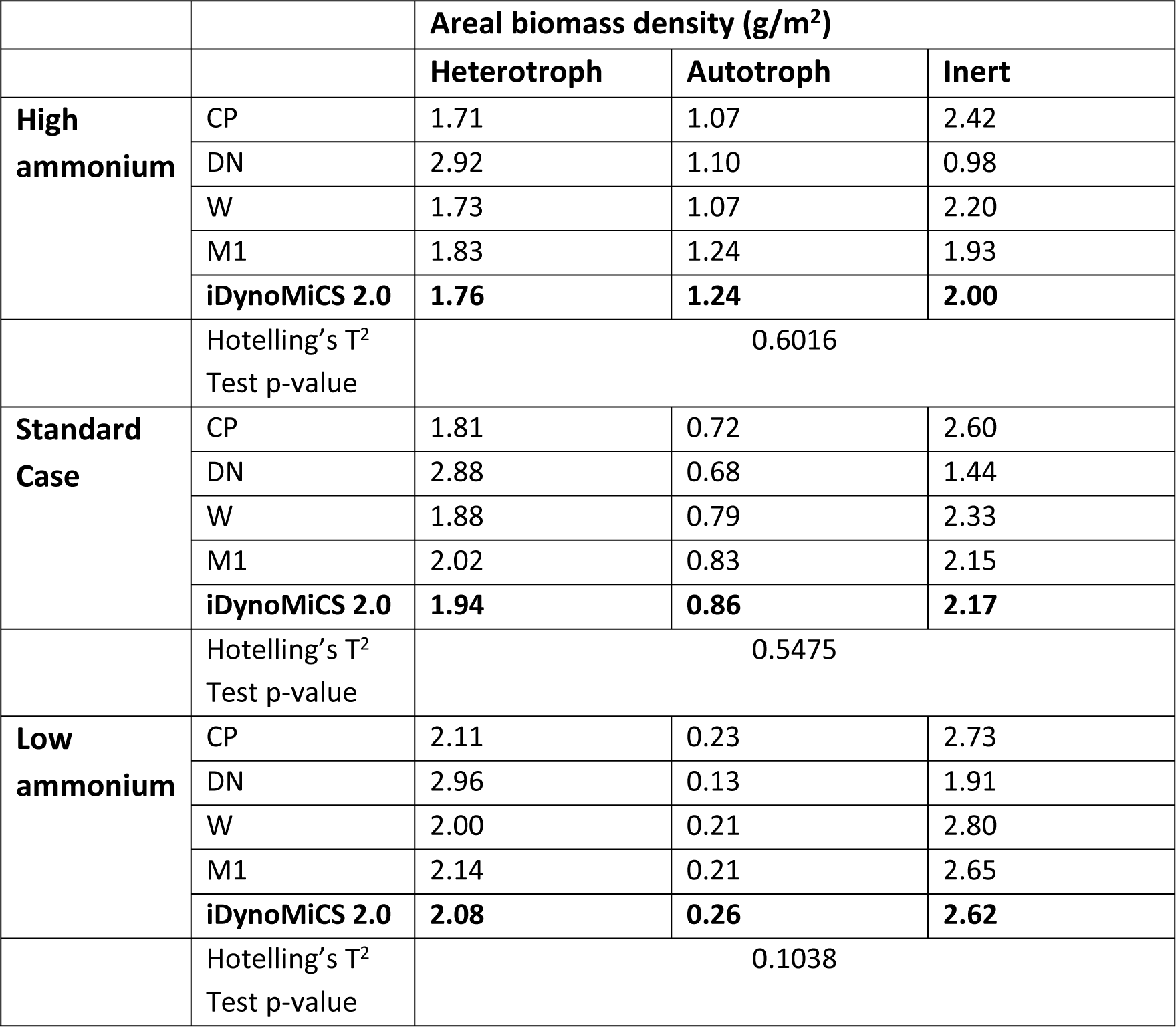
**Steady state areal biomass density** (mass per unit surface area) of different types of biomass in the biofilm. The iDynoMiCS 2.0 results are from simulations using the Shoving algorithm. Hotelling’s T^2^ tests were performed to compare the results from iDynoMiCS 2.0 to those from the IWA models. Biomass density was not reported on the NUFEB model benchmark, and it is thus not included in this comparison.

### S1.6 Supplementary Information for “Comparing the effect of different biomass spreading mechanisms: Biofilms promote altruism case study”

**Table S11.**
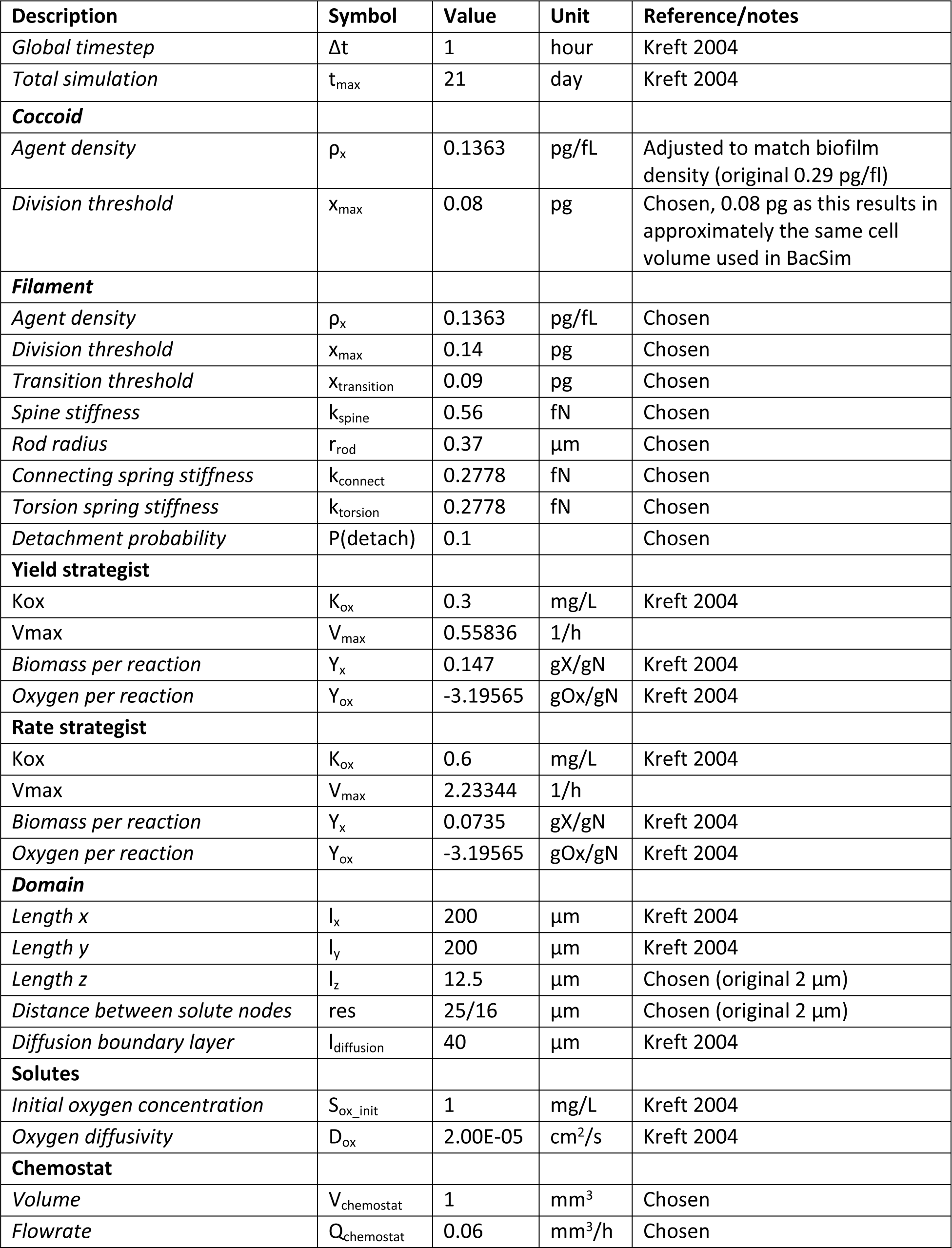
Model parameters for 3D simulations of the “biofilms promote altruism” case study.

Shoving as used in BacSim results in more open space between agents compared to mechanical relaxation as used in iDynoMiCS 2.0, which by default only resolves overlap between agents. To compensate for this effect, we have reduced agent density in iDynoMiCS 2.0 by 53%, such that overall biofilm densities remain similar. In order to compare biofilm densities, the computational domain was split into 100×100 bins. Each bin receives a number equal to the total mass of all agents with their center of gravity in the bin, division by the bin volume reveals the local biofilm density (Fig S6).

For both the iDynoMiCS 2.0 and BacSim simulations, inspection of biofilm density at different heights revealed that the majority of bin rows were not significantly different in terms of density in comparison to the total amount of filled bins. For both platforms we observed a significant drop-off of biofilm density at the top surface where bins may be partially filled and the expansion front may result in a sparser local agent population.

In BacSim simulations we observe a significant biofilm density drop in the first row of bins (at the base of the biofilm). This may be a result of how the BacSim shoving algorithm resolves interactions with a hard surface in combination with less efficient sphere packing at a flat surface. No significant biofilm density drop was observed at the base of the iDynoMiCS 2.0 simulations. With both platforms, occasional small but significant peaks or drops in biofilm density (2.5 < P < 5.0) are observed in some bins. These occasional drops and peaks are likely artifacts as a result of a spatial aliasing effect between the binning resolution and local sphere packing.

With both platforms, density bins, excluding bins at the biofilm extremes, follow a normal distribution. After the before mentioned agent density adjustment, no significant difference in the overall biofilm density is observed between simulations of the two platforms. The standard deviation of density bins is higher in BacSim simulations. This can be explained by the difference in agent properties, maximum agent size is kept the same in iDynoMiCS 2.0, translating into a higher overall amount of agents with a lower mass, resulting in less bin-to-bin mass fluctuation.

**Fig S6.**
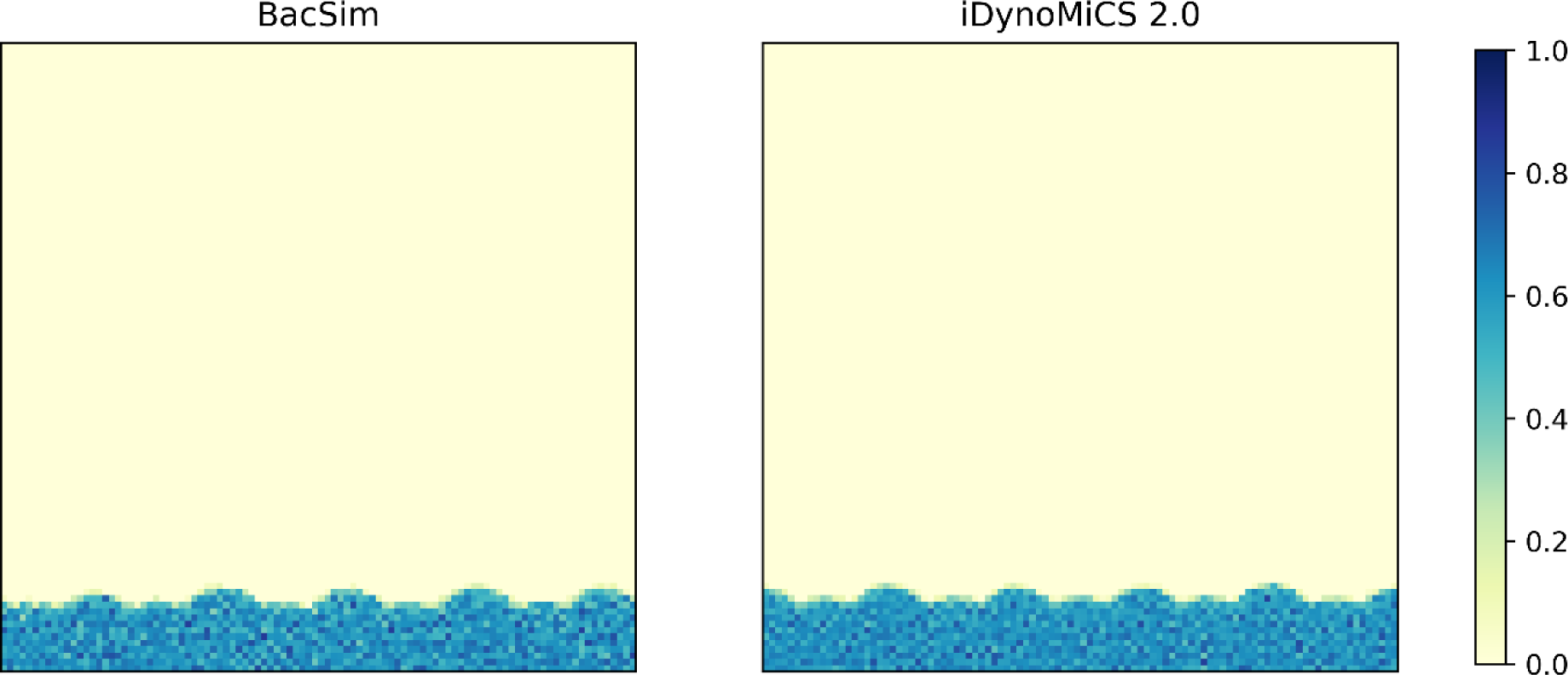
Comparison of agent density (pg/µm^3^) distributions in biofilms simulated in BacSim using the shoving algorithm vs iDynoMiCS 2.0 using FbM. The panels correspond to Fig 5A (left) and B (right). The computational domain was split into 100×100 grid elements, each received the full mass of agents whose center of gravity was inside the grid element, division by the grid element volume gave the local biofilm density (pg/µm^3^). iDynoMiCS 2.0 agent density was reduced by 53% in order to achieve a similar overall biofilm density. With BacSim the biofilm density at the base was observed to be significantly lower than in the rest of the biofilm. The BacSim simulations further showed a higher standard deviation amongst bins, due to the higher agent density in these simulations.

**Fig S7.**
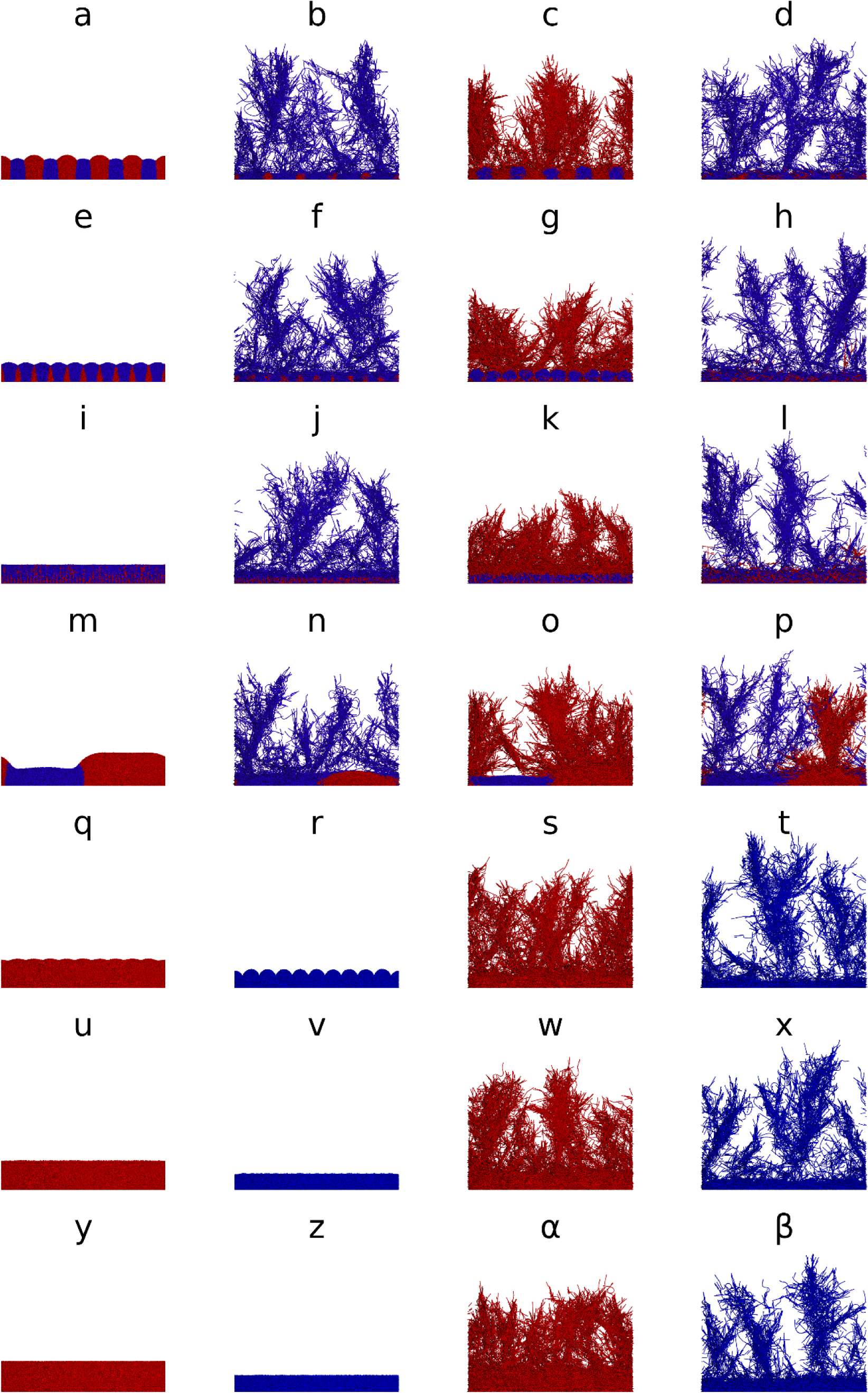
Replicates of biofilms promote altruism case study simulations shown in. Fig. 6.

### S1.7 Model initiation

#### Box S1. Example of a simple iDynoMiCS 2.0 protocol file used to specify a particular model to be read and executed by the platform.

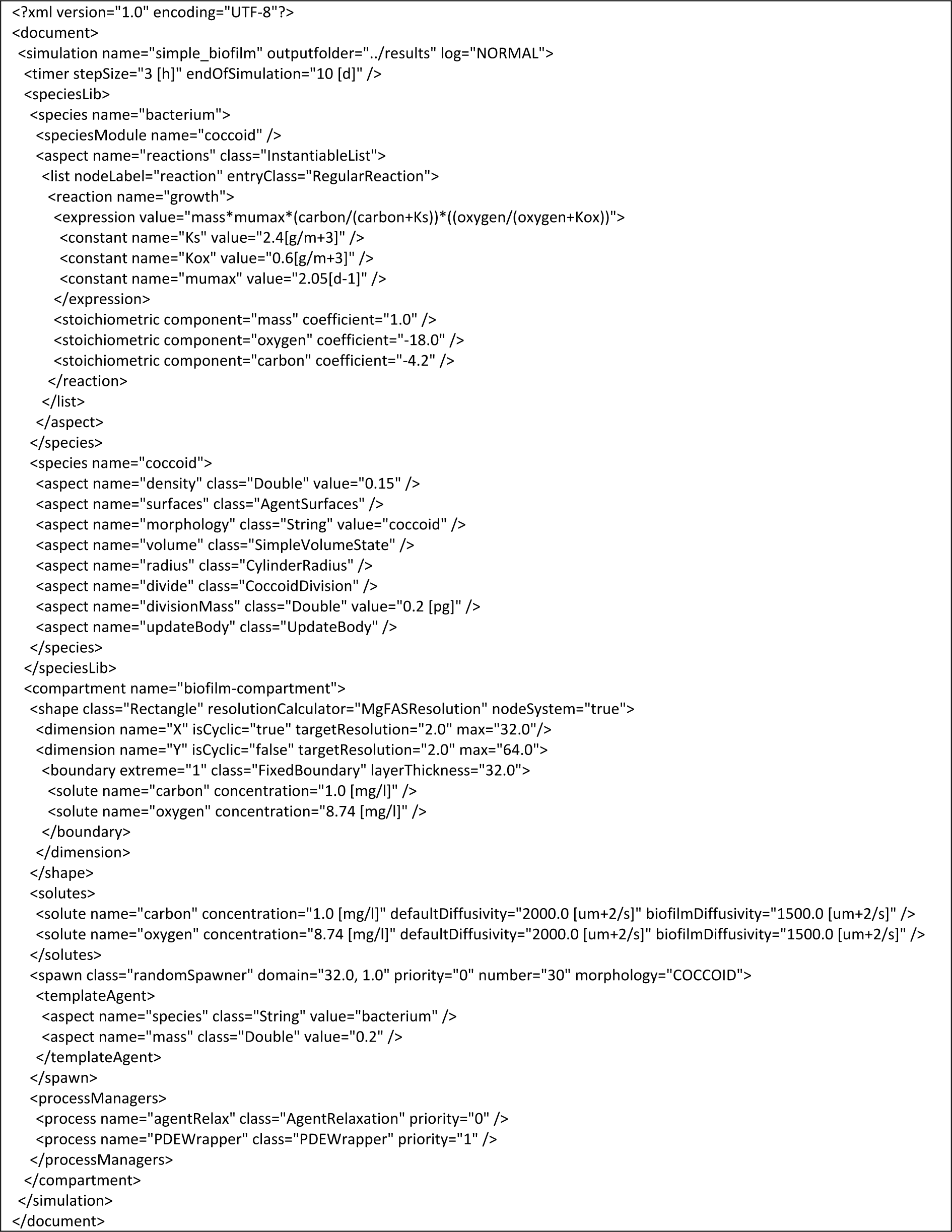

### S1.8 Software structure

#### Box S2. This example shows how with a few lines of code a new aspect class can be created.

In this case it is a class that calculates a coccoid radius from its volume. Because here the abstract super class “Calculated” is extended, the newly written class integrates seamlessly in the framework as initialization and data handling is handled automatically.

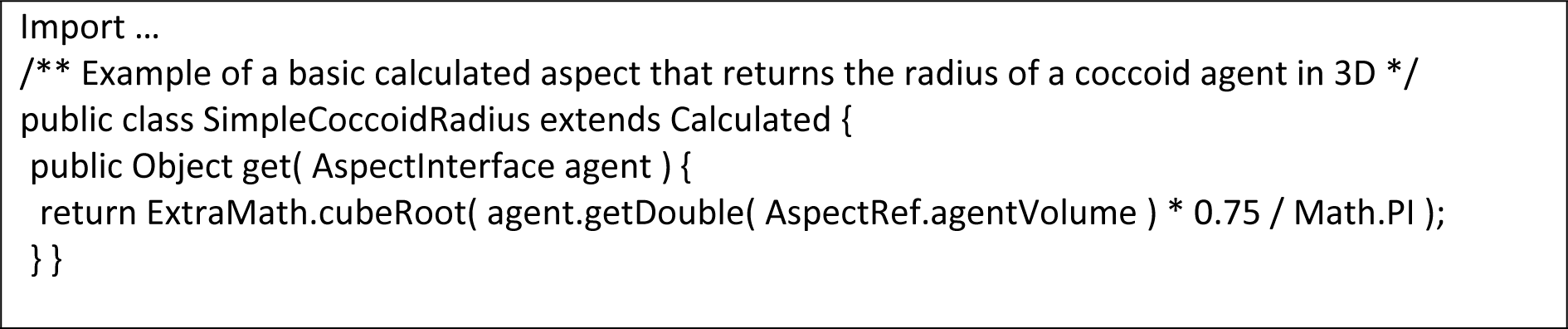

### S1.9 Included test scenarios

**Table S12.** A selection of test protocols that are included with iDynoMiCS 2.0.

| Title | File | Description |
| --- | --- | --- |
| Simple | simple.xml | Minimalistic protocol file testing basic functionality with default parameters. |
| Chemostat | chemostat.xml | A basic chemostat setup with a reaction occurring in the environment (non-agent mediated). |
| Fed-batch | fedbatch.xml | Basic fedbatch scenario, agents grow in a non-spatial compartment with constant “feed” increasing volume and supplying solutes. |
| Sensing | simple_sensing.xml | Basic scenario with local solute sensing, agents “differentiate” when the signal surpasses a threshold concentration. |
| Conditional colouring | conditional_colouring.xml | A basic nitrifying biofilm with conditional coloring, agents receive a color gradient based on the amount of internally stored eps. |
| Bacilli | bacilli.xml | A basic setup with 2 colonies of rod shaped agents merging. |
| Plasmid spatial | plasmid.xml | A basic setup testing plasmid transfer in a spatial compartment. |
| Plasmid chemostat | plasmid_chemostat.xml | A basic setup testing plasmid transfer in a well-mixed environment. |
| Stress test | stress_test_7c.xml | A large scale nitrifying biofilm used to test the limits of iDynoMiCS 2.0. |
| Benchmark 3 | /BM3/<br>(6 files) | A set of model scenarios corresponding the 3 different ammonium concentrations as analog to the benchmark 3 cases using FbM or shoving. |
| Altruism 2.5D | c10_ld_fb.xml,<br>d10_ld_fb.xml,<br>e10_ld_fb.xml | A set of model scenarios corresponding to those found in “Biofilms promote altruism” Kreft 2004, Fig 4. scenario c, d and e. |
| Altruism 3D | /altruism 3D/<br>(28 files) | A set of model scenarios corresponding to scenarios presented in Fig 6. |
| Reaction diffusion tests | /reaction<br>diffusion/<br>(4 files) | A set of model scenarios corresponding to the reaction and diffusion tests as presented in S1.3 |
| Unit tests | /unit-tests/<br>(7 files) | A set of protocols used for automated software and solver testing. |

### S1.10 The graphical user interface

**Fig S8.**
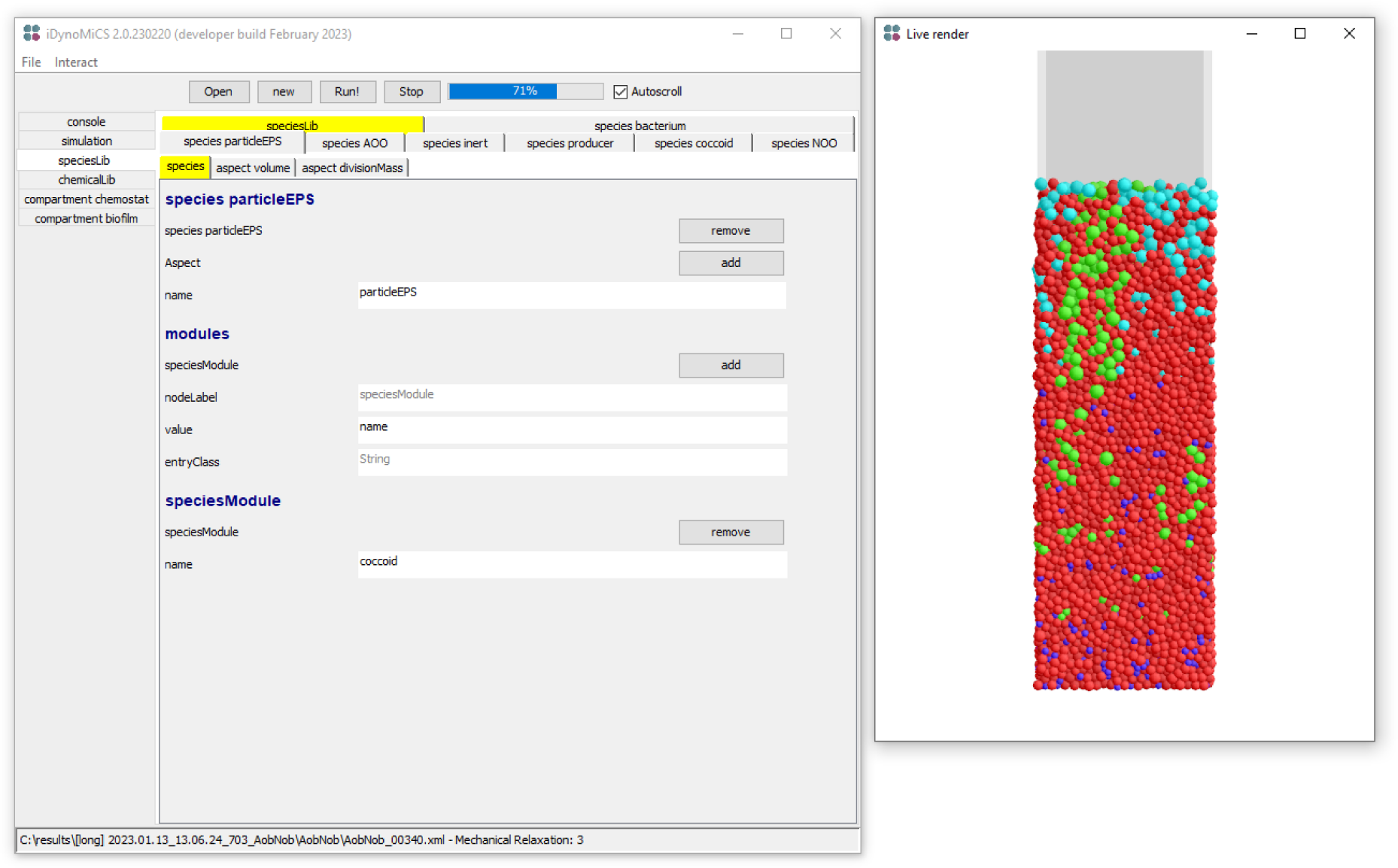
Preview of the iDynoMiCS 2.0 GUI during simulation. The GUI may be used to review, edit or create protocol files before running them. During the simulation, the simulation state may be viewed but no longer edited (left). The GUI can further provide useful feedback including key information such as substrate concentration, species abundance, convergence of the reaction diffusion solver, etc. through the console. Spatial compartments can be rendered directly to provide insights in agent species distribution and concentration gradients (right). Lastly the GUI can be used to extract key data from iDynoMiCS 2.0 output files, convert between EXI and XML files and convert numbers between different unit systems including SI and iDynoMiCS 2.0 base units.

### S1.11 Plasmid Dynamics

For plasmid dynamics, two distinct processes were implemented: conjugative transfer from donor to recipient cells and loss of the plasmid due to segregation. Plasmid loss can only happen upon cell division and hence was encoded as a probability. Conjugation was considered pili-driven, with a maximum length pili can reach before they start retracting. The dynamics of transfer were incorporated based on the live cell imaging by Clarke *et al.* (2008) [97].

From the imaging, certain aspects of F-pilus extension and retraction can be observed: During extension, the filament elongates from the base. A fully extended 4-μm pilus retracts completely. F-pili on the same cell are independently regulated, with about three pili growing and retracting asynchronously. The average time required for extension and retraction of the pili informed the extension and retraction speeds of the pili used in the plasmid dynamics process manager.

In the model, the conjugation process begins with pili extension, assuming pili extend in all directions from the cell surface. On encountering a recipient, the pilus tries to attach to its surface and if a pilus attaches to a recipient cell, all pili start retracting. Certain pili are capable of transferring the plasmid without pili retraction, but others transfer the plasmid only upon cell surface to surface contact, with the pilus only bringing the cells together by retracting with the recipient cell attached.

Like in iDynoMiCS 1, plasmid carrying cells search the neighborhood within the reach of the pilus. A difference arises in the method implemented for this search. Instead of the “scan speed” in iDynoMiCS 1, the required parameters are maximum pilus length and “transfer probability”. Using the F-pili data from live cell imaging by Clarke *et al.* (2008), the pilus length can be calculated for each time step of the process as a function of extension speed. The current length is used as the maximum distance for neighborhood search. The closest neighbors are prioritized by increasing the neighbor search range in increments of 0.01 µm until the current pilus length is reached or a neighbor is found. As an example, with pilus length of 3.2 µm, a neighborhood search will be performed 320 times with the distance searched increased from 0.01 to 3.2 in increments of 0.01 µm. The search will be terminated early if a plasmid-free neighbor is found.

Once a plasmid free neighbor is found, the transfer event happens instantly with a success probability given by the parameter “transfer probability”. Biologically, the transfer happens after pilus retraction and then the plasmid goes into a “cool down” period. To reduce the computational requirement for agent movements, the time for retraction of the longest pilus is added to the cool down period as a wait time between plasmid transfer attempts.

**Table S13.** Parameters required for the plasmid dynamics process manager in iDynoMiCS 2.0.

| Parameter | Definition |
| --- | --- |
| Transfer probability | Governs the success of plasmid transfer; user must provide either one of the parameters with the relation being:<br>Transfer probability = frequency × number of neighbors |
| Transfer frequency |  |
| Loss probability | Governs the success of loss event upon cell division |
| Pilus length | Maximum length of pilus extension from donor cell surface. If plasmid transfer is not pilus driven, this can be set to 0 |
| Aspects to Transfer | Agent aspects to change on plasmid acquisition or loss (MIC, Fitness cost, etc.) |
| Cool down | Rest time between plasmid transfers |
| Extension speed | Extension speed for pilus related to this plasmid |
| Retraction speed | Retraction speed for pilus related to this plasmid. The time for complete retraction is added to the cool down time before the next transfer can start |

Since each plasmid is a separate aspect specified in the protocol file, there can be multiple plasmids included. These will implicitly be considered compatible with each other as plasmid incompatibility is not currently implemented.

Plasmid loss due to segregation is defined as an event in the code, thus requiring inclusion as an aspect in the protocol file. However, the event is triggered upon cell division only if the “loss probability” parameter is defined in the included plasmid dynamics process manager. Thus, upon cell division, the daughter cells can retain the plasmid or one can lose it at the probability defined in the protocol file.

For chemostats, the transfer process is governed by the following equation to determine the number of transfers for each agent with plasmid (donor):

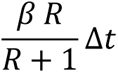

Where β is the transfer frequency, R is the number of plasmid-free agents (recipients) in the population and Δt is the time step size. This equation is calculated for each donor so implicitly, the rate is proportional to donor concentration. It is analogous to the infection rate in Susceptible-Infectious-Recovered compartmental epidemiological models, where recipients are considered Susceptible and plasmid transfer is analogous to the process of infection. The recipients for the plasmid are selected randomly from the whole population as a chemostat is considered well-mixed.

### S1.12 Agent density scaling for 2D simulations

Due to the virtual third dimension of 1 µm in 2D simulations, cell radii and/or lengths can differ between 2D and 3D compartments. iDynoMiCS 2.0 can scale agent densities, such that the dimensions of agents in 2D match what they would be in a 3D environment.

Users define an actual 3D density, ρ_3D_, which is then used to calculate a scaled density in the 2D compartment, ρ_2D_. The exact calculation depends on the shape of the agent or filament section in question.

For spherical (coccoid) agents, the radius of a sphere is calculated based on the agent’s mass and actual density:

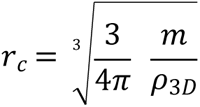

Where *r*_*c*_is the radius and m the agent’s total mass. The scaled density is thus given by:

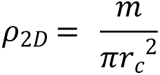

For agents or filament elements with a rod (bacillus) shape, it is the length, rather than the radius, that must be calculated. The 3D length of the line-segment connecting the agent’s points is given by:

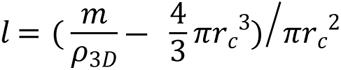

And the 2D scaled density is given by:

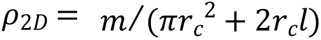

### S1.13 Microbial IbM publications on PubMed

**Fig S9.**
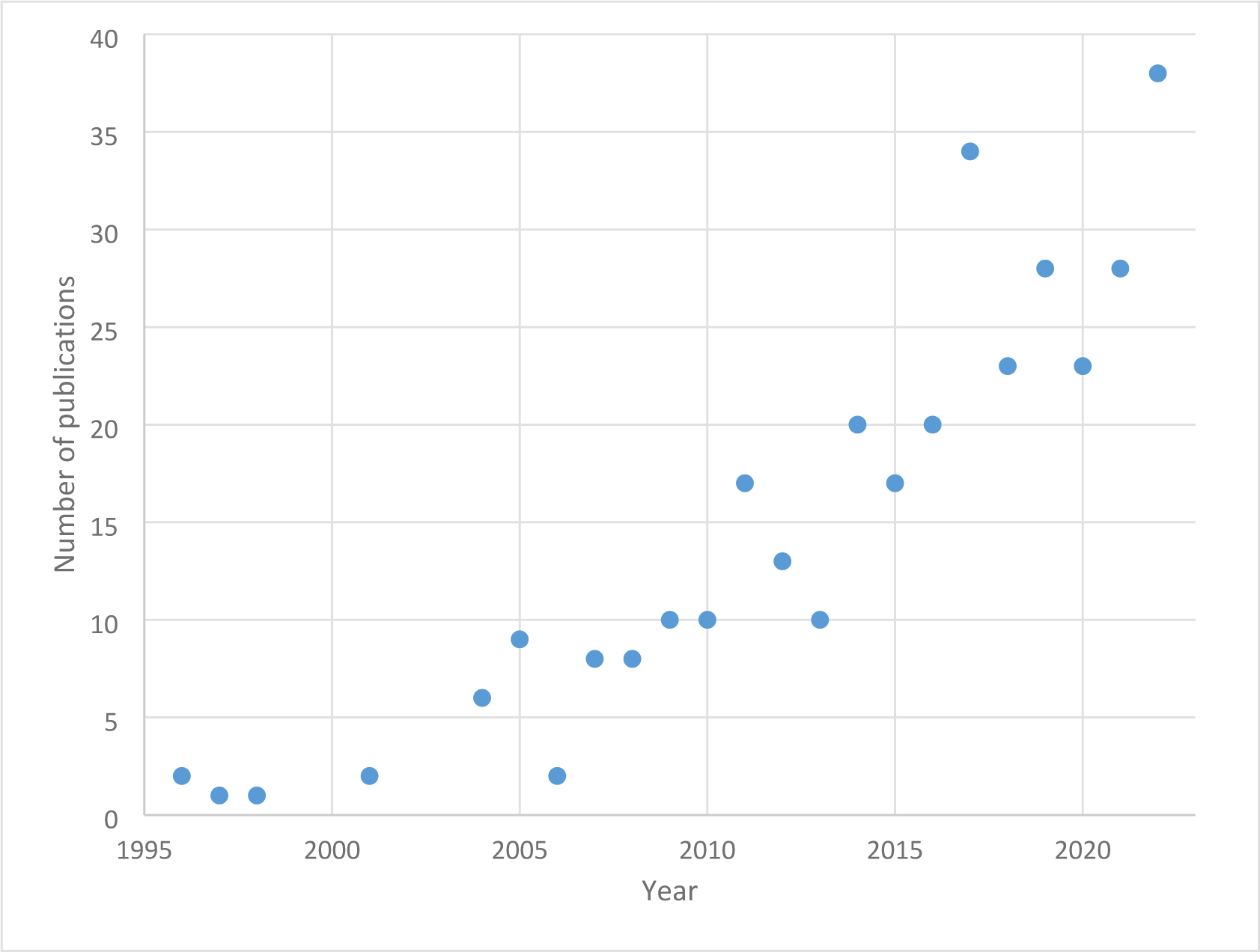
The number of publications on microbial IbM on PubMed since 1995. A simple search query on PubMed reveals a growing trend in applying IbM to microorganisms. The following query was used: “((biofilm) OR (microbial)) AND ((individual-based) OR (agent-based)) AND (model)”.

## References

1. Flemming H-C, Wuertz S. Bacteria and archaea on Earth and their abundance in biofilms. Nat Rev Microbiol. 2019;17: 247–260. doi:10.1038/s41579-019-0158-9

2. Widder S, Allen RJ, Pfeiffer T, Curtis TP, Wiuf C, Sloan WT, et al. Challenges in microbial ecology: building predictive understanding of community function and dynamics. ISME J. 2016;10: 2557– 2568. doi:10.1038/ismej.2016.45

3. Parsek MR, Fuqua C. Biofilms 2003: Emerging Themes and Challenges in Studies of Surface-Associated Microbial Life. J Bacteriol. 2004;186: 4427–4440. doi:10.1128/JB.186.14.4427-4440.2004

4. Becker P, Hufnagle W, Peters G, Herrmann M. Detection of Differential Gene Expression in Biofilm-Forming versus Planktonic Populations of Staphylococcus aureus Using Micro-Representational-Difference Analysis. Appl Environ Microbiol. 2001;67: 2958–2965. doi:10.1128/AEM.67.7.2958-2965.2001

5. Ackermann M. Microbial individuality in the natural environment. ISME J. 2013;7: 465–467. doi:10.1038/ismej.2012.131

6. Zimmermann M, Escrig S, Hübschmann T, Kirf MK, Brand A, Inglis RF, et al. Phenotypic heterogeneity in metabolic traits among single cells of a rare bacterial species in its natural environment quantified with a combination of flow cell sorting and NanoSIMS. Front Microbiol. 2015;06. doi:10.3389/fmicb.2015.00243

7. Wanner O, Eberl HJ, Morgenroth E, Noguera DR, Picioreanu C, Rittmann BE, et al. Mathematical modeling of biofilms. London: IWA Publishing; 2006.

8. Kissel JC, McCarty PL, Street RL. Numerical Simulation of Mixed-Culture Biofilm. J Environ Eng. 1984;110: 393–411. doi:10.1061/(ASCE)0733-9372(1984)110:2(393)

9. Wanner O, Gujer W. A multispecies biofilm model. Biotechnol Bioeng. 1986;28: 314–328. doi:10.1002/bit.260280304

10. Rittmann BE, Manem JA. Development and experimental evaluation of a steady state, multispecies biofilm model. Biotechnol Bioeng. 1992;39: 914–922.

11. Dockery J, Klapper I. Finger formation in biofilm layers. SIAM J Appl Math. 2001;62: 853–869.

12. Picioreanu C, van Loosdrecht MCM, Heijnen JJ. Mathematical modeling of biofilm structure with a hybrid differential-discrete cellular automaton approach. Biotechnol Bioeng. 1998;58: 101–116. doi:10.1002/(SICI)1097-0290(19980405)58:1<101::AID-BIT11>3.0.CO;2-M

13. van Loosdrecht MCM, Heijnen JJ, Eberl HJ, Kreft JU, Picioreanu C. Mathematical modelling of biofilm structures. Antonie Van Leeuwenhoek Int J Gen Mol Microbiol. 2002;81: 245–256. doi:10.1023/A:1020527020464

14. Hartmann R, Singh PK, Pearce P, Mok R, Song B, Díaz-Pascual F, et al. Emergence of three-dimensional order and structure in growing biofilms. Nat Phys. 2019;15: 251–256. doi:10.1038/s41567-018-0356-9

15. Kreft J-U, Picioreanu C, Wimpenny JWT, van Loosdrecht MCM. Individual-based modelling of biofilms. Microbiology. 2001;147: 2897–2912. doi:10.1099/00221287-147-11-2897

16. Hellweger FL, Fredrick ND, McCarthy MJ, Gardner WS, Wilhelm SW, Paerl HW. Dynamic, mechanistic, molecular-level modeling of cyanobacteria: *Anabaena* and nitrogen interaction. Environ Microbiol. 2016;18: 2721–2731. doi:10.1111/1462-2920.13299

17. Lardon LA, Merkey BV, Martins S, Dötsch A, Picioreanu C, Kreft J-U, et al. iDynoMiCS: next-generation individual-based modelling of biofilms. Environ Microbiol. 2011;13: 2416–2434. doi:10.1111/j.1462-2920.2011.02414.x

18. Gorochowski TE, Matyjaszkiewicz A, Todd T, Oak N, Kowalska K, Reid S, et al. BSim: an agent-based tool for modeling bacterial populations in systems and synthetic biology. PLoS ONE. 2012;7: e42790. doi:10.1371/journal.pone.0042790

19. Jang SS, Oishi KT, Egbert RG, Klavins E. Specification and Simulation of Synthetic Multicelled Behaviors. ACS Synth Biol. 2012;1: 365–374. doi:10.1021/sb300034m

20. Goñi-Moreno A, Amos M. DiSCUS: A Simulation Platform for Conjugation Computing. In: Calude CS, Dinneen MJ, editors. Unconventional Computation and Natural Computation. Cham: Springer International Publishing; 2015. pp. 181–191. doi:10.1007/978-3-319-21819-9_13

21. Li B, Taniguchi D, Gedara JP, Gogulancea V, Gonzalez-Cabaleiro R, Chen J, et al. NUFEB: A massively parallel simulator for individual-based modelling of microbial communities. Darling AE, editor. PLOS Comput Biol. 2019;15: e1007125. doi:10.1371/journal.pcbi.1007125

22. Breitwieser L, Hesam A, De Montigny J, Vavourakis V, Iosif A, Jennings J, et al. BioDynaMo: a modular platform for high-performance agent-based simulation. Wren J, editor. Bioinformatics. 2022;38: 453–460. doi:10.1093/bioinformatics/btab649

23. Bogdanowski A, Banitz T, Muhsal LK, Kost C, Frank K. McComedy: A user-friendly tool for next-generation individual-based modeling of microbial consumer-resource systems. Harcombe WR, editor. PLOS Comput Biol. 2022;18: e1009777. doi:10.1371/journal.pcbi.1009777

24. Grimm V, Berger U, Bastiansen F, Eliassen S, Ginot V, Giske J, et al. A standard protocol for describing individual-based and agent-based models. Ecol Model. 2006;198: 115–126. doi:10.1016/j.ecolmodel.2006.04.023

25. Grimm V, Berger U, DeAngelis DL, Polhill J, Giske J, Railsback SF. The ODD protocol: a review and first update. Ecol Model. 2010;221: 2760–2768. doi:10.1016/j.ecolmodel.2010.08.019

26. Trunk T, S. Khalil H, C. Leo J, Bacterial Cell Surface Group, Section for Genetics and Evolutionary Biology, Department of Biosciences, University of Oslo, Oslo, Norway. Bacterial autoaggregation. AIMS Microbiol. 2018;4: 140–164. doi:10.3934/microbiol.2018.1.140

27. Clegg RJ, Kreft J-U. Reducing discrepancies between 3D and 2D simulations due to cell packing density. J Theor Biol. 2017;423: 26–30. doi:10.1016/j.jtbi.2017.04.016

28. Janulevicius A, Van Loosdrecht MC, Simone A, Picioreanu C. Cell Flexibility Affects the Alignment of Model Myxobacteria. Biophys J. 2010;99: 3129–3138.

29. Celler K, Hödl I, Simone A, Battin TJ, Picioreanu C. A mass-spring model unveils the morphogenesis of phototrophic *Diatoma* biofilms. Sci Rep. 2014;4. doi:10.1038/srep03649

30. Storck T, Picioreanu C, Virdis B, Batstone DJ. Variable cell morphology approach for Individual-based Modeling of microbial communities. Biophys J. 2014;106: 2037–2048. doi:10.1016/j.bpj.2014.03.015

31. Ericson C. Real-time collision detection. Amsterdam; Boston: Elsevier; 2005.

32. Purcell EM. Life at low Reynolds number. Am J Phys. 1977;45: 3–11.

33. Berg HC. Random walks in biology. Princeton: Princeton University Press; 1993.

34. Palsson E. A three-dimensional model of cell movement in multicellular systems. Future Gener Comput Syst. 2001;17: 835–852. doi:10.1016/S0167-739X(00)00062-5

35. Ricardo H. A modern introduction to differential equations. Third edition. London [England]; San Diego, CA: Academic Press; 2021.

36. Stevens AB, Hrenya CM. Comparison of soft-sphere models to measurements of collision properties during normal impacts. Powder Technol. 2005;154: 99–109. doi:10.1016/j.powtec.2005.04.033

37. Rittmann BE, McCarty PL. Environmental Biotechnology: Principles and Applications. New York, N.Y.: McGraw-Hill Education; 2018.

38. Heijnen JJ. A new thermodynamically based correlation of chemotropic biomass yields. Antonie Van Leeuwenhoek Int J Gen Mol Microbiol. 1991;60: 235–256.

39. Gogulancea V, González-Cabaleiro R, Li B, Taniguchi D, Jayathilake PG, Chen J, et al. Individual Based Model Links Thermodynamics, Chemical Speciation and Environmental Conditions to Microbial Growth. Front Microbiol. 2019;10. doi:10.3389/fmicb.2019.01871

40. Dyke P. Advanced calculus. London: Macmillan Press, Ltd.; 1998.

41. Picioreanu C, van Loosdrecht MCM, Heijnen JJ. Effect of diffusive and convective substrate transport on biofilm structure formation: A two-dimensional modeling study. Biotechnol Bioeng. 2000;69: 504–515.

42. Gunawardena J. Time-scale separation – Michaelis and Menten’s old idea, still bearing fruit. FEBS J. 2014;281: 473–488. doi:10.1111/febs.12532

43. Rittmann BE, Schwarz AO, Eberl H, Morgenroth E, Peréz J, Van Loosdrecht MCM, et al. Results from the multi-species Benchmark Problem (BM3) using one-dimensional models. Water Sci Technol. 2004;49: 163–168.

44. Noguera DR, Picioreanu C. Results from the multi-species Benchmark Problem 3 (BM3) using two-dimensional models. Water Sci Technol. 2004;49: 169–176.

45. Oyebamiji OK, Wilkinson DJ, Li B, Jayathilake PG, Zuliani P, Curtis TP. Bayesian emulation and calibration of an individual-based model of microbial communities. J Comput Sci. 2019;30: 194–208. doi:10.1016/j.jocs.2018.12.007

46. Kreft J-U. Biofilms promote altruism. Microbiology. 2004;150: 2751–2760. doi:10.1099/mic.0.26829-0

47. Xavier JB, Foster KR. Cooperation and conflict in microbial biofilms. Proc Natl Acad Sci U S A. 2007;104: 876–881. doi:10.1073/pnas.0607651104

48. Boddy L. Saprotrophic cord-forming fungi: warfare strategies and other ecological aspects. Mycol Res. 1993;97: 641–655. doi:10.1016/S0953-7562(09)80141-X

49. Nadell CD, Bucci V, Drescher K, Levin SA, Bassler BL, Xavier JB. Cutting through the complexity of cell collectives. Proc R Soc B. 2013;280: 20122770–20122770. doi:10.1098/rspb.2012.2770

50. Horn H, Lackner S. Modeling of Biofilm Systems: A Review. In: Muffler K, Ulber R, editors. Productive Biofilms. Springer International Publishing; 2014. pp. 53–76. Available: http://link.springer.com/chapter/10.1007/10_2014_275

51. Song H-S, Cannon WR, Beliaev AS, Konopka A. Mathematical Modeling of Microbial Community Dynamics: A Methodological Review. Processes. 2014;2: 711–752. doi:10.3390/pr2040711

52. Esser DS, Leveau JHJ, Meyer KM. Modeling microbial growth and dynamics. Appl Microbiol Biotechnol. 2015; 1–16. doi:10.1007/s00253-015-6877-6

53. Hellweger FL, Clegg RJ, Clark JR, Plugge CM, Kreft J-U. Advancing microbial sciences by individual-based modelling. Nat Rev Microbiol. 2016;14: 461–471. doi:10.1038/nrmicro.2016.62

54. Gorochowski TE. Agent-based modelling in synthetic biology. Pinheiro VB, editor. Essays Biochem. 2016;60: 325–336. doi:10.1042/EBC20160037

55. Mattei MR, Frunzo L, D’Acunto B, Pechaud Y, Pirozzi F, Esposito G. Continuum and discrete approach in modeling biofilm development and structure: a review. J Math Biol. 2017; 1–59. doi:10.1007/s00285-017-1165-y

56. Koshy-Chenthittayil S, Archambault L, Senthilkumar D, Laubenbacher R, Mendes P, Dongari-Bagtzoglou A. Agent Based Models of Polymicrobial Biofilms and the Microbiome—A Review. Microorganisms. 2021;9: 417. doi:10.3390/microorganisms9020417

57. Van Den Berg NI, Machado D, Santos S, Rocha I, Chacón J, Harcombe W, et al. Ecological modelling approaches for predicting emergent properties in microbial communities. Nat Ecol Evol. 2022;6: 855–865. doi:10.1038/s41559-022-01746-7

58. Nagarajan K, Ni C, Lu T. Agent-Based Modeling of Microbial Communities. ACS Synth Biol. 2022;11: 3564–3574. doi:10.1021/acssynbio.2c00411

59. Alpkvist E, Klapper I. A multidimensional multispecies continuum model for heterogeneous biofilm development. Bull Math Biol. 2007;69: 765–789. doi:10.1007/s11538-006-9168-7

60. Rahman KA, Sudarsan R, Eberl HJ. A mixed-culture biofilm model with cross-diffusion. Bull Math Biol. 2015;77: 2086–2124. doi:10.1007/s11538-015-0117-1

61. Angert ER. Alternatives to binary fission in bacteria. Nat Rev Microbiol. 2005;3: 214–224. doi:10.1038/nrmicro1096

62. Schulz HN, Jørgensen BB. Big bacteria. Annu Rev Microbiol. 2001;55: 105–137. doi:10.1146/annurev.micro.55.1.105

63. Young KD. The Selective Value of Bacterial Shape. Microbiol Mol Biol Rev. 2006;70: 660–703. doi:10.1128/MMBR.00001-06

64. Yang DC, Blair KM, Salama NR. Staying in Shape: the Impact of Cell Shape on Bacterial Survival in Diverse Environments. Microbiol Mol Biol Rev. 2016;80: 187–203. doi:10.1128/MMBR.00031-15

65. Winkle JJ, Igoshin OA, Bennett MR, Josić K, Ott W. Modeling mechanical interactions in growing populations of rod-shaped bacteria. Phys Biol. 2017;14: 055001. doi:10.1088/1478-3975/aa7bae

66. Smith WPJ, Davit Y, Osborne JM, Kim W, Foster KR, Pitt-Francis JM. Cell morphology drives spatial patterning in microbial communities. Proc Natl Acad Sci. 2017;114: E280–E286. doi:10.1073/pnas.1613007114

67. Aguilar-Trigueros CA, Boddy L, Rillig MC, Fricker MD. Network traits predict ecological strategies in fungi. ISME Commun. 2022;2: 1–11. doi:10.1038/s43705-021-00085-1

68. Pfeffer C, Larsen S, Song J, Dong M, Besenbacher F, Meyer RL, et al. Filamentous bacteria transport electrons over centimetre distances. Nature. 2012;491: 218–221. doi:10.1038/nature11586

69. Martins AMP, Picioreanu C, Heijnen JJ, van Loosdrecht MCM. Three-dimensional dual-morphotype species Modeling of activated sludge flocs. Environ Sci Technol. 2004;38: 5632– 5641. doi:10.1021/es0496591

70. Ofiţeru ID, Bellucci M, Picioreanu C, Lavric V, Curtis TP. Multi-scale modelling of bioreactor-separator system for wastewater treatment with two-dimensional activated sludge floc dynamics. Water Res. 2014;50: 382–395. doi:10.1016/j.watres.2013.10.053

71. Mitchell JG. The Energetics and Scaling of Search Strategies in Bacteria. Am Nat. 2002;160: 727– 740. doi:10.1086/343874

72. Gorochowski TE, Hauert S, Kreft J-U, Marucci L, Shatil N, Tang T-YD, et al. Toward engineering biosystems with emergent collective functions. Front Bioeng Biotechnol. 2020;8: Article 705. doi:10.3389/fbioe.2020.00705

73. Rudge TJ, Steiner PJ, Phillips A, Haseloff J. Computational modeling of synthetic microbial biofilms. ACS Synth Biol. 2012;1: 345–352. doi:10.1021/sb300031n

74. Matyjaszkiewicz A, Fiore G, Annunziata F, Grierson CS, Savery NJ, Marucci L, et al. BSim 2.0: An Advanced Agent-Based Cell Simulator. ACS Synth Biol. 2017;6: 1969–1972. doi:10.1021/acssynbio.7b00121

75. Gutiérrez M, Gregorio-Godoy P, Pérez del Pulgar G, Muñoz LE, Sáez S, Rodríguez-Patón A. A New Improved and Extended Version of the Multicell Bacterial Simulator gro. ACS Synth Biol. 2017;6: 1496–1508. doi:10.1021/acssynbio.7b00003

76. Kang S, Kahan S, McDermott J, Flann N, Shmulevich I. *Biocellion* : accelerating computer simulation of multicellular biological system models. Bioinformatics. 2014;30: 3101–3108. doi:10.1093/bioinformatics/btu498

77. Naylor J, Fellermann H, Ding Y, Mohammed WK, Jakubovics NS, Mukherjee J, et al. Simbiotics: A Multiscale Integrative Platform for 3D Modeling of Bacterial Populations. ACS Synth Biol. 2017;6: 1194–1210. doi:10.1021/acssynbio.6b00315

78. Bauer E, Zimmermann J, Baldini F, Thiele I, Kaleta C. BacArena: Individual-based metabolic modeling of heterogeneous microbes in complex communities. PLOS Comput Biol. 2017;13: e1005544. doi:10.1371/journal.pcbi.1005544

79. Karimian E, Motamedian E. ACBM: An Integrated Agent and Constraint Based Modeling Framework for Simulation of Microbial Communities. Sci Rep. 2020;10: 8695. doi:10.1038/s41598-020-65659-w

80. Borer B, Ataman M, Hatzimanikatis V, Or D. Modeling metabolic networks of individual bacterial agents in heterogeneous and dynamic soil habitats (IndiMeSH). PLOS Comput Biol. 2019;15: e1007127. doi:10.1371/journal.pcbi.1007127

81. Mirams GR, Arthurs CJ, Bernabeu MO, Bordas R, Cooper J, Corrias A, et al. Chaste: an open source C++ library for computational physiology and biology. PLoS Comput Biol. 2013;9: e1002970. doi:10.1371/journal.pcbi.1002970

82. Ghaffarizadeh A, Heiland R, Friedman SH, Mumenthaler SM, Macklin P. PhysiCell: An open source physics-based cell simulator for 3-D multicellular systems. Poisot T, editor. PLOS Comput Biol. 2018;14: e1005991. doi:10.1371/journal.pcbi.1005991

83. Swat MH, Thomas GL, Belmonte JM, Shirinifard A, Hmeljak D, Glazier JA. Multi-Scale Modeling of Tissues Using CompuCell3D. Methods in Cell Biology. Elsevier; 2012. pp. 325–366. doi:10.1016/B978-0-12-388403-9.00013-8

84. Starruß J, De Back W, Brusch L, Deutsch A. Morpheus: a user-friendly modeling environment for multiscale and multicellular systems biology. Bioinformatics. 2014;30: 1331–1332. doi:10.1093/bioinformatics/btt772

85. Wilensky, U. NetLogo. Center for Connected Learning and Computer-Based Modeling, Northwestern University, Evanston, IL; 1999. Available: http://ccl.northwestern.edu/netlogo/

86. Chin L, Worth D, Greenough C, Coakley S, Holcombe M, Gheorghe M. FLAME-II : a redesign of the flexible large-scale agent-based modelling environment. Rutherford Appleton Lab Tech Rep. 2012.

87. Luke S, Cioffi-Revilla C, Panait L, Sullivan K, Balan G. Mason: A multiagent simulation environment. Simulation. 2005;81: 517–527.

88. North MJ, Tatara E, Collier NT, Ozik J, others. Visual agent-based model development with repast simphony. Tech. rep., Argonne National Laboratory; 2007.

89. Kreft J-U. Mathematical Modeling of Microbial Ecology: Spatial Dynamics of Interactions in Biofilms and Guts. In: Jaykus L-A, Wang HH, Schlesinger LS, editors. Food-Borne Microbes: Shaping the Host Ecosystem. Washington, DC: ASM Press; 2009. pp. 347–377. Available: https://onlinelibrary.wiley.com/doi/abs/10.1128/9781555815479.ch19

90. Hubaux N, Wells G, Morgenroth E. Impact of coexistence of flocs and biofilm on performance of combined nitritation-anammox granular sludge reactors. Water Res. 2015;68: 127–139. doi:10.1016/j.watres.2014.09.036

91. Reichert P. Aquasim: A tool for simulation and data analysis of aquatic systems. Water Sci Technol. 1994;30: 21–30.

92. Wanner O, Reichert P. Mathematical modeling of mixed-culture biofilm. Biotechnol Bioeng. 1996;49: 172–184.

93. Reichert P, Wanner O. Movement of solids in biofilms: significance of liquid phase transport. Water Sci Technol. 1997;36: 321–328.

94. Morgenroth E, Wilderer PA. Influence of detachment mechanisms on competition in biofilms. Water Res. 2000;34: 417–426. doi:10.1016/S0043-1354(99)00157-8

95. Noguera DR, Pizarro GE, Regan JM. Modeling Biofilms. Microbial Biofilms. John Wiley & Sons, Ltd; 2004. pp. 222–249. doi:10.1128/9781555817718.ch13

96. Picioreanu C, Kreft J-U, van Loosdrecht MCM. Particle-based multidimensional multispecies biofilm model. Appl Environ Microbiol. 2004;70: 3024–3040. doi:10.1128/AEM.70.5.3024-3040.2004

97. Clarke M, Maddera L, Harris RL, Silverman PM. F-pili dynamics by live-cell imaging. Proc Natl Acad Sci. 2008;105: 17978–17981. doi:10.1073/pnas.0806786105

